# Opposing Network Patterns of Integration–Segregation in Psychedelic and Sedated States of Consciousness

**DOI:** 10.64898/2026.02.17.706398

**Authors:** Rui Dai, Hyunwoo Jang, Anthony G. Hudetz, Zirui Huang, George A. Mashour

## Abstract

Understanding the relationship of conscious states to large-scale brain dynamics remains a central challenge in neuroscience. Here, diverse pharmacological and physiological perturbations—including psychedelic states, sleep, and propofol sedation—were used to examine integration–segregation organization of brain networks across altered states of consciousness in humans using functional MRI. Across analyses, psychedelic and sedative states exhibited a robust “mirror-image” pattern of integration and segregation. Psychedelics were characterized by increased large-scale integration and reduced segregation of brain network interactions, whereas sleep and propofol sedation showed the opposite configuration. These opposing integration–segregation patterns were consistently indexed by complementary measures of functional connectivity, network topology, and interaction complexity, capturing non-redundant dimensions of large-scale brain organization. Importantly, this mirror-image organization generalized across multiple spatial levels and reliably differentiated conscious states in an unbiased, data-driven manner. Together, these findings demonstrate that psychedelic and sedated states are characterized by systematic and opposing shifts in large-scale integration and segregation, with implications for mechanisms of consciousness.

## Introduction

The investigation of consciousness has become increasingly tractable using neuroimaging techniques such as functional magnetic resonance imaging (fMRI) that allow measurement of large-scale brain dynamics and their relationship to cognitive and conscious states^2,3^. Pharmacological interventions further extend this toolkit by enabling controlled and reversible perturbations of consciousness. Two major classes of psychoactive compounds are particularly useful to advance this approach. Anesthetic agents reliably induce behavioral unresponsiveness and disrupt large-scale cortical communication, providing a well-established model of depressed consciousness^4,5^. Psychedelic compounds, in contrast, alter conscious experience in qualitatively different ways, dramatically altering phenomenology—including intensified perception and altered self-experience—while typically preserving wakefulness^6–8^. However, they are rarely investigated together in systematic studies.

A central open question in the field is whether integration and segregation constitute key determinants of conscious state. Although anesthesia and psychedelics are assumed to produce neural changes in divergent directions, no study has systematically tested whether these shifting neural signatures reflect structured, large-scale transformations along this integration–segregation axis of brain organization. Neuroimaging studies of anesthetic-induced unconsciousness reveal a highly convergent systems-level profile characterized by weakened long-range interactions and constrained neural dynamics. Resting-state fMRI demonstrates robust reductions in communication across higher-order association cortices, with relative preservation of primary sensory systems^9–11^. Network-level analyses further indicate a shift toward more locally clustered and modular configurations, accompanied by reduced efficiency of information exchange across distributed systems^12,13^. Complementing these spatial findings, information-theoretic approaches find a contraction of the brain’s dynamical repertoire, with neural activity becoming increasingly predictable and structurally constrained^14,15^. Together, these results depict anesthesia as a transition toward a constrained and fragmented mode of large-scale brain organization that disrupts the conditions supporting conscious processing.

In contrast, neuroimaging of psychedelic states demonstrates a large-scale reorganization in the opposite direction, marked by enhanced integration, reduced segregation, and enriched neural dynamics. Resting-state fMRI reveals increased global connectivity, reduced within-network connectivity, and strengthened cross-network coupling across associative systems such as the default mode, salience, and frontoparietal networks^6,8,16,17^. Graph-theoretical and cortical gradient analyses show a collapse of modular structure and flattening of the unimodal–transmodal gradient, indicating more globally distributed and flexible modes of cortical communication^18–20^. At the level of neural dynamics, information-theoretic and spectral approaches reveal expanded state repertoires, the emergence of higher-frequency components, and increased signal diversity, indicative of a more flexible and exploratory dynamical regime^15,21–23^. Together, these findings stand in direct contrast to anesthesia-induced network contraction and may underlie the profound alterations of conscious experience elicited by psychedelics. Although derived from diverse analytical approaches, this body of work converges on a common picture in which conscious states are systematically differentiated by shifts in large-scale integration and segregation^24^.

Here, we report a systematic test of whether integration and segregation offer a unifying organizational framework for conscious brain states. To do so, we analyzed fMRI data spanning multiple pharmacological and natural perturbations of consciousness, including psychedelic states (LSD, psilocybin, subanesthetic ketamine, and subanesthetic nitrous oxide), anesthetic-induced unresponsiveness with propofol, and sleep. Across these conditions, we quantified large-scale brain organization at multiple hierarchical levels—from the whole brain, to a unimodal–attentional–transmodal hierarchy, to canonical functional networks—using functional connectivity, network topology, and interaction complexity. This multi-level, multi-metric approach enables a data-driven evaluation of whether shifts in integration and segregation consistently organize diverse conscious states across perturbations, spatial scales, and analytical domains.

## Results

We investigated four classical (i.e., serotonergic) and non-classical psychedelic agents— LSD (n=15), psilocybin (n=7), ketamine (n=12), and nitrous oxide (n=15)—all known to induce non-ordinary states of consciousness ^6,8,20^, as well as sleep and sedation, which are associated with diminished consciousness. For sleep, we analyzed non-REM sleep stages N1 (n=33) and N2 (n=29). N3 sleep was not analyzed due to insufficient sample size (n=3). For sedation, we examined the effects of propofol at various effect-site concentrations: 1.9 μg/ml (n=12), 2.4 μg/ml (n=26), and 2.7 μg/ml (n=27). Each altered state of consciousness was compared to its corresponding baseline wakefulness.

To characterize brain organization across altered states of consciousness, we employed a set of complementary measures that capture distinct aspects of integration and segregation across multiple spatial scales (**Figure 1**). Functional connectivity quantifies the strength of statistical coupling between brain regions and provides a measure of functional association that can be examined across different spatial scales. Graph-theoretical measures of global efficiency characterize the topological conditions that support efficient communication across the network. Interaction complexity captures the diversity and dimensionality of interaction patterns over time, reflecting dynamical segregation rather than static coupling strength. Together, these measures dissociate changes in functional association, network topology, and interaction dynamics, enabling a data-driven assessment of integration–segregation across states of consciousness.

**Figure 1.**
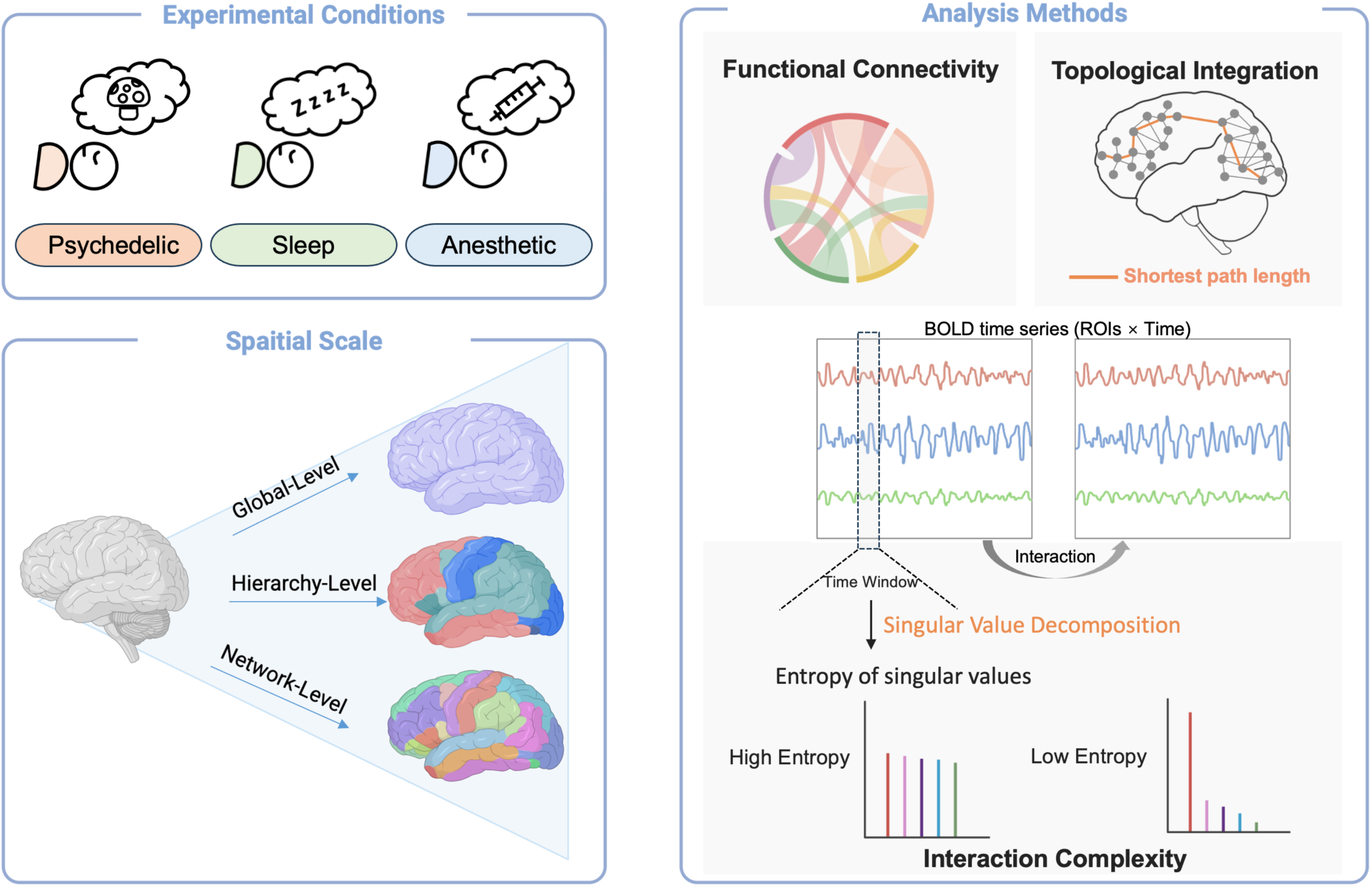
**Schematic overview of the experimental conditions, spatial scales, and analytical framework**. Participants were examined across three experimental conditions: psychedelic, sleep, and anesthetic states (top left). Neural data were analyzed across three spatial scales (bottom left), including a global level (whole-brain average), a hierarchical level capturing unimodal–association–transmodal organization, and a network level based on canonical large-scale functional networks. Three complementary analytical approaches were applied (right). Functional connectivity was quantified from pairwise correlations among regional blood-oxygen level dependent (BOLD) time series. Topological integration was assessed using graph-theoretical metrics derived from shortest path lengths within functional connectivity networks. Interaction complexity was computed from sliding-window BOLD time series matrices (regions of interest × time) using singular value decomposition (SVD), with complexity indexed by the entropy of the resulting singular value spectrum. Together, these methods are used to characterize large-scale brain organization across altered states of consciousness.

At the global level, states of consciousness were robustly differentiated using summary measures that aggregate functional connectivity across the entire brain, without reference to specific network–network interactions. Across all four psychedelic conditions, a consistent pattern of increased large-scale integration was observed relative to normal wakefulness (**Figure 2**, **Table S1-S3**). Psychedelic states were characterized by elevated whole-brain average functional connectivity, increased between-network functional connectivity, and reduced within-network functional connectivity, indicating enhanced coupling across distributed brain networks (**Figure 2a**). Convergent effects were observed in network topology (**Figure 2b**) and interaction complexity (**Figure 2c**): global efficiency was increased, and whole-brain signal complexity was elevated under psychedelic conditions, reflecting both greater topological integration and more diverse interaction patterns over time.

**Figure 2.**
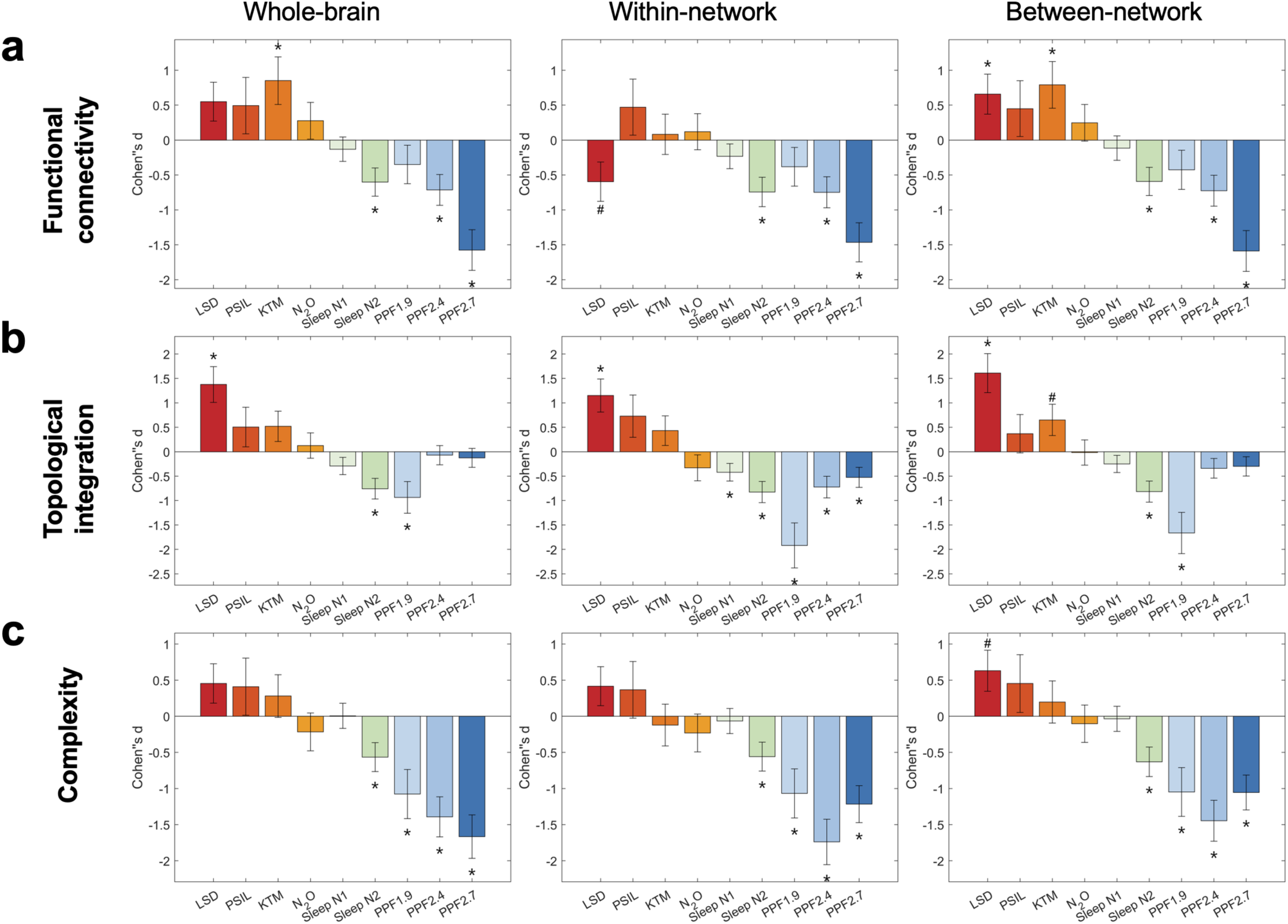
Global-level patterns of functional connectivity, topology, and interaction complexity across conscious states. (a) Changes in global functional connectivity measures, including whole-brain average functional connectivity, between-network functional connectivity, and within-network functional connectivity, shown for psychedelic states, sleep, and propofol anesthesia relative to their corresponding baseline conditions. (b) Changes in global topological integration, quantified using normalized efficiency, shown for the same set of conscious states relative to baseline. (c) Changes in global complexity, quantified from the normalized singular value entropy of functional connectivity patterns within sliding time windows, shown for the same set of conscious states relative to baseline. All values represent group-level effects relative to baseline. Error bars indicate the standard error of Cohen’s d. Statistical significance was assessed using paired two-sided tests, with multiple comparisons controlled using false discovery rate (FDR) correction at α = 0.05. Asterisks (*) denote effects that remain significant after FDR correction (FDR-corrected p < 0.05), whereas hashes (#) indicate effects that are significant prior to correction (uncorrected p < 0.05). Exact effect sizes (Cohen’s d) and FDR-corrected p values are reported in Supplementary Table S1-S3. LSD: lysergic acid diethylamide, PSIL: psilocybin, KTM: ketamine, N2O: nitrous oxide. PPF1.9/2.4/2.7: propofol administered at effect-site concentrations of 1.9, 2.4, and 2.7 μg·mL⁻¹.

In contrast, states associated with diminished consciousness exhibited the opposite configuration across the same measures. Both non-REM sleep and propofol sedation were characterized by reductions in whole-brain functional connectivity, accompanied by decreased between-network and within-network connectivity. These changes were paralleled by reductions in global efficiency and interaction complexity, indicating a breakdown of large-scale integration and a concomitant reduction in the diversity of network interactions.

Together, these results reveal a robust mirror-image pattern of large-scale brain organization across altered states of consciousness: psychedelic states consistently shift brain networks toward greater integration and interactional diversity, whereas non-REM sleep and propofol sedation converge on a state of reduced integration and diminished network dynamics.

At the level of the cortical functional hierarchy, large-scale brain organization exhibited systematic and graded differences across unimodal, attentional, and transmodal systems (**Figure 3**, **Table S4**). We examined interactions within functional systems as well as interactions between hierarchical levels of functional organization, spanning unimodal–attention, unimodal–transmodal, and attention–transmodal relationships, across functional connectivity (**Figure 3a**), topological integration (**Figure 3b**), and interaction complexity (**Figure 3c**).

**Figure 3.**
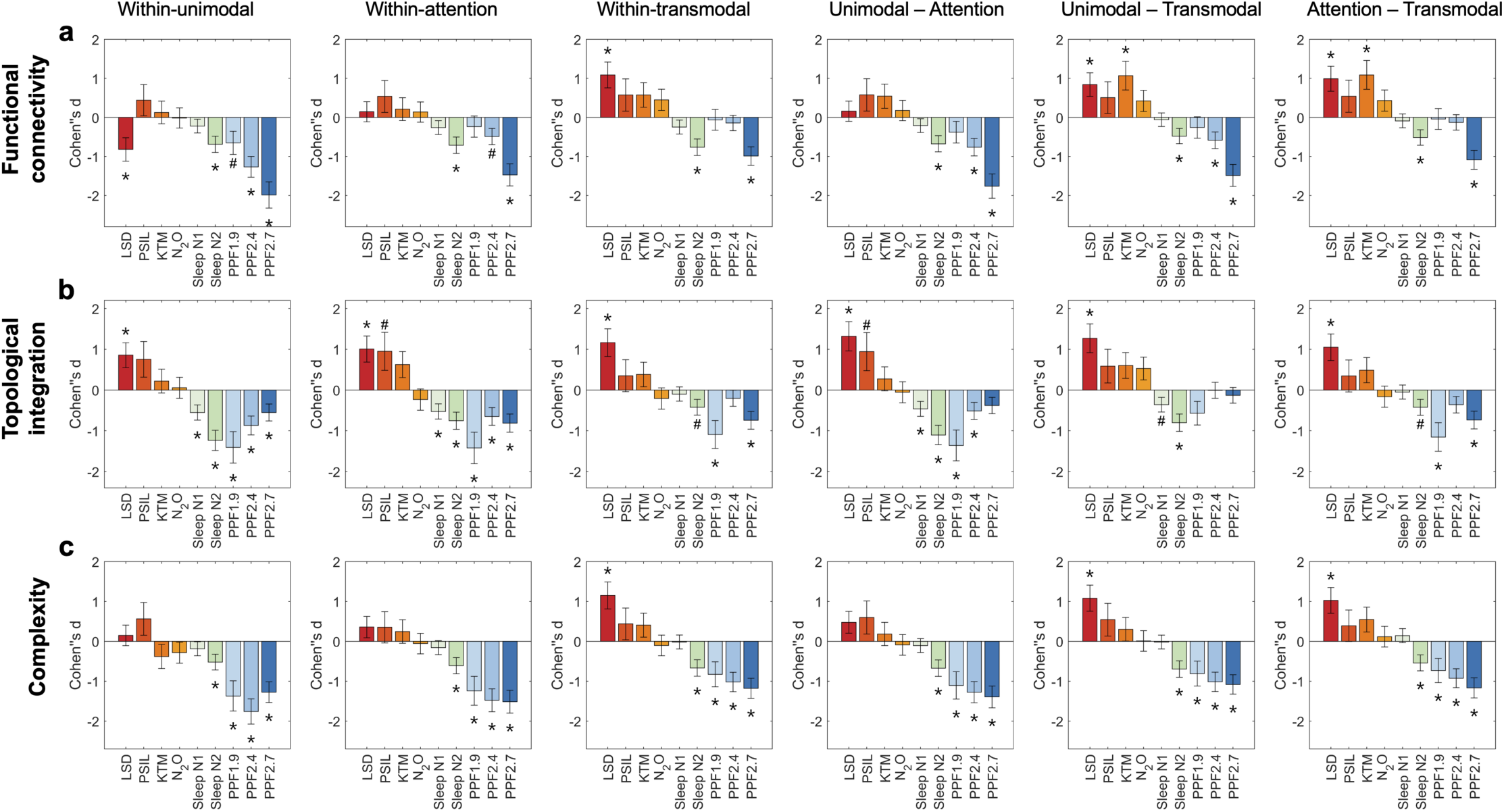
Hierarchical-level patterns of functional connectivity, topology, and interaction complexity across conscious states. (a) Functional connectivity within and between hierarchical levels of cortical organization, including unimodal, attentional, and transmodal systems, shown for psychedelic states, sleep, and propofol sedation relative to their corresponding baseline conditions. (b) Topological integration within and between hierarchical levels, quantified using graph-theoretical global efficiency, shown for the same set of conscious states relative to baseline. (c) Interaction complexity within and between hierarchical levels, quantified from the dimensionality of network–network interaction patterns within sliding time windows, shown for the same set of conscious states relative to baseline. All values represent group-level effects relative to baseline. Error bars indicate the standard error of Cohen’s d. Statistical significance was assessed using paired two-sided tests with false discovery rate (FDR) correction for multiple comparisons. Asterisks (*) denote effects significant after FDR correction (p < 0.05), while hashes (#) indicate effects significant prior to correction (uncorrected p < 0.05). Exact effect sizes and FDR-corrected *p* values are reported in Supplementary Table S4. LSD: lysergic acid diethylamide, PSIL: psilocybin, KTM: ketamine, N2O: nitrous oxide. PPF1.9/2.4/2.7: propofol administered at effect-site concentrations of 1.9, 2.4, and 2.7 μg·mL⁻¹.

Across all three measures, altered states of consciousness exhibited a robust mirror-image pattern across both within-system and between-system interactions along the cortical functional hierarchy. Relative to baseline wakefulness, psychedelic states were associated with increases in hierarchical interactions, whereas non-REM sleep and propofol sedation showed concordant reductions across the same hierarchical contrasts.

Within this overall mirror-image organization, psychedelic states tended to show stronger effects on transmodal-related interactions. In functional connectivity, increases were apparent both within transmodal systems and in interactions linking transmodal regions with attentional and unimodal systems, with a similar pattern observed in interaction complexity.

In contrast, states associated with diminished consciousness showed the largest reductions at lower levels of the cortical hierarchy. Across non-REM sleep and propofol sedation, reductions were most pronounced within unimodal systems, within attentional systems, and in unimodal–attention interactions, a pattern that was consistently observed across functional connectivity, topological integration, and interaction complexity.

Together, these findings demonstrate that alterations in conscious state are differentially expressed across levels of the cortical functional hierarchy. Psychedelic states tend to show stronger effects on higher-order interactions involving attentional and transmodal systems, whereas diminished states of consciousness are characterized by pronounced reductions at lower hierarchical levels, including unimodal systems and unimodal-attention interactions. Across these hierarchical contrasts, interaction complexity provided a particularly sensitive index of state-dependent differentiation.

At the level of functional networks, altered states of consciousness continued to exhibit a robust mirror-image pattern in large-scale brain organization (**Figure 4**, **Table S5**). We examined pairwise interactions among seven functional networks—including visual, somatomotor, dorsal attention, ventral attention, limbic, frontoparietal, and default-mode networks—across functional connectivity, topological integration, and interaction complexity. Across all psychedelic conditions, network-level analyses revealed widespread increases in inter-network functional connectivity, particularly involving frontoparietal and default-mode networks (**Figure 4a**). These increases were accompanied by corresponding increases in topological integration (**Figure 4b**) and interaction complexity (**Figure 4c**). In contrast, sleep and propofol sedation showed the opposite network-level pattern. Both the deeper stage of non-REM sleep and increasing effect-site concentrations of propofol displayed robust reductions in inter-network functional connectivity, together with parallel decreases in topological integration and interaction complexity across a broad range of network pairs (**Figure 4a–c**). The magnitude of these reductions increased systematically with anesthetic concentration. Taken together, network-level analyses revealed consistent, state-dependent differences in functional connectivity, topological integration, and interaction complexity across large-scale brain networks. By resolving these effects at the level of individual functional networks, these findings reveal that the mirror-like reorganization observed at global and hierarchical levels is underpinned by coherent, network-specific patterns of integration and segregation.

**Figure 4.**
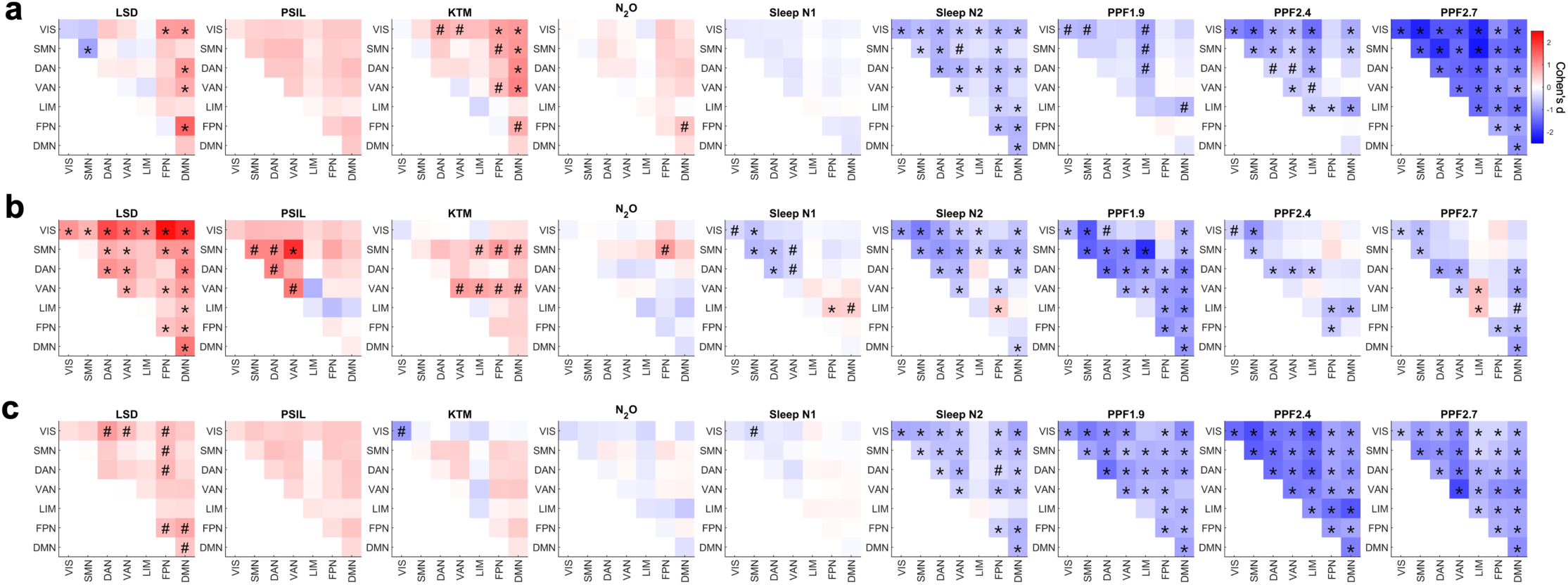
Network-level patterns of functional connectivity, topology, and interaction complexity across conscious states. (a) Inter-network functional connectivity between seven canonical functional networks, shown for psychedelic states, sleep, and propofol sedation relative to their corresponding baseline conditions. (b) Network-level topological integration between functional networks, quantified using graph-theoretical measures based on shortest path length, shown for the same set of conscious states relative to baseline. (c) Network-level interaction complexity between functional networks, quantified from the dimensionality of network–network interaction patterns within sliding time windows, shown for the same set of conscious states relative to baseline. All values represent group-level effects relative to baseline. Error bars indicate the standard error of Cohen’s d. Statistical significance was assessed using paired two-sided tests with false discovery rate (FDR) correction for multiple comparisons. Asterisks (*) denote effects significant after FDR correction (p < 0.05), while hashes (#) indicate effects significant prior to correction (uncorrected p < 0.05). Exact effect sizes and FDR-corrected p values are reported in Supplementary Table S5. LSD: lysergic acid diethylamide, PSIL: psilocybin, KTM: ketamine, N2O: nitrous oxide. PPF1.9/2.4/2.7: propofol administered at effect-site concentrations of 1.9, 2.4, and 2.7 μg·mL⁻¹. VIS: visual network; SMN: somatomotor network; DAN: dorsal attention network; VAN: ventral attention network; LIM: limbic network; FPN: frontoparietal network; DMN: default-mode network.

To assess the consistency of effects across metrics and spatial scales, we next examined whether large-scale network features yielded a common ordering of conscious states. We first pooled the effect size rank of the states by averaging the ranks of Cohen’s d of nine states at each global- and hierarchical-level analysis (**Figure 5a**). This yielded a clear ordering, with psychedelic states occupying higher ranks (i.e., overall positive Cohen’s d) and sleep and propofol sedation occupying progressively lower ranks (i.e., overall negative Cohen’s d). To evaluate which level or metric best aligned with this overall rank, we assessed the Kendall’s tau between the average rank and within-metric rank (**Figure 5b**). Across spatial scales, interaction complexity showed the strongest agreement with the average rank, followed by functional connectivity and topological integration, indicating that complexity-based measures most consistently captured the overall ordering of conscious states. To further evaluate whether this ordering can be reproduced in a network-level approach, we performed an unsupervised principal component analysis (PCA) on network-level Cohen’s d values. Projection onto the first principal component revealed a clear separation of psychedelic states from sleep and propofol sedation along a single dominant axis (**Figure 5c**). Notably, the ordering along this axis perfectly recapitulated the ordering in average rank obtained from global- and hierarchy-level analyses. Examination of the first principal component revealed that the separation was driven by a coordinated contribution of multiple features, including functional connectivity, topological integration, and interaction complexity, rather than by any single metric or network class (**Figure 5d–f**). Together, analyses across multiple metrics and spatial levels consistently revealed mirror-image patterns of integration and segregation across states of consciousness.

**Figure 5.**
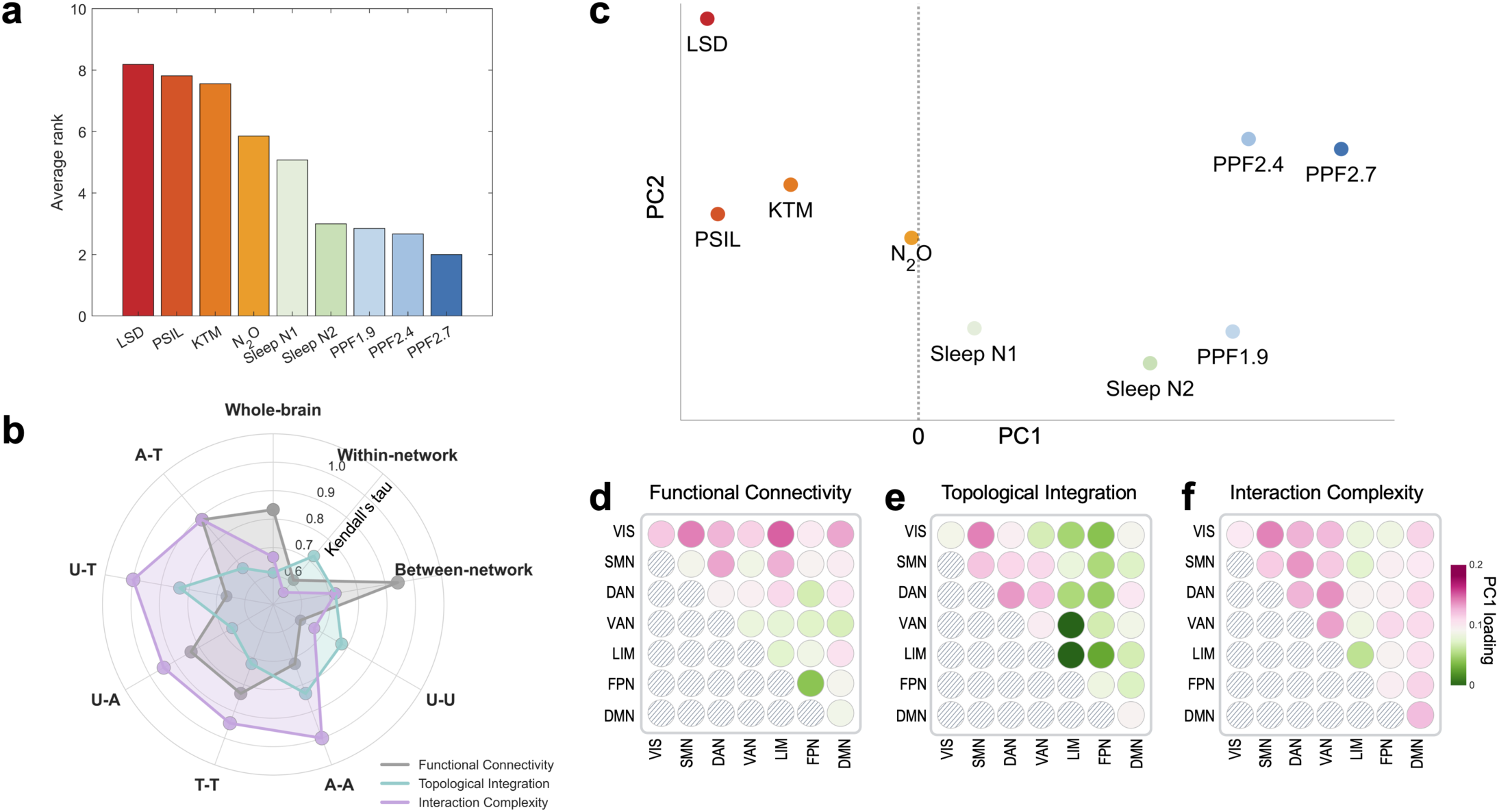
Data-driven consistency of integration–segregation effects across metrics and spatial scales. (a) Average rank of conscious states, obtained by combining effect sizes across all analyses, integrating results from multiple metrics (functional connectivity, topological integration, and interaction complexity) and multiple spatial scales. (b) Agreement between individual rankings and the average rank, quantified using Kendall’s tau, shown separately for each metric and spatial level. (c) Principal component analysis (PCA) applied to network-level feature vectors derived from functional connectivity, topological integration, and interaction complexity measures, showing the projection of all states onto the first two principal components. (d–f) Loadings of the first principal component (PC1) for network-level features derived from (d) functional connectivity, (e) topological integration, and (f) interaction complexity. U-U: unimodal-unimodal; A-A: attention-attention; T-T: transmodal-transmodal; U-A: unimodal-attention; U-T: unimodal-transmodal; A-T: attention-transmodal. LSD: lysergic acid diethylamide, PSIL: psilocybin, KTM: ketamine, N2O: nitrous oxide. PPF1.9/2.4/2.7: propofol administered at effect-site concentrations of 1.9, 2.4, and 2.7 μg·mL⁻¹. VIS: visual network; SMN: somatomotor network; DAN: dorsal attention network; VAN: ventral attention network; LIM: limbic network; FPN: frontoparietal network; DMN: default-mode network.

## Discussion

This study systematically investigated a broad spectrum of altered conscious states, revealing a striking “mirror image” pattern in network features of integration and segregation. Across psychedelic states sleep, and propofol sedation, changes in functional connectivity, network topology, and interaction complexity consistently occurred in opposite directions. Psychedelic states were characterized by increased large-scale interactions across brain networks, whereas sleep and propofol sedation showed convergent reductions across the same metrics. Importantly, this mirror-image pattern was preserved across global, hierarchical, and network-level analyses, indicating that opposing patterns of integration and segregation are a stable feature of altered conscious states rather than a property of any single spatial scale or analytical approach.

While prior research has identified alterations in between-network and within-network integration during psychedelic states^8,22,25,26^, sleep^27,28^, and sedation^4,9,24,29–32^, most studies have examined these effects within a limited set of metrics or spatial scales or studied the states in isolation. Here, we extend this literature by systematically characterizing large-scale brain organization across a broad spectrum of altered states using complementary measures of functional connectivity, network topology, and interaction complexity. By jointly considering these metrics and states, our approach provides an integrated framework for assessing how large-scale patterns of brain interaction reorganize across different modes of consciousness.

Crucially, we examined large-scale brain organization across multiple spatial levels, encompassing global measures of whole-brain coupling, an intermediate cortical functional hierarchy defined by a unimodal–attentional–transmodal axis^33^, and network-specific analyses based on canonical functional systems. Together, these levels provide complementary perspectives on how large-scale brain organization reconfigures across conscious states. This hierarchical level provides an intermediate scale between global measures of large-scale coupling and finer-grained network-specific analyses. The unimodal-attention-transmodal hierarchy captures a graded transition from primary sensory processing to higher-order associative integration and has emerged as a fundamental organizational principle of the human cortex^20,34–38^. By quantifying interactions both within and between unimodal, attentional, and transmodal systems, this approach allows state-dependent reconfigurations to be localized along the functional hierarchy while remaining anchored to global and network-level findings. These findings are consistent with prior psychedelic studies reporting enhanced interactions among higher-order systems^18,39^, and extend them by embedding such effects within a unified hierarchical framework that generalizes across pharmacological and natural perturbations of consciousness.

A key strength of the present study lies in the use of multiple, complementary metrics to characterize large-scale brain organization across altered states of consciousness. Functional connectivity provides a sensitive measure of statistical coupling between brain systems, but on its own cannot distinguish whether increased coupling reflects more efficient integration or simply stronger correlations within a fixed network architecture. Topological integration, derived from graph-theoretical measures, contextualizes connectivity within the broader network structure by characterizing the conditions for efficient large-scale interactions, yet remains relatively insensitive to temporal variability in interaction patterns. Interaction complexity, in contrast, captures the diversity and dimensionality of network interactions over time, but does not directly index connection strength or network efficiency. By combining these metrics, our approach overcomes the limitations inherent to any single measure and provides a more complete characterization of large-scale brain organization. The convergence of functional connectivity, topological integration, and interaction complexity on the same mirror-image pattern across psychedelic states, sleep, and propofol sedation indicates that the observed effects are not driven by metric-specific biases or analytical choices. Instead, these findings reflect a robust reconfiguration of large-scale brain organization that manifests consistently across multiple, non-redundant dimensions. This multi-metric convergence strengthens the interpretability of the results and supports the view that changes in conscious state are accompanied by coordinated shifts in large-scale brain organization, reflecting opposing modes of integration and segregation across multiple, non-redundant dimensions.

Importantly, the mirror-image organization identified here was not specific to a single dataset, compound, or experimental manipulation, but was consistently observed across a diverse set of pharmacological and natural perturbations of consciousness, including multiple psychedelic agents, sleep, and propofol sedation. Despite substantial heterogeneity in study designs, acquisition protocols, and subject populations, all the altered states converged onto the same opposing pattern of large-scale brain organization. Moreover, this distinction did not rely on any predefined ordering of conscious states or theory-driven assumptions. Instead, unsupervised, data-driven analyses of network-level features—most notably principal component analysis—independently recovered a low-dimensional organization that cleanly separated psychedelic states from sleep and anesthetic conditions. The convergence of this data-driven separation with the multi-metric, multi-level results underscores that the observed mirror-image pattern reflects a robust and generalizable organizational principle of large-scale brain dynamics, rather than an artifact of specific metrics, spatial scales, consciousness states, or analytical choices.

This study has limitations. First, it is important to acknowledge that increased connectivity or integration does not always equate to a greater capacity for consciousness. For example, excessive connectivity can lead to pathological states, such as seizures, where neural synchronization is heightened but consciousness is impaired. Second, the reliance on multiple datasets introduces heterogeneity, which may create confounds due to different experimental settings or populations. Finally, although our analyses span multiple spatial scales, they are based on resting-state fMRI and therefore cannot directly address content-specific or task-evoked mechanisms of consciousness. Resting-state approaches are inherently limited in their ability to probe stimulus-driven or perceptual content, and thus our findings should be interpreted as complementary to task-based investigations rather than a complete description of conscious states.

Despite these limitations, the present study makes several substantive contributions. By jointly examining multiple altered states of consciousness across diverse pharmacological and physiological perturbations, we identify a robust mirror-image organization of large-scale brain dynamics that generalizes across compounds, datasets, and experimental contexts. Crucially, this pattern emerges consistently across multiple spatial levels—from global measures to a cortical functional hierarchy and network-specific analyses—and across complementary metrics capturing connection strength, network topology, and interaction diversity. Rather than relying on any single analytical lens or theoretical framework, our multi-level, multi-metric, and data-driven approach reveals opposing modes of large-scale brain organization that distinguish classes of conscious states. This convergence across independent dimensions strengthens the interpretability of the findings and suggests that integration–segregation features constitute a fundamental organizational principle of conscious brain dynamics. By situating diverse altered states within a unified systems-level framework, this work advances a more integrative understanding of consciousness that accommodates heterogeneity across states while identifying shared, generalizable organizational signatures.

## Methods

### Dataset-1: LSD

Fifteen healthy subjects (4 females; mean age ± standard deviation: 30.5 ± 8.0 years) were sourced from the OpenNEURO database^6^. The study design involved two experimental sessions, where participants received either a placebo or 75 μg of intravenous LSD in a counterbalanced order. Subjective experiences were assessed using the 11-Dimensional Altered States of Consciousness (11D-ASC) questionnaire ^40^ at the end of each dosing day. Each participant completed three scans: the first and third were eyes-closed resting-state sessions, while the second involved a music listening task. The first eyes-closed resting-state sessions, under both drug and no-drug (baseline) conditions, were used in this study. Each scan duration was 7 minutes. This study utilized a 3T GE HDx scanner. For echo-planar imaging (EPI), the scan settings were as follows: 35 slices, a repetition time/echo of 2000/35 ms, 3.4 mm slice thickness, a 220 mm field of view, and a 90° flip angle. High-resolution T1 images were acquired as well.

In addition, only preprocessed data were available in the released dataset, followed by several steps. Initially, the first three volumes of each scan were removed to ensure stability. De-spiking was then conducted to correct signal artifacts, followed by slice time correction to align image acquisition timings. Motion correction was applied to counteract participant movement, and brain extraction isolated brain tissue from other elements in the images. The images were aligned to anatomical scans via rigid body registration and further aligned to a 2mm MNI brain template through non-linear registration. The dataset was then scrubbed using a FD threshold of 0.4, with a maximum of 7.1% of volumes scrubbed per scan, replacing them with the mean of surrounding volumes. Further processing included applying a 6mm kernel for spatial smoothing, filtering the data within the 0.01–0.1 Hz range, removing signal drifts through linear and quadratic de-trending, and eliminating motion- and anatomy-related artifacts through regression.

### Dataset-2: Psilocybin

Seven healthy adults (aged 18–45 years) were enrolled in a randomized cross-over functional brain mapping study conducted at Washington University in St. Louis. The dataset was obtained from the OpenNeuro database^25^.

Participants had at least one prior lifetime exposure to a psychedelic substance but no psychedelic use within the six months preceding enrollment. Individuals with a history of psychiatric illness, including depression, psychosis, or substance use disorder, were excluded.

For the present study, analyses focused exclusively on resting-state fMRI data acquired during the first non-drug (baseline) session and the first psilocybin dosing session for each participant. Psilocybin was administered orally at a dose of 25 mg, and resting-state fMRI acquisition commenced approximately 60–180 minutes following ingestion, corresponding to peak drug effects.

Functional MRI data were acquired on a 3T Siemens Prisma scanner at Washington University Medical Center. High-resolution structural images (T1- and T2-weighted) were collected at 0.9 mm isotropic resolution. Functional MRI data were acquired using a multiband, multi-echo echo-planar imaging sequence with the following parameters: voxel size = 2 mm isotropic, repetition time (TR) = 1,761 ms, flip angle = 68°, multiband factor = 6, five echo times (TEs = 14.20, 38.93, 63.66, 88.39, 113.12 ms), and in-plane acceleration factor = 2. Each resting-state scan comprised 510 volumes (15 min 49 s). Two resting-state scans were collected per session. During resting-state acquisition, participants were instructed to fixate on a centrally presented white cross on a black background.

Head motion was monitored in real time using Framewise Integrated Real-time MRI Monitoring (FIRMM), and eye tracking was used to monitor alertness during scanning.

### Dataset-3: Ketamine

The research was approved by the Institutional Review Board of Huashan Hospital, Fudan University, and all participants provided written informed consent ^8,41^. Twelve right-handed individuals (5 females; mean age ± standard deviation: 41.4 ± 8.6 years) were enrolled. Participants had no history of neurological disorders, significant organ dysfunction, or neuropsychiatric medication use, and were classified as American Society of Anesthesiologists physical status I or II. Intravenous ketamine was administered while fMRI scans were performed without interruption throughout the experiment, which spanned 44 to 62 minutes. A 10-minute baseline recording of the conscious state was conducted at the start, with the exception of two participants who had shorter baseline durations of 6 and 11 minutes. Ketamine was then administered at a rate of 0.05 mg/kg per minute over 10 minutes (total dose: 0.5 mg/kg), followed by an increased rate of 0.1 mg/kg per minute for another 10 minutes (cumulative dose: 1.0 mg/kg). Two participants only received the second dose. Once the ketamine infusion was complete, participants naturally regained consciousness. Behavioral responsiveness was monitored every 30 seconds during the scanning session using a button-press task. Our analysis focused on the 10-minute subanesthetic ketamine administration, which is associated with psychedelic experiences. This study utilized a 3T Siemens MAGNETOM scanner. For EPI, the scan settings were as follows: 33 slices, a repetition time/echo time of 2000/30 ms, 5mm slice thickness, a 210 mm field of view, a 64 × 64 image matrix, and a 90° flip angle. High-resolution T1 images were acquired as well.

### Dataset-4: Nitrous oxide

Eighteen healthy participants (9 females, mean age ± standard deviation: 24.9 ± 3.5 years) were included for this study, approved by the Institutional Review Board of the University of Michigan Medical School (HUM00096321), with all participants giving written informed consent ^8,20^. All participants were classified as American Society of Anesthesiologists physical status I, exclusion criteria included any history of drug abuse, psychosis, and other medical conditions as detailed in the trial registry (https://www.clinicaltrials.gov/ct2/show/NCT03435055). Two participants were excluded due to excessive head motion (affecting 50% of their fMRI data) and one due to incomplete scanning, leaving a final sample of 15 healthy subjects.

This study utilized a within-subjects design with two conditions: a baseline condition and a condition with the administration of subanesthetic nitrous oxide (35%). Each condition included a 6-minute resting-state scan and a 3-minute passive viewing scan. Data from these two scans, acquired both before and during nitrous oxide administration, were used for this study. Neuroimaging data were acquired using a 3T Philips Achieva scanner. For EPI, the scan settings were as follows: 48 slices, a repetition time/echo time of 2000/30ms, 3 mm slice thickness, a 200 mm field of view, and a 90° flip angle. High-resolution T1 images were acquired as well. Subjective experiences were assessed both prior to the scanning session, using an 11D-ASC questionnaire ^40^, and after the scanning session to specifically evaluate experiences during the nitrous oxide administration.

### Dataset-5: Sleep

Thirty-three healthy participants (16 females, mean age ± standard deviation: 22.1 ± 3.2 years) were sourced from the OpenNEURO database ^42,43^, with informed consent obtained from all participants. The dataset includes three non-REM sleep stages (N1, N2, and N3), in addition to an awake resting-state condition. These stages were identified using electroencephalogram signatures analyzed by a registered polysomnographic technologist. This study utilized a 3T Siemens Prisma scanner. For EPI, the scan settings were as follows: 35 slices, a repetition time/echo time of 2100/25 ms, 4 mm slice thickness, a 240 mm field of view, and a 90° flip angle. High-resolution T1 images were acquired as well. Only awake (n=33), N1 (n=33) and N2 (n=29) were included in the analysis due to the limited number of subjects in N3 (n=3). REM sleep was not available in the publicly available dataset.

### Dataset-6: Propofol Sedation (1.9 μg/mL Doses)

Fifteen healthy participants (6 females, mean age ± standard deviation: 26.7 ± 4.8 years) were included for this study, approved by the Institutional Review Board of the Medical College of Wisconsin ^4,44^. All participants provided written informed consent, were classified as American Society of Anesthesiologists physical status I or II, and were scheduled for elective surgery to remove pituitary microadenomas. One participant was excluded due to excessive movement, leaving 14 participants for analysis.

Behavioral responsiveness was measured using the Observer’s Assessment of Alertness/Sedation (OAAS) scale. During baseline and recovery, participants were fully responsive to verbal cues, indicated by an OAAS score of 5. In the light sedation phase, participants responded lethargically to verbal commands, corresponding to an OAAS score of 4, while deep sedation was marked by the absence of a response, with OAAS scores ranging from 1 to 2. Individual propofol target plasma concentrations varied (light sedation: 0.98 ± 0.18 μg/mL; deep sedation: 1.88 ± 0.24 μg/mL), reflecting personal differences in sensitivity to the anesthetic. Propofol infusion rates were manually adjusted using STANPUMP to maintain steady sedation, balancing drug accumulation and elimination. Throughout the study, participants were monitored according to ASA standards, including electrocardiogram, blood pressure, pulse oximetry, and end-tidal CO2, with supplemental oxygen provided via nasal cannula. Resting-state data were collected across four 15-minute scans, each representing a different condition: baseline consciousness, light sedation, deep sedation, and recovery. Data from baseline consciousness, light sedation, and deep sedation conditions were used. This study utilized a 3T GE Signa 750 scanner. For EPI, the scan settings were as follows: 41 slices, a repetition time/echo time of 2000/25 ms, 3.5 mm slice thickness, a 224 mm field of view, and a 77° flip angle. High-resolution T1 images were acquired as well.

### Dataset-7: Propofol Sedation (2.4 μg/mL Doses)

Twenty-six healthy participants (13 females, mean age ± standard deviation: 25.0 ± 4.1 years) were included for this study, which was approved by the Institutional Review Board of the University of Michigan, and all participants provided written informed consent ^37,45,46^. All participants were classified as American Society of Anesthesiologists physical status I, exclusion criteria included any history of drug abuse, psychosis, and other medical conditions.

Before the study, participants fasted for eight hours. On the experiment day, a preoperative evaluation, including a physical exam, was conducted by an anesthesiologist. Two anesthesiologists continuously monitored vital signs, including breathing, heart rate, end-tidal CO_2_, pulse oximetry, and ECG. Noninvasive arterial pressure was recorded using an MR-compatible monitor. Participants received 0.5 mL of 1% lidocaine for local anesthesia before intravenous cannula insertion, and oxygen was delivered at 2 L/min through a nasal cannula. Propofol, chosen for its minimal impact on cerebral blood flow and precise titration capability, was administered via target-controlled bolus and infusion, based on the Marsh pharmacokinetic model using STANPUMP (http://opentci.org/code/stanpump). Dosages increased in 0.4 μg/mL increments until participants showed no behavioral response, with the target concentration maintained for an average of 21.6 ± 10.2 minutes before recovery.

Behavioral responsiveness was assessed by a rubber ball squeeze task, with responses quantified using the BIOPAC MP160 system and AcqKnowledge software. Sixty motor response trials, spaced 90 seconds apart, were conducted during scanning sessions. Between trials, participants engaged in mental imagery tasks such as imagining playing tennis or navigating a space. Further experimental details are available in prior publications ^37,45,46^. Four fMRI sessions were conducted as part of the protocol: a 15-minute conscious baseline, a 30-minute session during and post-propofol infusion, followed by a 15-minute recovery baseline. Data from conscious baseline and deep sedation periods were used in this study. This study utilized a 3T Philips scanner. For EPI, the scan settings were as follows: 28 slices, a repetition time/echo time of 800/25 ms (MB factor of 4), 4 mm slice thickness, a 220 mm field of view, and a 76°flip angle. High-resolution T1 images were acquired as well.

### Dataset-8: Propofol Sedation (2.7 μg/mL Doses)

Thirty healthy participants (20 females, mean age ± standard deviation: 24.4 ± 5.2 years) with complete scan data were included in this study, which was approved by the Institutional Review Board of the University of Michigan, and all participants provided written informed consent^38^. All participants were classified as American Society of Anesthesiologists physical status I, exclusion criteria included any history of drug abuse, psychosis, and other medical conditions. Three participants were excluded due to excessive movement and MRI technical issues, leaving 27 participants for analysis.

The anesthetic procedure was similar to Dataset-6, with propofol manually adjusted to achieve effect-site concentrations of 1.5, 2.0, 2.5, and 3.0 μg/mL. Each level was held for 4 minutes to titrate the dosage and determine the threshold for loss of responsiveness (LOR). To minimize head motion artifacts, the concentration was maintained one step higher than the LOR threshold for about 32 minutes (e.g., if LOR occurred at 2.0 μg/mL, 2.5 μg/mL was maintained). In rare cases, if participants remained responsive at 3.0 μg/mL, the concentration was raised to a maximum of 4.0 μg/mL. The infusion was then stopped, and participants engaged in behavioral tasks, rest, or listened to music.

Eight fMRI scans, each lasting 16 minutes, were conducted over a 2.5-hour session. These included baseline scans (Rest1 and Music1), LOR scans (Rest2 and Music2), and recovery scans (Rest3 and Music3). Between each scan, participants had 1–5 minute breaks. Resting-state scans required participants to lie still with eyes closed, while music-listening involved tracks from Jazz, Rock, Pop, and Country genres, played in random order. During behavioral testing, participants were prompted to squeeze a rubber ball every 10 seconds for 96 cycles, following an audio cue delivered through headphones. Grip strength was measured using the BIOPAC MP160 system. Behavioral transitions during propofol administration were identified by missed and completed squeezes, marking the onset and recovery of responsiveness. Data from conscious baseline and deep sedation periods were used in this study. This study utilized a 3T Philips scanner. For EPI, the scan settings were as follows: 40 slices, a repetition time/echo time of 1400/30 ms (MB factor of 4), 2.9 mm slice thickness, a 220 mm field of view, and a 76° flip angle. High-resolution T1 images were acquired as well.

### fMRI data preprocessing

For the preprocessing of fMRI data in this study (except for dataset-1 and dataset-2), we utilized the AFNI software. The procedure encompassed several steps: First, the initial two frames of each scan were removed to ensure signal stability. This was followed by slice-timing correction to adjust for temporal differences in the acquisition of slices. Second, head motion correction and realignment were performed. Head motion was assessed using frame-wise displacement (FD), calculated as the Euclidean Norm of the six motion parameters. Frames where the FD exceeded 0.8mm, along with the preceding frame, were excluded from the analysis. Third, T1 anatomical images were coregistered for precise alignment, followed by spatial normalization into Talairach space ^47^ and resampling to 3 mm isotropic voxels to standardize image coordinates. Fourth, time-censored data underwent band-pass filtering between 0.01–0.1Hz using AFNI’s 3dTproject. Simultaneously, linear regression was applied to eliminate unwanted components such as linear and nonlinear drift, head motion time series and its derivative, as well as mean time series from white matter and cerebrospinal fluid. Fifth, spatial smoothing (6mm Gaussian kernel) was performed. Finally, each voxel’s time series was normalized to zero mean and unit variance to ensure data consistency.

### Multi-level Functional Connectivity Analysis

Functional connectivity (FC) was quantified as the temporal correlation between regional BOLD time series. Using a functional connectivity–based parcellation scheme^48,49^, time series were extracted from 400 cortical regions of interest (ROIs), and Pearson correlation coefficients were computed between all pairs of ROIs to generate a 400 × 400 functional connectivity matrix for each participant and condition.

At the global level, whole-brain average functional connectivity was computed as the mean of all unique pairwise correlation coefficients across the full ROI-to-ROI matrix, excluding self-connections. In addition, global within-network and between-network functional connectivity were calculated by averaging correlations among ROIs belonging to the same functional network and between ROIs belonging to different networks, respectively, providing complementary indices of functional segregation and integration.

At the functional hierarchy-level, brain regions were grouped into unimodal, attention, and transmodal systems based on canonical network affiliation^49^. Unimodal regions comprised visual and somatomotor networks, attention regions comprised dorsal and ventral attention networks, and transmodal regions comprised default mode and frontoparietal control networks. Within-hierarchy functional connectivity was calculated by averaging pairwise correlations among ROIs belonging to the same hierarchical category, whereas between-hierarchy functional connectivity was calculated by averaging correlations between ROIs belonging to different hierarchical categories.

At the network level, network-specific functional connectivity was examined by averaging ROI-to-ROI correlations within and between the seven predefined functional networks (visual, somatomotor, dorsal attention, ventral attention, limbic, frontoparietal, and default mode^49^). This yielded a 7 × 7 network-level functional connectivity matrix for each participant and condition, in which diagonal elements represent within-network connectivity and off-diagonal elements represent connectivity between specific network pairs. These network-level connectivity estimates were used to characterize how global and hierarchical connectivity changes are distributed across individual network–network interactions.

### Topological Integration Analysis

Topological integration was quantified using normalized weighted global efficiency. For a weighted network, global efficiency is defined as the average inverse shortest path lengths between all node pairs:

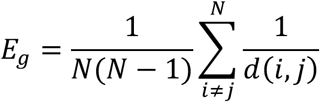

where *N* is the number of nodes and *d*(*i*,*j*) denotes the shortest path length between nodes *i* and *j*. Shortest paths were computed using the Flyd-Washall algorithm (*distance_wei_floyd* function in Brain Connectivity Toolbox^50^) with a logarithmic weight-to-distance transformation, where stronger function connections correspond to shorter distance.

To control for average connectivity changes, all efficiency results reported in the main text were normalized with the corresponding value from a weight-distribution-preserving random null model generated with *randomize_matrix* function^51^):

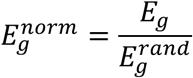

To ensure consistency with the multi-scale functional connectivity analyses described above, topological integration was evaluated across three spatial levels: global, functional hierarchy, and network level. Whole-brain efficiency was computed using all nodes and non-negative edges in the functional connectivity matrix. Within-network or within-hierarchy efficiency was calculated by first generating a smaller functional connectivity matrix composed of nodes within certain functional network or hierarchy, accompanied by the steps described above. Between-network or between-hierarchy efficiency was calculated by (1) constructing a smaller functional connectivity matrix composed of nodes belonging to two networks/hierarchies of interest, (2) obtaining the distance matrix, (3) setting values of within-network or within-hierarchy pairs as NaN, and (4) calculating the average inverse of the remaining values. Random null model generation was done before distance matrix calculation. Global-level within-network and between-network analyses were done by averaging seven within-network and 21 between-network efficiency values, respectively, each obtained by the process described above.

### Interaction Complexity Analysis

Interaction complexity was used to characterize the effective dimensionality of dynamic interactions between brain regions, independent of static connectivity strength. For each participant and condition, regional time series were segmented into non-overlapping sliding windows of 5 TRs. Within each window, a matrix *M* representing the interaction of two sets of nodes was constructed by computing the cross-product of the windowed time series *X*_1_ and *X*_2_(time × region):

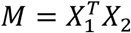

This yields an interaction matrix capturing covariance-like pairwise interactions between regions of two subsets within a short time window. *M* can be rectangular if the numbers of nodes in set 1 and 2 differ. Self-interaction can also be calculated when set 1 = set 2, yielding a symmetric square matrix. The interaction matrix was then mean-centered by subtracting its global mean.

To quantify the complexity of the rectangular matrix *M*, singular value decomposition (SVD) was applied to each mean-centered interaction matrix:

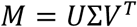

where Σ is a diagonal matrix containing the singular values *s_k_*, which captures the strength of orthogonal interaction modes.

The effective dimensionality of network interactions was quantified as the normalized entropy of the singular value spectrum:

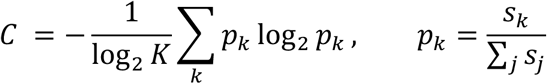

where *K* is the number of singular values. Thus, lower *C* indicates dominance of certain interaction modes (i.e., low-dimensional structure) while higher *C* indicates a more uniform contribution of all modes.

This procedure yielded a time-resolved complexity estimate *C* for each sliding window. Complexity values were then averaged across windows to obtain a single interaction complexity estimate per participant and condition. Identical windowing parameters and normalization procedures were applied across datasets and experimental conditions.

To ensure consistency with the multi-level framework applied to functional connectivity and topological integration, interaction complexity was evaluated across three spatial levels: global, functional hierarchy, and network-level. Whole-brain complexity was computed by using whole ROIs as set 1 and set 2, yielding a 400 × 400 *M*. Within-network and within-hierarchy complexity was calculated by using nodes within a given functional network or hierarchy as set 1 and set 2, yielding a smaller symmetric *M*. Between-network or between-hierarchy efficiency was calculated by choosing two networks or hierarchies of interest as set 1 and set 2, yielding a rectangular *M*. Global-level within-network and between-network complexity was assessed by averaging seven within-network and 21 between-network complexity values, respectively.

### Composite rank of conscious states across analyses

For each analytical metric (functional connectivity, topological integration, and interaction complexity), effect sizes (Cohen’s d) of all conscious states relative to their corresponding baseline conditions were computed at the global and functional hierarchy levels, yielding nine analyses (three global and six hierarchical) per metric. For each analysis-metric pair, nine states were ranked according to the effect size value (not the magnitude or absolute Cohen’s d), with higher ranks generally indicating positive effects, while lower ranks often indicate negative effects. To obtain an overall, data-driven ordering of conscious states, a total of 27 rankings were averaged across analysis-metric pairs.

### Agreement between individual and average rankings

To quantify the consistency between individual rankings and the composite ranking, we assessed rank agreement using Kendall’s tau. For 27 analysis-metric pairs, Kendall’s tau with composite ranking was computed, providing a measure of concordance that captures the extent to which each metric and spatial level aligns with the analysis-general ordering of conscious states.

### Principal component analysis of network-level effects

We performed principal component analysis (PCA) on network-level effect size values. For each conscious state, its feature vector was constructed by concatenating the upper triangular entries of a 7 × 7 matrix from three metrics, yielding 84 unique features of nine states.

PCA was then applied to this 84 × 9 matrix. The matrix was not centered because zero conveys a meaningful context (i.e., no effect). The first two principal components were used for visualization and interpretation, capturing the dominant axes of variance across states.

### Loadings of principal components

To interpret the contributions of individual network-level features to the dominant axis of variance, we examined the loadings of the first principal component (PC1). PC1 loadings were reorganized and visualized in the form of three upper triangular matrices, revealing which network interaction and metric contributed more strongly to discriminating among conscious states.

### Statistics and Reproducibility

All statistical analyses were performed to compare each altered state of consciousness (psychedelic, sleep, and sedation) with its corresponding baseline condition using paired two-sided *t*-tests. Analyses were conducted separately for each analytical metric—functional connectivity, topological integration, and interaction complexity—and at each spatial scale, including the global level, functional hierarchy–level, and network level, as described above.

Because baseline measures can differ across datasets due to variations in experimental protocols, acquisition parameters, and preprocessing pipelines, all statistical comparisons were performed relative to each participant’s own baseline within each dataset. This within-subject design minimized the influence of inter-dataset variability and ensured that observed effects reflected condition-specific changes rather than baseline differences across studies.

To facilitate quantitative comparison across metrics, spatial scales, and conscious states, effect sizes were computed for all analyses using Cohen’s d for paired samples. Effect sizes were used as the primary summary statistic for cross-condition comparisons and subsequent integrative analyses. Statistical significance was assessed using false discovery rate (FDR) correction to control for multiple comparisons, with significance defined as an FDR-adjusted p < 0.05. All statistical tests were two-sided to allow for unbiased detection of effects in either direction.

## Data availability

Data used in the analyses are publicly available from Zenodo repository (https://doi.org/10.5281/zenodo.14029241). The natural sleep fMRI dataset is available from OpenNEURO (https://openneuro.org/datasets/ds003768/versions/1.0.11). The LSD dataset is available from Openneuro (doi: 10.18112/openneuro.ds003059.v1.0.0). The psilocybin dataset is available from Openneuro (openneuro.org/datasets/ds006072).

## Code availability

Publicly available software and toolboxes used for analysis and visualization include AFNI (https://afni.nimh.nih.gov), Brain Connectivity toolbox (https://doi.org/10.1016/j.neuroimage.2009.10.003), JASP (v0.16.3; https://jasp-stats.org), MATLAB R2022a (https://www.mathworks.com), Prism (version 9.5.0; https://www.graphpad.com), and PyCharm (v2023.2.1; https://www.jetbrains.com/pycharm). Custom-built code is available in the Zenodo repository (https://doi.org/10.5281/zenodo.14029241).

### Acknowledgements

This work was funded by National Institutes of Health (Bethesda, Maryland, USA) grants R01-GM103894 (to A.G.H. and Z.H.), R01-GM111293 (to G.A.M.) and T32-GM103730 (to G.A.M., PI, and R.D., Z.H., Fellows).

## Authorship contribution statement

Conceptualization: R.D., H.J., Z.H., G.A.M. Methodology: R.D., H.J., Z.H., G.A.M. Investigation: R.D., H.J., Z.H., G.A.M. Data analysis and visualization: R.D., H.J., Z.H., Supervision: G.A.M, A.G.H., Writing—original draft: R.D., Writing—review & editing: All authors.

---

**Supplementary Table S1.**
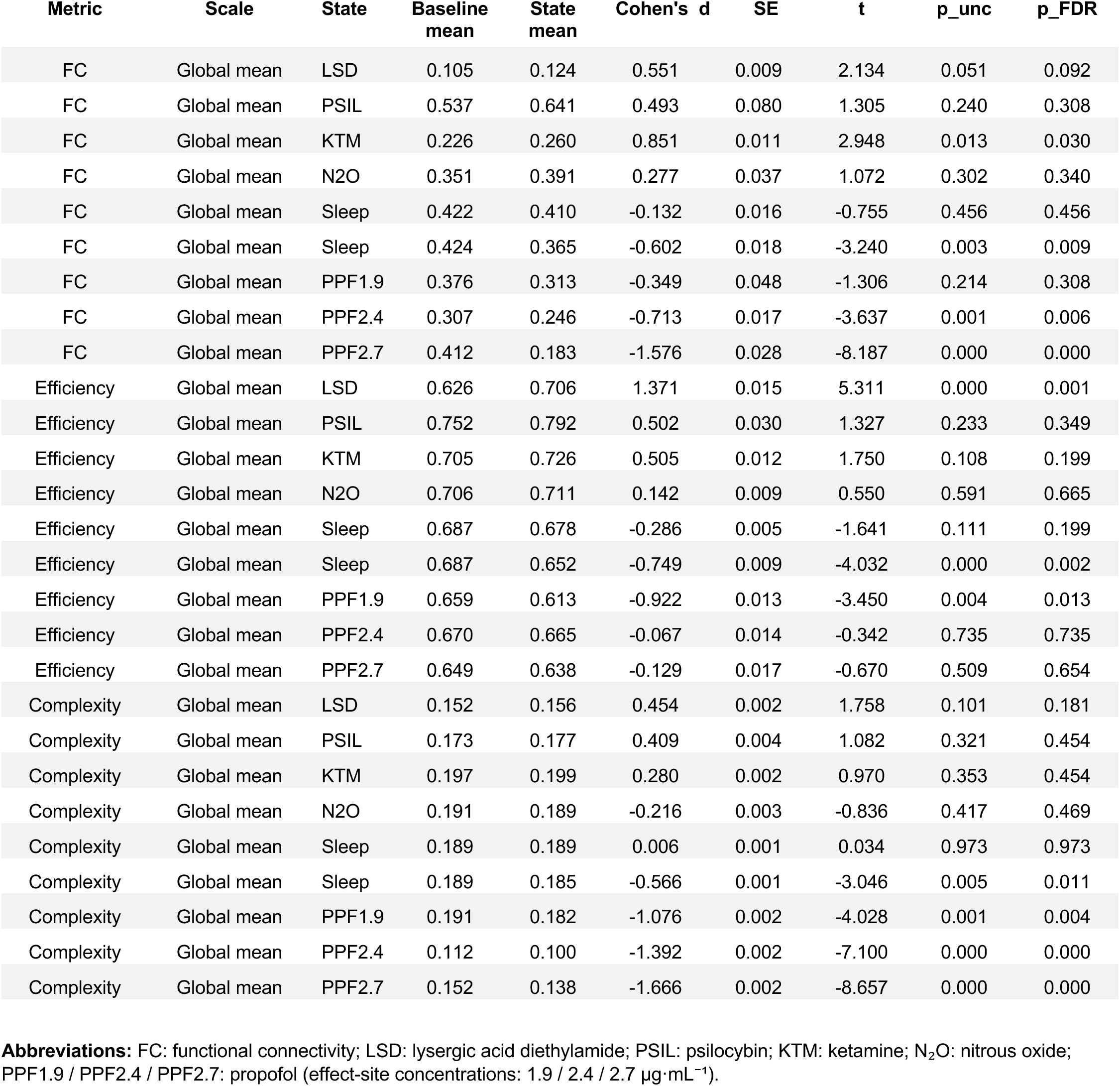
Global within-network results.

**Supplementary Table S2.** Global between-network results.

| Metric | Scale | State | Baseline mean | State mean | Cohen's d | SE | t | p_unc | p_FDR |
| --- | --- | --- | --- | --- | --- | --- | --- | --- | --- |
| FC | Global between-network mean | LSD | 0.067 | 0.090 | 0.658 | 0.009 | 2.547 | 0.023 | 0.042 |
| FC | Global between-network mean | PSIL | 0.518 | 0.618 | 0.450 | 0.084 | 1.190 | 0.279 | 0.359 |
| FC | Global between-network mean | KTM | 0.201 | 0.234 | 0.791 | 0.012 | 2.740 | 0.019 | 0.042 |
| FC | Global between-network mean | N2O | 0.314 | 0.349 | 0.247 | 0.037 | 0.957 | 0.355 | 0.399 |
| FC | Global between-network mean | Sleep N1 | 0.388 | 0.377 | -0.116 | 0.016 | -0.664 | 0.511 | 0.511 |
| FC | Global between-network mean | Sleep N2 | 0.389 | 0.332 | -0.593 | 0.018 | -3.192 | 0.003 | 0.010 |
| FC | Global between-network mean | PPF1.9 | 0.346 | 0.273 | -0.426 | 0.046 | -1.595 | 0.135 | 0.202 |
| FC | Global between-network mean | PPF2.4 | 0.282 | 0.221 | -0.725 | 0.017 | -3.699 | 0.001 | 0.005 |
| FC | Global between-network mean | PPF2.7 | 0.372 | 0.150 | -1.588 | 0.027 | -8.251 | 0.000 | 0.000 |
| Efficiency | Global between-network mean | LSD | 0.649 | 0.695 | 1.581 | 0.007 | 6.124 | 0.000 | 0.000 |
| Efficiency | Global between-network mean | PSIL | 0.754 | 0.782 | 0.362 | 0.029 | 0.959 | 0.375 | 0.422 |
| Efficiency | Global between-network mean | KTM | 0.706 | 0.723 | 0.633 | 0.008 | 2.193 | 0.051 | 0.114 |
| Efficiency | Global between-network mean | N2O | 0.709 | 0.708 | -0.020 | 0.006 | -0.078 | 0.939 | 0.939 |
| Efficiency | Global between-network mean | Sleep N1 | 0.697 | 0.692 | -0.270 | 0.003 | -1.552 | 0.130 | 0.171 |
| Efficiency | Global between-network mean | Sleep N2 | 0.696 | 0.672 | -0.813 | 0.005 | -4.376 | 0.000 | 0.000 |
| Efficiency | Global between-network mean | PPF1.9 | 0.663 | 0.610 | -1.666 | 0.008 | -6.232 | 0.000 | 0.000 |
| Efficiency | Global between-network mean | PPF2.4 | 0.699 | 0.684 | -0.326 | 0.009 | -1.664 | 0.109 | 0.171 |
| Efficiency | Global between-network mean | PPF2.7 | 0.673 | 0.651 | -0.298 | 0.014 | -1.550 | 0.133 | 0.171 |
| Complexity | Global between-network mean | LSD | 0.244 | 0.249 | 0.630 | 0.002 | 2.438 | 0.029 | 0.052 |
| Complexity | Global between-network mean | PSIL | 0.269 | 0.275 | 0.454 | 0.005 | 1.200 | 0.275 | 0.413 |
| Complexity | Global between-network mean | KTM | 0.305 | 0.306 | 0.198 | 0.003 | 0.684 | 0.508 | 0.653 |
| Complexity | Global between-network mean | N2O | 0.288 | 0.286 | -0.103 | 0.004 | -0.398 | 0.697 | 0.784 |
| Complexity | Global between-network mean | Sleep N1 | 0.286 | 0.286 | -0.035 | 0.001 | -0.202 | 0.841 | 0.841 |
| Complexity | Global between-network mean | Sleep N2 | 0.286 | 0.280 | -0.630 | 0.002 | -3.392 | 0.002 | 0.005 |
| Complexity | Global between-network mean | PPF1.9 | 0.292 | 0.279 | -1.049 | 0.003 | -3.923 | 0.002 | 0.005 |
| Complexity | Global between-network mean | PPF2.4 | 0.174 | 0.162 | -1.446 | 0.002 | -7.374 | 0.000 | 0.000 |
| Complexity | Global between-network mean | PPF2.7 | 0.230 | 0.220 | -1.055 | 0.002 | -5.482 | 0.000 | 0.000 |

**Abbreviations:** FC: functional connectivity; LSD: lysergic acid diethylamide; PSIL: psilocybin; KTM: ketamine; N<sub>2</sub>O: nitrous oxide; PPF1.9 / PPF2.4 / PPF2.7: propofol (effect-site concentrations: 1.9 / 2.4 / 2.7 µg·mL<sup>-1</sup>).

**Supplementary Table S3.** Global within-network results.

| Metric | Scale | State | Baseline mean | State mean | Cohen's d | SE | t | p_unc | p_FDR |
| --- | --- | --- | --- | --- | --- | --- | --- | --- | --- |
| FC | Global within-network mean | LSD | 0.323 | 0.304 | -0.597 | 0.008 | -2.311 | 0.037 | 0.082 |
| FC | Global within-network mean | PSIL | 0.629 | 0.702 | 0.472 | 0.059 | 1.248 | 0.259 | 0.333 |
| FC | Global within-network mean | KTM | 0.362 | 0.365 | 0.083 | 0.009 | 0.286 | 0.780 | 0.780 |
| FC | Global within-network mean | N2O | 0.481 | 0.495 | 0.120 | 0.031 | 0.465 | 0.649 | 0.730 |
| FC | Global within-network mean | Sleep N1 | 0.550 | 0.533 | -0.233 | 0.012 | -1.337 | 0.191 | 0.286 |
| FC | Global within-network mean | Sleep N2 | 0.550 | 0.495 | -0.744 | 0.014 | -4.005 | 0.000 | 0.002 |
| FC | Global within-network mean | PPF1.9 | 0.512 | 0.458 | -0.382 | 0.037 | -1.430 | 0.176 | 0.286 |
| FC | Global within-network mean | PPF2.4 | 0.446 | 0.388 | -0.748 | 0.015 | -3.813 | 0.001 | 0.002 |
| FC | Global within-network mean | PPF2.7 | 0.554 | 0.348 | -1.465 | 0.027 | -7.613 | 0.000 | 0.000 |
| Efficiency | Global within-network mean | LSD | 0.804 | 0.833 | 1.259 | 0.006 | 4.878 | 0.000 | 0.001 |
| Efficiency | Global within-network mean | PSIL | 0.864 | 0.891 | 0.718 | 0.014 | 1.899 | 0.106 | 0.137 |
| Efficiency | Global within-network mean | KTM | 0.824 | 0.832 | 0.452 | 0.006 | 1.567 | 0.145 | 0.163 |
| Efficiency | Global within-network mean | N2O | 0.825 | 0.819 | -0.370 | 0.005 | -1.433 | 0.174 | 0.174 |
| Efficiency | Global within-network mean | Sleep N1 | 0.824 | 0.814 | -0.426 | 0.004 | -2.449 | 0.020 | 0.030 |
| Efficiency | Global within-network mean | Sleep N2 | 0.823 | 0.793 | -0.867 | 0.006 | -4.670 | 0.000 | 0.000 |
| Efficiency | Global within-network mean | PPF1.9 | 0.808 | 0.754 | -1.901 | 0.008 | -7.113 | 0.000 | 0.000 |
| Efficiency | Global within-network mean | PPF2.4 | 0.819 | 0.798 | -0.690 | 0.006 | -3.521 | 0.002 | 0.004 |
| Efficiency | Global within-network mean | PPF2.7 | 0.817 | 0.788 | -0.530 | 0.010 | -2.754 | 0.011 | 0.019 |
| Complexity | Global within-network mean | LSD | 0.248 | 0.252 | 0.417 | 0.002 | 1.615 | 0.129 | 0.232 |
| Complexity | Global within-network mean | PSIL | 0.264 | 0.267 | 0.367 | 0.004 | 0.971 | 0.369 | 0.494 |
| Complexity | Global within-network mean | KTM | 0.301 | 0.300 | -0.121 | 0.002 | -0.420 | 0.683 | 0.709 |
| Complexity | Global within-network mean | N2O | 0.286 | 0.283 | -0.232 | 0.003 | -0.898 | 0.384 | 0.494 |
| Complexity | Global within-network mean | Sleep N1 | 0.283 | 0.283 | -0.066 | 0.001 | -0.377 | 0.709 | 0.709 |
| Complexity | Global within-network mean | Sleep N2 | 0.283 | 0.278 | -0.559 | 0.002 | -3.010 | 0.005 | 0.012 |
| Complexity | Global within-network mean | PPF1.9 | 0.288 | 0.277 | -1.068 | 0.003 | -3.995 | 0.002 | 0.005 |
| Complexity | Global within-network mean | PPF2.4 | 0.181 | 0.168 | -1.739 | 0.001 | -8.869 | 0.000 | 0.000 |
| Complexity | Global within-network mean | PPF2.7 | 0.234 | 0.223 | -1.216 | 0.002 | -6.321 | 0.000 | 0.000 |

**Abbreviations:** FC: functional connectivity; LSD: lysergic acid diethylamide; PSIL: psilocybin; KTM: ketamine; N<sub>2</sub>O: nitrous oxide; PPF1.9 / PPF2.4 / PPF2.7: propofol (effect-site concentrations: 1.9 / 2.4 / 2.7 µg·mL<sup>-1</sup>).

**Supplementary Table S4.** Hierarchical results along the unimodal–attention–transmodal (UAT) axis.

| Metric | Pair | State | Baseline mean | State mean | Cohen's d | SE | t | p_unc | p_FDR |
| --- | --- | --- | --- | --- | --- | --- | --- | --- | --- |
| FC | U-U | LSD | 0.325 | 0.265 | -0.821 | 0.019 | -3.179 | 0.007 | 0.015 |
| FC | U-U | PSIL | 0.586 | 0.675 | 0.442 | 0.076 | 1.169 | 0.287 | 0.368 |
| FC | U-U | KTM | 0.338 | 0.344 | 0.126 | 0.013 | 0.438 | 0.670 | 0.754 |
| FC | U-U | N2O | 0.443 | 0.440 | -0.016 | 0.038 | -0.060 | 0.953 | 0.953 |
| FC | U-U | Sleep N1 | 0.545 | 0.524 | -0.221 | 0.016 | -1.271 | 0.213 | 0.319 |
| FC | U-U | Sleep N2 | 0.546 | 0.472 | -0.687 | 0.020 | -3.702 | 0.001 | 0.003 |
| FC | U-U | PPF1.9 | 0.519 | 0.396 | -0.651 | 0.051 | -2.437 | 0.030 | 0.054 |
| FC | U-U | PPF2.4 | 0.482 | 0.340 | -1.266 | 0.022 | -6.454 | 0.000 | 0.000 |
| FC | U-U | PPF2.7 | 0.592 | 0.294 | -1.987 | 0.029 | -10.325 | 0.000 | 0.000 |
| FC | A-A | LSD | 0.278 | 0.292 | 0.145 | 0.024 | 0.561 | 0.583 | 0.601 |
| FC | A-A | PSIL | 0.619 | 0.721 | 0.538 | 0.071 | 1.423 | 0.205 | 0.368 |
| FC | A-A | KTM | 0.343 | 0.354 | 0.214 | 0.015 | 0.740 | 0.475 | 0.601 |
| FC | A-A | N2O | 0.476 | 0.500 | 0.138 | 0.045 | 0.535 | 0.601 | 0.601 |
| FC | A-A | Sleep N1 | 0.489 | 0.469 | -0.261 | 0.014 | -1.500 | 0.143 | 0.323 |
| FC | A-A | Sleep N2 | 0.491 | 0.423 | -0.713 | 0.018 | -3.839 | 0.001 | 0.003 |
| FC | A-A | PPF1.9 | 0.453 | 0.409 | -0.235 | 0.051 | -0.880 | 0.395 | 0.592 |
| FC | A-A | PPF2.4 | 0.382 | 0.338 | -0.489 | 0.018 | -2.496 | 0.020 | 0.059 |
| FC | A-A | PPF2.7 | 0.513 | 0.264 | -1.473 | 0.033 | -7.654 | 0.000 | 0.000 |
| FC | T-T | LSD | 0.148 | 0.187 | 1.089 | 0.009 | 4.219 | 0.001 | 0.003 |
| FC | T-T | PSIL | 0.572 | 0.668 | 0.573 | 0.063 | 1.517 | 0.180 | 0.232 |
| FC | T-T | KTM | 0.268 | 0.291 | 0.575 | 0.012 | 1.992 | 0.072 | 0.161 |
| FC | T-T | N2O | 0.377 | 0.432 | 0.450 | 0.032 | 1.744 | 0.103 | 0.186 |
| FC | T-T | Sleep N1 | 0.456 | 0.439 | -0.247 | 0.012 | -1.419 | 0.166 | 0.232 |
| FC | T-T | Sleep N2 | 0.458 | 0.399 | -0.763 | 0.014 | -4.108 | 0.000 | 0.001 |
| FC | T-T | PPF1.9 | 0.407 | 0.398 | -0.063 | 0.037 | -0.237 | 0.816 | 0.816 |
| FC | T-T | PPF2.4 | 0.321 | 0.310 | -0.143 | 0.015 | -0.728 | 0.474 | 0.533 |
| FC | T-T | PPF2.7 | 0.432 | 0.282 | -0.988 | 0.029 | -5.132 | 0.000 | 0.000 |
| FC | U-A | LSD | 0.201 | 0.213 | 0.159 | 0.020 | 0.616 | 0.548 | 0.548 |
| FC | U-A | PSIL | 0.558 | 0.677 | 0.577 | 0.078 | 1.527 | 0.178 | 0.272 |
| FC | U-A | KTM | 0.273 | 0.299 | 0.545 | 0.014 | 1.890 | 0.085 | 0.192 |
| FC | U-A | N2O | 0.407 | 0.437 | 0.177 | 0.044 | 0.686 | 0.504 | 0.548 |
| FC | U-A | Sleep N1 | 0.465 | 0.445 | -0.208 | 0.016 | -1.198 | 0.240 | 0.308 |
| FC | U-A | Sleep N2 | 0.466 | 0.391 | -0.676 | 0.020 | -3.643 | 0.001 | 0.003 |
| FC | U-A | PPF1.9 | 0.418 | 0.341 | -0.377 | 0.055 | -1.412 | 0.181 | 0.272 |
| FC | U-A | PPF2.4 | 0.345 | 0.266 | -0.762 | 0.020 | -3.886 | 0.001 | 0.003 |
| FC | U-A | PPF2.7 | 0.482 | 0.205 | -1.760 | 0.030 | -9.148 | 0.000 | 0.000 |
| FC | U-T | LSD | -0.018 | 0.022 | 0.842 | 0.012 | 3.260 | 0.006 | 0.014 |
| FC | U-T | PSIL | 0.508 | 0.625 | 0.507 | 0.087 | 1.342 | 0.228 | 0.293 |
| FC | U-T | KTM | 0.142 | 0.206 | 1.071 | 0.017 | 3.710 | 0.003 | 0.014 |
| FC | U-T | N2O | 0.307 | 0.369 | 0.425 | 0.038 | 1.645 | 0.122 | 0.183 |
| FC | U-T | Sleep N1 | 0.377 | 0.371 | -0.058 | 0.019 | -0.335 | 0.740 | 0.740 |
| FC | U-T | Sleep N2 | 0.380 | 0.324 | -0.477 | 0.022 | -2.569 | 0.016 | 0.028 |
| FC | U-T | PPF1.9 | 0.309 | 0.254 | -0.258 | 0.058 | -0.964 | 0.353 | 0.397 |
| FC | U-T | PPF2.4 | 0.225 | 0.163 | -0.585 | 0.021 | -2.981 | 0.006 | 0.014 |
| FC | U-T | PPF2.7 | 0.342 | 0.095 | -1.487 | 0.032 | -7.725 | 0.000 | 0.000 |
| FC | A-T | LSD | -0.003 | 0.045 | 0.990 | 0.012 | 3.835 | 0.002 | 0.008 |
| FC | A-T | PSIL | 0.516 | 0.643 | 0.542 | 0.089 | 1.433 | 0.202 | 0.303 |
| FC | A-T | KTM | 0.198 | 0.244 | 1.089 | 0.012 | 3.774 | 0.003 | 0.009 |
| FC | A-T | N2O | 0.328 | 0.396 | 0.432 | 0.041 | 1.675 | 0.116 | 0.209 |
| FC | A-T | Sleep N1 | 0.384 | 0.375 | -0.088 | 0.018 | -0.508 | 0.615 | 0.692 |
| FC | A-T | Sleep N2 | 0.387 | 0.331 | -0.513 | 0.020 | -2.764 | 0.010 | 0.022 |
| FC | A-T | PPF1.9 | 0.328 | 0.319 | -0.041 | 0.057 | -0.152 | 0.881 | 0.881 |
| FC | A-T | PPF2.4 | 0.248 | 0.235 | -0.128 | 0.019 | -0.652 | 0.520 | 0.669 |
| FC | A-T | PPF2.7 | 0.364 | 0.168 | -1.086 | 0.035 | -5.644 | 0.000 | 0.000 |
| Efficiency | U-U | LSD | 0.687 | 0.735 | 0.855 | 0.015 | 3.313 | 0.005 | 0.009 |
| Efficiency | U-U | PSIL | 0.742 | 0.812 | 0.752 | 0.035 | 1.990 | 0.094 | 0.121 |
| Efficiency | U-U | KTM | 0.752 | 0.759 | 0.219 | 0.009 | 0.758 | 0.464 | 0.522 |
| Efficiency | U-U | N2O | 0.709 | 0.710 | 0.056 | 0.007 | 0.217 | 0.831 | 0.831 |
| Efficiency | U-U | Sleep N1 | 0.694 | 0.671 | -0.552 | 0.007 | -3.174 | 0.003 | 0.007 |
| Efficiency | U-U | Sleep N2 | 0.695 | 0.628 | -1.237 | 0.010 | -6.659 | 0.000 | 0.000 |
| Efficiency | U-U | PPF1.9 | 0.685 | 0.606 | -1.407 | 0.015 | -5.264 | 0.000 | 0.000 |
| Efficiency | U-U | PPF2.4 | 0.703 | 0.645 | -0.870 | 0.013 | -4.435 | 0.000 | 0.000 |
| Efficiency | U-U | PPF2.7 | 0.701 | 0.637 | -0.551 | 0.022 | -2.864 | 0.008 | 0.012 |
| Efficiency | A-A | LSD | 0.749 | 0.798 | 1.006 | 0.012 | 3.895 | 0.002 | 0.004 |
| Efficiency | A-A | PSIL | 0.844 | 0.887 | 0.949 | 0.017 | 2.511 | 0.046 | 0.059 |
| Efficiency | A-A | KTM | 0.788 | 0.819 | 0.625 | 0.014 | 2.164 | 0.053 | 0.060 |
| Efficiency | A-A | N2O | 0.822 | 0.813 | -0.234 | 0.010 | -0.906 | 0.380 | 0.380 |
| Efficiency | A-A | Sleep N1 | 0.803 | 0.788 | -0.527 | 0.005 | -3.027 | 0.005 | 0.007 |
| Efficiency | A-A | Sleep N2 | 0.801 | 0.769 | -0.753 | 0.008 | -4.057 | 0.000 | 0.001 |
| Efficiency | A-A | PPF1.9 | 0.779 | 0.708 | -1.421 | 0.013 | -5.318 | 0.000 | 0.001 |
| Efficiency | A-A | PPF2.4 | 0.797 | 0.764 | -0.650 | 0.010 | -3.314 | 0.003 | 0.005 |
| Efficiency | A-A | PPF2.7 | 0.815 | 0.760 | -0.813 | 0.013 | -4.225 | 0.000 | 0.001 |
| Efficiency | T-T | LSD | 0.746 | 0.789 | 1.162 | 0.009 | 4.500 | 0.000 | 0.003 |
| Efficiency | T-T | PSIL | 0.823 | 0.849 | 0.350 | 0.028 | 0.927 | 0.390 | 0.491 |
| Efficiency | T-T | KTM | 0.780 | 0.795 | 0.381 | 0.011 | 1.321 | 0.213 | 0.384 |
| Efficiency | T-T | N2O | 0.777 | 0.769 | -0.207 | 0.009 | -0.800 | 0.437 | 0.491 |
| Efficiency | T-T | Sleep N1 | 0.793 | 0.790 | -0.099 | 0.006 | -0.571 | 0.572 | 0.572 |
| Efficiency | T-T | Sleep N2 | 0.790 | 0.772 | -0.422 | 0.008 | -2.274 | 0.031 | 0.069 |
| Efficiency | T-T | PPF1.9 | 0.769 | 0.709 | -1.092 | 0.015 | -4.084 | 0.001 | 0.004 |
| Efficiency | T-T | PPF2.4 | 0.783 | 0.776 | -0.203 | 0.007 | -1.033 | 0.312 | 0.467 |
| Efficiency | T-T | PPF2.7 | 0.774 | 0.728 | -0.743 | 0.012 | -3.861 | 0.001 | 0.003 |
| Efficiency | U-A | LSD | 0.648 | 0.719 | 1.319 | 0.014 | 5.109 | 0.000 | 0.001 |
| Efficiency | U-A | PSIL | 0.740 | 0.806 | 0.941 | 0.027 | 2.491 | 0.047 | 0.071 |
| Efficiency | U-A | KTM | 0.700 | 0.711 | 0.273 | 0.011 | 0.947 | 0.364 | 0.409 |
| Efficiency | U-A | N2O | 0.694 | 0.692 | -0.054 | 0.009 | -0.210 | 0.836 | 0.836 |
| Efficiency | U-A | Sleep N1 | 0.660 | 0.645 | -0.462 | 0.006 | -2.651 | 0.012 | 0.027 |
| Efficiency | U-A | Sleep N2 | 0.661 | 0.610 | -1.104 | 0.009 | -5.947 | 0.000 | 0.000 |
| Efficiency | U-A | PPF1.9 | 0.626 | 0.572 | -1.361 | 0.011 | -5.093 | 0.000 | 0.001 |
| Efficiency | U-A | PPF2.4 | 0.658 | 0.625 | -0.513 | 0.013 | -2.617 | 0.015 | 0.027 |
| Efficiency | U-A | PPF2.7 | 0.660 | 0.623 | -0.379 | 0.019 | -1.967 | 0.060 | 0.077 |
| Efficiency | U-T | LSD | 0.591 | 0.667 | 1.268 | 0.016 | 4.909 | 0.000 | 0.001 |
| Efficiency | U-T | PSIL | 0.734 | 0.789 | 0.586 | 0.036 | 1.549 | 0.172 | 0.222 |
| Efficiency | U-T | KTM | 0.683 | 0.703 | 0.604 | 0.009 | 2.091 | 0.061 | 0.091 |
| Efficiency | U-T | N2O | 0.687 | 0.703 | 0.528 | 0.008 | 2.044 | 0.060 | 0.091 |
| Efficiency | U-T | Sleep N1 | 0.656 | 0.645 | -0.359 | 0.005 | -2.065 | 0.047 | 0.091 |
| Efficiency | U-T | Sleep N2 | 0.657 | 0.619 | -0.802 | 0.009 | -4.321 | 0.000 | 0.001 |
| Efficiency | U-T | PPF1.9 | 0.608 | 0.580 | -0.570 | 0.013 | -2.131 | 0.053 | 0.091 |
| Efficiency | U-T | PPF2.4 | 0.631 | 0.630 | -0.006 | 0.016 | -0.031 | 0.976 | 0.976 |
| Efficiency | U-T | PPF2.7 | 0.616 | 0.603 | -0.129 | 0.019 | -0.672 | 0.508 | 0.571 |
| Efficiency | A-T | LSD | 0.656 | 0.699 | 1.048 | 0.011 | 4.060 | 0.001 | 0.004 |
| Efficiency | A-T | PSIL | 0.774 | 0.804 | 0.345 | 0.033 | 0.912 | 0.397 | 0.510 |
| Efficiency | A-T | KTM | 0.707 | 0.729 | 0.487 | 0.013 | 1.688 | 0.120 | 0.179 |
| Efficiency | A-T | N2O | 0.731 | 0.725 | -0.162 | 0.009 | -0.628 | 0.540 | 0.608 |
| Efficiency | A-T | Sleep N1 | 0.735 | 0.734 | -0.049 | 0.004 | -0.284 | 0.778 | 0.778 |
| Efficiency | A-T | Sleep N2 | 0.734 | 0.720 | -0.424 | 0.006 | -2.285 | 0.030 | 0.068 |
| Efficiency | A-T | PPF1.9 | 0.686 | 0.631 | -1.154 | 0.013 | -4.318 | 0.001 | 0.004 |
| Efficiency | A-T | PPF2.4 | 0.728 | 0.709 | -0.361 | 0.010 | -1.841 | 0.077 | 0.139 |
| Efficiency | A-T | PPF2.7 | 0.720 | 0.669 | -0.737 | 0.013 | -3.831 | 0.001 | 0.004 |
| Complexity | U-U | LSD | 0.203 | 0.204 | 0.149 | 0.002 | 0.577 | 0.573 | 0.573 |
| Complexity | U-U | PSIL | 0.211 | 0.216 | 0.565 | 0.003 | 1.496 | 0.185 | 0.322 |
| Complexity | U-U | KTM | 0.247 | 0.243 | -0.380 | 0.002 | -1.318 | 0.214 | 0.322 |
| Complexity | U-U | N2O | 0.230 | 0.227 | -0.284 | 0.003 | -1.099 | 0.290 | 0.330 |
| Complexity | U-U | Sleep N1 | 0.228 | 0.227 | -0.186 | 0.001 | -1.069 | 0.293 | 0.330 |
| Complexity | U-U | Sleep N2 | 0.228 | 0.223 | -0.521 | 0.002 | -2.808 | 0.009 | 0.020 |
| Complexity | U-U | PPF1.9 | 0.234 | 0.224 | -1.371 | 0.002 | -5.129 | 0.000 | 0.001 |
| Complexity | U-U | PPF2.4 | 0.147 | 0.131 | -1.760 | 0.002 | -8.974 | 0.000 | 0.000 |
| Complexity | U-U | PPF2.7 | 0.191 | 0.178 | -1.275 | 0.002 | -6.624 | 0.000 | 0.000 |
| Complexity | A-A | LSD | 0.216 | 0.220 | 0.360 | 0.003 | 1.393 | 0.185 | 0.334 |
| Complexity | A-A | PSIL | 0.228 | 0.232 | 0.351 | 0.004 | 0.930 | 0.388 | 0.466 |
| Complexity | A-A | KTM | 0.264 | 0.266 | 0.245 | 0.002 | 0.849 | 0.414 | 0.466 |
| Complexity | A-A | N2O | 0.251 | 0.251 | -0.057 | 0.004 | -0.220 | 0.829 | 0.829 |
| Complexity | A-A | Sleep N1 | 0.250 | 0.249 | -0.159 | 0.001 | -0.911 | 0.369 | 0.466 |
| Complexity | A-A | Sleep N2 | 0.250 | 0.244 | -0.611 | 0.002 | -3.288 | 0.003 | 0.006 |
| Complexity | A-A | PPF1.9 | 0.255 | 0.242 | -1.241 | 0.003 | -4.644 | 0.000 | 0.001 |
| Complexity | A-A | PPF2.4 | 0.154 | 0.141 | -1.478 | 0.002 | -7.536 | 0.000 | 0.000 |
| Complexity | A-A | PPF2.7 | 0.204 | 0.189 | -1.514 | 0.002 | -7.867 | 0.000 | 0.000 |
| Complexity | T-T | LSD | 0.184 | 0.193 | 1.151 | 0.002 | 4.459 | 0.001 | 0.002 |
| Complexity | T-T | PSIL | 0.207 | 0.212 | 0.437 | 0.005 | 1.156 | 0.292 | 0.375 |
| Complexity | T-T | KTM | 0.235 | 0.239 | 0.406 | 0.002 | 1.408 | 0.187 | 0.280 |
| Complexity | T-T | N2O | 0.229 | 0.227 | -0.102 | 0.003 | -0.396 | 0.698 | 0.785 |
| Complexity | T-T | Sleep N1 | 0.226 | 0.226 | -0.017 | 0.001 | -0.095 | 0.925 | 0.925 |
| Complexity | T-T | Sleep N2 | 0.226 | 0.220 | -0.668 | 0.002 | -3.599 | 0.001 | 0.003 |
| Complexity | T-T | PPF1.9 | 0.228 | 0.220 | -0.829 | 0.003 | -3.101 | 0.008 | 0.015 |
| Complexity | T-T | PPF2.4 | 0.136 | 0.127 | -1.019 | 0.002 | -5.196 | 0.000 | 0.000 |
| Complexity | T-T | PPF2.7 | 0.182 | 0.173 | -1.177 | 0.001 | -6.117 | 0.000 | 0.000 |
| Complexity | U-A | LSD | 0.207 | 0.213 | 0.477 | 0.003 | 1.846 | 0.086 | 0.155 |
| Complexity | U-A | PSIL | 0.223 | 0.231 | 0.598 | 0.005 | 1.583 | 0.164 | 0.247 |
| Complexity | U-A | KTM | 0.258 | 0.260 | 0.184 | 0.003 | 0.636 | 0.538 | 0.616 |
| Complexity | U-A | N2O | 0.246 | 0.245 | -0.090 | 0.004 | -0.347 | 0.734 | 0.734 |
| Complexity | U-A | Sleep N1 | 0.243 | 0.242 | -0.106 | 0.001 | -0.607 | 0.548 | 0.616 |
| Complexity | U-A | Sleep N2 | 0.243 | 0.236 | -0.673 | 0.002 | -3.623 | 0.001 | 0.003 |
| Complexity | U-A | PPF1.9 | 0.247 | 0.234 | -1.106 | 0.003 | -4.140 | 0.001 | 0.003 |
| Complexity | U-A | PPF2.4 | 0.146 | 0.132 | -1.276 | 0.002 | -6.506 | 0.000 | 0.000 |
| Complexity | U-A | PPF2.7 | 0.197 | 0.181 | -1.393 | 0.002 | -7.239 | 0.000 | 0.000 |
| Complexity | U-T | LSD | 0.179 | 0.187 | 1.082 | 0.002 | 4.191 | 0.001 | 0.002 |
| Complexity | U-T | PSIL | 0.203 | 0.211 | 0.543 | 0.005 | 1.435 | 0.201 | 0.302 |
| Complexity | U-T | KTM | 0.229 | 0.232 | 0.302 | 0.003 | 1.048 | 0.317 | 0.408 |
| Complexity | U-T | N2O | 0.222 | 0.222 | 0.011 | 0.003 | 0.041 | 0.968 | 0.968 |
| Complexity | U-T | Sleep N1 | 0.218 | 0.218 | -0.017 | 0.001 | -0.098 | 0.923 | 0.968 |
| Complexity | U-T | Sleep N2 | 0.218 | 0.213 | -0.694 | 0.002 | -3.739 | 0.001 | 0.002 |
| Complexity | U-T | PPF1.9 | 0.220 | 0.211 | -0.809 | 0.003 | -3.026 | 0.010 | 0.018 |
| Complexity | U-T | PPF2.4 | 0.128 | 0.118 | -1.015 | 0.002 | -5.176 | 0.000 | 0.000 |
| Complexity | U-T | PPF2.7 | 0.172 | 0.163 | -1.081 | 0.002 | -5.617 | 0.000 | 0.000 |
| Complexity | A-T | LSD | 0.191 | 0.199 | 1.026 | 0.002 | 3.973 | 0.001 | 0.004 |
| Complexity | A-T | PSIL | 0.219 | 0.226 | 0.390 | 0.007 | 1.031 | 0.342 | 0.440 |
| Complexity | A-T | KTM | 0.250 | 0.254 | 0.546 | 0.002 | 1.891 | 0.085 | 0.128 |
| Complexity | A-T | N2O | 0.239 | 0.241 | 0.117 | 0.004 | 0.452 | 0.658 | 0.658 |
| Complexity | A-T | Sleep N1 | 0.239 | 0.240 | 0.144 | 0.001 | 0.826 | 0.415 | 0.467 |
| Complexity | A-T | Sleep N2 | 0.239 | 0.234 | -0.540 | 0.002 | -2.909 | 0.007 | 0.016 |
| Complexity | A-T | PPF1.9 | 0.242 | 0.232 | -0.731 | 0.004 | -2.737 | 0.017 | 0.031 |
| Complexity | A-T | PPF2.4 | 0.139 | 0.130 | -0.925 | 0.002 | -4.716 | 0.000 | 0.000 |
| Complexity | A-T | PPF2.7 | 0.189 | 0.177 | -1.168 | 0.002 | -6.068 | 0.000 | 0.000 |

**Abbreviations:** FC: functional connectivity; LSD: lysergic acid diethylamide; PSIL: psilocybin; KTM: ketamine; N<sub>2</sub>O: nitrous oxide; PPF1.9 / PPF2.4 / PPF2.7: propofol (effect-site concentrations: 1.9 / 2.4 / 2.7 µg·mL<sup>-1</sup>). U-U: unimodal-unimodal; A-A: attention-attention; T-T: transmodal-transmodal; U-A: unimodal-attention; U-T: unimodal-transmodal; A-T: attention-transmodal.

**Supplementary Table S5.** Within- and between-network results at the 7-network level.

| Metric | Net1 | Net2 | Edge Type | State | Baseline mean | State mean | Cohen's d | SE | t | p_unc | p_FDR |
| --- | --- | --- | --- | --- | --- | --- | --- | --- | --- | --- | --- |
| FC | VIS | VIS | within | LSD | 0.315 | 0.285 | -0.465 | 0.017 | -1.800 | 0.094 | 0.168 |
| FC | VIS | VIS | within | PSIL | 0.655 | 0.703 | 0.252 | 0.073 | 0.665 | 0.530 | 0.682 |
| FC | VIS | VIS | within | KTM | 0.410 | 0.405 | -0.072 | 0.017 | -0.248 | 0.808 | 0.907 |
| FC | VIS | VIS | within | N2O | 0.523 | 0.527 | 0.031 | 0.027 | 0.119 | 0.907 | 0.907 |
| FC | VIS | VIS | within | Sleep N1 | 0.635 | 0.615 | -0.223 | 0.016 | -1.282 | 0.209 | 0.314 |
| FC | VIS | VIS | within | Sleep N2 | 0.638 | 0.549 | -0.822 | 0.020 | -4.426 | 0.000 | 0.000 |
| FC | VIS | VIS | within | PPF1.9 | 0.567 | 0.459 | -0.676 | 0.043 | -2.529 | 0.025 | 0.057 |
| FC | VIS | VIS | within | PPF2.4 | 0.534 | 0.422 | -1.174 | 0.019 | -5.986 | 0.000 | 0.000 |
| FC | VIS | VIS | within | PPF2.7 | 0.648 | 0.402 | -1.766 | 0.027 | -9.174 | 0.000 | 0.000 |
| FC | VIS | SMN | between | LSD | 0.220 | 0.174 | -0.546 | 0.022 | -2.114 | 0.053 | 0.095 |
| FC | VIS | SMN | between | PSIL | 0.524 | 0.632 | 0.455 | 0.090 | 1.203 | 0.274 | 0.316 |
| FC | VIS | SMN | between | KTM | 0.254 | 0.271 | 0.328 | 0.015 | 1.135 | 0.281 | 0.316 |
| FC | VIS | SMN | between | N2O | 0.349 | 0.352 | 0.018 | 0.043 | 0.069 | 0.946 | 0.946 |
| FC | VIS | SMN | between | Sleep N1 | 0.460 | 0.437 | -0.213 | 0.019 | -1.222 | 0.231 | 0.316 |
| FC | VIS | SMN | between | Sleep N2 | 0.463 | 0.376 | -0.670 | 0.024 | -3.610 | 0.001 | 0.004 |
| FC | VIS | SMN | between | PPF1.9 | 0.448 | 0.294 | -0.726 | 0.057 | -2.717 | 0.018 | 0.040 |
| FC | VIS | SMN | between | PPF2.4 | 0.413 | 0.239 | -1.401 | 0.024 | -7.145 | 0.000 | 0.000 |
| FC | VIS | SMN | between | PPF2.7 | 0.523 | 0.192 | -2.158 | 0.030 | -11.212 | 0.000 | 0.000 |
| FC | VIS | DAN | between | LSD | 0.141 | 0.175 | 0.530 | 0.017 | 2.052 | 0.059 | 0.107 |
| FC | VIS | DAN | between | PSIL | 0.570 | 0.669 | 0.470 | 0.080 | 1.243 | 0.260 | 0.335 |
| FC | VIS | DAN | between | KTM | 0.235 | 0.273 | 0.818 | 0.013 | 2.833 | 0.016 | 0.037 |
| FC | VIS | DAN | between | N2O | 0.378 | 0.412 | 0.204 | 0.043 | 0.790 | 0.443 | 0.443 |
| FC | VIS | DAN | between | Sleep N1 | 0.478 | 0.460 | -0.179 | 0.017 | -1.031 | 0.310 | 0.349 |
| FC | VIS | DAN | between | Sleep N2 | 0.482 | 0.381 | -0.764 | 0.025 | -4.114 | 0.000 | 0.001 |
| FC | VIS | DAN | between | PPF1.9 | 0.416 | 0.338 | -0.399 | 0.053 | -1.494 | 0.159 | 0.239 |
| FC | VIS | DAN | between | PPF2.4 | 0.369 | 0.275 | -0.821 | 0.022 | -4.189 | 0.000 | 0.001 |
| FC | VIS | DAN | between | PPF2.7 | 0.488 | 0.243 | -1.650 | 0.029 | -8.575 | 0.000 | 0.000 |
| FC | VIS | VAN | between | LSD | 0.118 | 0.148 | 0.328 | 0.023 | 1.270 | 0.225 | 0.325 |
| FC | VIS | VAN | between | PSIL | 0.521 | 0.631 | 0.441 | 0.094 | 1.166 | 0.288 | 0.325 |
| FC | VIS | VAN | between | KTM | 0.190 | 0.228 | 0.679 | 0.016 | 2.352 | 0.038 | 0.086 |
| FC | VIS | VAN | between | N2O | 0.337 | 0.347 | 0.061 | 0.040 | 0.238 | 0.815 | 0.815 |
| FC | VIS | VAN | between | Sleep N1 | 0.380 | 0.361 | -0.188 | 0.017 | -1.079 | 0.289 | 0.325 |
| FC | VIS | VAN | between | Sleep N2 | 0.378 | 0.317 | -0.549 | 0.021 | -2.958 | 0.006 | 0.019 |
| FC | VIS | VAN | between | PPF1.9 | 0.327 | 0.247 | -0.412 | 0.052 | -1.541 | 0.147 | 0.265 |
| FC | VIS | VAN | between | PPF2.4 | 0.263 | 0.191 | -0.722 | 0.020 | -3.680 | 0.001 | 0.005 |
| FC | VIS | VAN | between | PPF2.7 | 0.405 | 0.146 | -1.657 | 0.030 | -8.609 | 0.000 | 0.000 |
| FC | VIS | LIM | between | LSD | 0.036 | 0.032 | -0.072 | 0.011 | -0.281 | 0.783 | 0.848 |
| FC | VIS | LIM | between | PSIL | 0.462 | 0.531 | 0.293 | 0.090 | 0.775 | 0.468 | 0.601 |
| FC | VIS | LIM | between | KTM | 0.154 | 0.192 | 0.628 | 0.018 | 2.176 | 0.052 | 0.094 |
| FC | VIS | LIM | between | N2O | 0.259 | 0.233 | -0.205 | 0.033 | -0.793 | 0.441 | 0.601 |
| FC | VIS | LIM | between | Sleep N1 | 0.346 | 0.342 | -0.034 | 0.020 | -0.193 | 0.848 | 0.848 |
| FC | VIS | LIM | between | Sleep N2 | 0.344 | 0.298 | -0.479 | 0.018 | -2.581 | 0.015 | 0.035 |
| FC | VIS | LIM | between | PPF1.9 | 0.311 | 0.165 | -0.915 | 0.042 | -3.425 | 0.005 | 0.014 |
| FC | VIS | LIM | between | PPF2.4 | 0.305 | 0.182 | -1.410 | 0.017 | -7.190 | 0.000 | 0.000 |
| FC | VIS | LIM | between | PPF2.7 | 0.300 | 0.089 | -2.105 | 0.019 | -10.938 | 0.000 | 0.000 |
| FC | VIS | FPN | between | LSD | -0.025 | 0.042 | 0.973 | 0.018 | 3.769 | 0.002 | 0.006 |
| FC | VIS | FPN | between | PSIL | 0.514 | 0.629 | 0.532 | 0.081 | 1.408 | 0.209 | 0.269 |
| FC | VIS | FPN | between | KTM | 0.114 | 0.184 | 0.925 | 0.022 | 3.206 | 0.008 | 0.019 |
| FC | VIS | FPN | between | N2O | 0.282 | 0.349 | 0.543 | 0.032 | 2.102 | 0.054 | 0.089 |
| FC | VIS | FPN | between | Sleep N1 | 0.350 | 0.337 | -0.140 | 0.017 | -0.805 | 0.427 | 0.480 |
| FC | VIS | FPN | between | Sleep N2 | 0.351 | 0.284 | -0.674 | 0.018 | -3.627 | 0.001 | 0.005 |
| FC | VIS | FPN | between | PPF1.9 | 0.279 | 0.246 | -0.193 | 0.045 | -0.724 | 0.482 | 0.482 |
| FC | VIS | FPN | between | PPF2.4 | 0.214 | 0.179 | -0.388 | 0.017 | -1.978 | 0.059 | 0.089 |
| FC | VIS | FPN | between | PPF2.7 | 0.318 | 0.133 | -1.110 | 0.032 | -5.768 | 0.000 | 0.000 |
| FC | VIS | DMN | between | LSD | 0.016 | 0.056 | 0.979 | 0.011 | 3.790 | 0.002 | 0.005 |
| FC | VIS | DMN | between | PSIL | 0.504 | 0.615 | 0.541 | 0.077 | 1.431 | 0.202 | 0.260 |
| FC | VIS | DMN | between | KTM | 0.140 | 0.205 | 1.008 | 0.019 | 3.493 | 0.005 | 0.009 |
| FC | VIS | DMN | between | N2O | 0.322 | 0.356 | 0.269 | 0.033 | 1.041 | 0.316 | 0.355 |
| FC | VIS | DMN | between | Sleep N1 | 0.402 | 0.391 | -0.099 | 0.019 | -0.569 | 0.573 | 0.573 |
| FC | VIS | DMN | between | Sleep N2 | 0.403 | 0.328 | -0.632 | 0.022 | -3.406 | 0.002 | 0.005 |
| FC | VIS | DMN | between | PPF1.9 | 0.314 | 0.228 | -0.416 | 0.055 | -1.555 | 0.144 | 0.216 |
| FC | VIS | DMN | between | PPF2.4 | 0.245 | 0.165 | -0.817 | 0.019 | -4.165 | 0.000 | 0.001 |
| FC | VIS | DMN | between | PPF2.7 | 0.355 | 0.105 | -1.553 | 0.031 | -8.068 | 0.000 | 0.000 |
| FC | SMN | SMN | within | LSD | 0.497 | 0.397 | -0.909 | 0.028 | -3.521 | 0.003 | 0.010 |
| FC | SMN | SMN | within | PSIL | 0.641 | 0.724 | 0.506 | 0.062 | 1.338 | 0.229 | 0.344 |
| FC | SMN | SMN | within | KTM | 0.427 | 0.420 | -0.125 | 0.015 | -0.433 | 0.674 | 0.758 |
| FC | SMN | SMN | within | N2O | 0.540 | 0.526 | -0.079 | 0.046 | -0.306 | 0.764 | 0.764 |
| FC | SMN | SMN | within | Sleep N1 | 0.621 | 0.605 | -0.196 | 0.014 | -1.125 | 0.269 | 0.346 |
| FC | SMN | SMN | within | Sleep N2 | 0.621 | 0.576 | -0.463 | 0.018 | -2.495 | 0.019 | 0.042 |
| FC | SMN | SMN | within | PPF1.9 | 0.603 | 0.519 | -0.421 | 0.053 | -1.574 | 0.139 | 0.251 |
| FC | SMN | SMN | within | PPF2.4 | 0.557 | 0.450 | -0.849 | 0.025 | -4.331 | 0.000 | 0.001 |
| FC | SMN | SMN | within | PPF2.7 | 0.666 | 0.386 | -1.623 | 0.033 | -8.432 | 0.000 | 0.000 |
| FC | SMN | DAN | between | LSD | 0.220 | 0.232 | 0.101 | 0.033 | 0.391 | 0.702 | 0.702 |
| FC | SMN | DAN | between | PSIL | 0.560 | 0.691 | 0.658 | 0.076 | 1.741 | 0.132 | 0.206 |
| FC | SMN | DAN | between | KTM | 0.308 | 0.335 | 0.489 | 0.016 | 1.693 | 0.119 | 0.206 |
| FC | SMN | DAN | between | N2O | 0.433 | 0.487 | 0.295 | 0.047 | 1.141 | 0.273 | 0.307 |
| FC | SMN | DAN | between | Sleep N1 | 0.503 | 0.478 | -0.250 | 0.017 | -1.437 | 0.161 | 0.206 |
| FC | SMN | DAN | between | Sleep N2 | 0.507 | 0.408 | -0.758 | 0.024 | -4.080 | 0.000 | 0.002 |
| FC | SMN | DAN | between | PPF1.9 | 0.462 | 0.363 | -0.419 | 0.064 | -1.567 | 0.141 | 0.206 |
| FC | SMN | DAN | between | PPF2.4 | 0.368 | 0.274 | -0.765 | 0.024 | -3.899 | 0.001 | 0.002 |
| FC | SMN | DAN | between | PPF2.7 | 0.503 | 0.182 | -2.053 | 0.030 | -10.666 | 0.000 | 0.000 |
| FC | SMN | VAN | between | LSD | 0.294 | 0.275 | -0.207 | 0.024 | -0.800 | 0.437 | 0.562 |
| FC | SMN | VAN | between | PSIL | 0.576 | 0.705 | 0.653 | 0.075 | 1.727 | 0.135 | 0.303 |
| FC | SMN | VAN | between | KTM | 0.333 | 0.338 | 0.094 | 0.017 | 0.324 | 0.752 | 0.752 |
| FC | SMN | VAN | between | N2O | 0.458 | 0.478 | 0.103 | 0.050 | 0.398 | 0.697 | 0.752 |
| FC | SMN | VAN | between | Sleep N1 | 0.484 | 0.469 | -0.169 | 0.016 | -0.969 | 0.340 | 0.562 |
| FC | SMN | VAN | between | Sleep N2 | 0.482 | 0.443 | -0.392 | 0.019 | -2.110 | 0.044 | 0.132 |
| FC | SMN | VAN | between | PPF1.9 | 0.449 | 0.396 | -0.241 | 0.059 | -0.902 | 0.383 | 0.562 |
| FC | SMN | VAN | between | PPF2.4 | 0.370 | 0.311 | -0.505 | 0.023 | -2.576 | 0.016 | 0.073 |
| FC | SMN | VAN | between | PPF2.7 | 0.517 | 0.245 | -1.456 | 0.036 | -7.564 | 0.000 | 0.000 |
| FC | SMN | LIM | between | LSD | 0.024 | 0.015 | -0.137 | 0.018 | -0.531 | 0.604 | 0.827 |
| FC | SMN | LIM | between | PSIL | 0.479 | 0.528 | 0.184 | 0.100 | 0.488 | 0.643 | 0.827 |
| FC | SMN | LIM | between | KTM | 0.160 | 0.179 | 0.231 | 0.024 | 0.802 | 0.440 | 0.791 |
| FC | SMN | LIM | between | N2O | 0.183 | 0.182 | -0.008 | 0.042 | -0.031 | 0.976 | 0.976 |
| FC | SMN | LIM | between | Sleep N1 | 0.320 | 0.319 | -0.007 | 0.019 | -0.039 | 0.969 | 0.976 |
| FC | SMN | LIM | between | Sleep N2 | 0.319 | 0.290 | -0.278 | 0.019 | -1.497 | 0.146 | 0.328 |
| FC | SMN | LIM | between | PPF1.9 | 0.319 | 0.144 | -0.842 | 0.055 | -3.152 | 0.008 | 0.023 |
| FC | SMN | LIM | between | PPF2.4 | 0.294 | 0.191 | -0.900 | 0.023 | -4.590 | 0.000 | 0.000 |
| FC | SMN | LIM | between | PPF2.7 | 0.292 | 0.075 | -2.247 | 0.019 | -11.674 | 0.000 | 0.000 |
| FC | SMN | FPN | between | LSD | -0.040 | 0.014 | 0.520 | 0.027 | 2.013 | 0.064 | 0.144 |
| FC | SMN | FPN | between | PSIL | 0.520 | 0.639 | 0.504 | 0.089 | 1.332 | 0.231 | 0.297 |
| FC | SMN | FPN | between | KTM | 0.161 | 0.220 | 0.839 | 0.020 | 2.906 | 0.014 | 0.053 |
| FC | SMN | FPN | between | N2O | 0.310 | 0.391 | 0.474 | 0.044 | 1.837 | 0.088 | 0.158 |
| FC | SMN | FPN | between | Sleep N1 | 0.361 | 0.352 | -0.079 | 0.019 | -0.454 | 0.653 | 0.653 |
| FC | SMN | FPN | between | Sleep N2 | 0.367 | 0.307 | -0.468 | 0.024 | -2.522 | 0.018 | 0.053 |
| FC | SMN | FPN | between | PPF1.9 | 0.299 | 0.269 | -0.124 | 0.064 | -0.464 | 0.651 | 0.653 |
| FC | SMN | FPN | between | PPF2.4 | 0.195 | 0.157 | -0.323 | 0.023 | -1.648 | 0.112 | 0.168 |
| FC | SMN | FPN | between | PPF2.7 | 0.336 | 0.069 | -1.421 | 0.036 | -7.384 | 0.000 | 0.000 |
| FC | SMN | DMN | between | LSD | -0.029 | -0.009 | 0.324 | 0.016 | 1.254 | 0.230 | 0.339 |
| FC | SMN | DMN | between | PSIL | 0.500 | 0.623 | 0.466 | 0.099 | 1.234 | 0.263 | 0.339 |
| FC | SMN | DMN | between | KTM | 0.145 | 0.209 | 1.019 | 0.018 | 3.528 | 0.005 | 0.021 |
| FC | SMN | DMN | between | N2O | 0.305 | 0.376 | 0.385 | 0.048 | 1.492 | 0.158 | 0.334 |
| FC | SMN | DMN | between | Sleep N1 | 0.379 | 0.381 | 0.016 | 0.021 | 0.091 | 0.928 | 0.928 |
| FC | SMN | DMN | between | Sleep N2 | 0.384 | 0.348 | -0.252 | 0.026 | -1.357 | 0.186 | 0.334 |
| FC | SMN | DMN | between | PPF1.9 | 0.326 | 0.268 | -0.232 | 0.066 | -0.870 | 0.400 | 0.450 |
| FC | SMN | DMN | between | PPF2.4 | 0.231 | 0.158 | -0.554 | 0.026 | -2.824 | 0.009 | 0.028 |
| FC | SMN | DMN | between | PPF2.7 | 0.346 | 0.085 | -1.515 | 0.033 | -7.871 | 0.000 | 0.000 |
| FC | DAN | DAN | within | LSD | 0.345 | 0.358 | 0.163 | 0.020 | 0.630 | 0.539 | 0.539 |
| FC | DAN | DAN | within | PSIL | 0.666 | 0.747 | 0.531 | 0.057 | 1.405 | 0.210 | 0.454 |
| FC | DAN | DAN | within | KTM | 0.396 | 0.413 | 0.312 | 0.015 | 1.081 | 0.303 | 0.454 |
| FC | DAN | DAN | within | N2O | 0.530 | 0.562 | 0.202 | 0.042 | 0.781 | 0.448 | 0.504 |
| FC | DAN | DAN | within | Sleep N1 | 0.573 | 0.559 | -0.187 | 0.013 | -1.076 | 0.290 | 0.454 |
| FC | DAN | DAN | within | Sleep N2 | 0.580 | 0.498 | -0.795 | 0.019 | -4.280 | 0.000 | 0.001 |
| FC | DAN | DAN | within | PPF1.9 | 0.544 | 0.500 | -0.242 | 0.049 | -0.904 | 0.383 | 0.492 |
| FC | DAN | DAN | within | PPF2.4 | 0.464 | 0.425 | -0.436 | 0.018 | -2.221 | 0.036 | 0.107 |
| FC | DAN | DAN | within | PPF2.7 | 0.579 | 0.373 | -1.416 | 0.028 | -7.360 | 0.000 | 0.000 |
| FC | DAN | VAN | between | LSD | 0.191 | 0.216 | 0.205 | 0.032 | 0.793 | 0.441 | 0.459 |
| FC | DAN | VAN | between | PSIL | 0.582 | 0.698 | 0.552 | 0.079 | 1.462 | 0.194 | 0.349 |
| FC | DAN | VAN | between | KTM | 0.292 | 0.313 | 0.328 | 0.018 | 1.135 | 0.281 | 0.421 |
| FC | DAN | VAN | between | N2O | 0.432 | 0.468 | 0.200 | 0.047 | 0.774 | 0.452 | 0.459 |
| FC | DAN | VAN | between | Sleep N1 | 0.441 | 0.417 | -0.282 | 0.015 | -1.618 | 0.116 | 0.260 |
| FC | DAN | VAN | between | Sleep N2 | 0.442 | 0.365 | -0.728 | 0.020 | -3.918 | 0.001 | 0.002 |
| FC | DAN | VAN | between | PPF1.9 | 0.395 | 0.352 | -0.204 | 0.056 | -0.763 | 0.459 | 0.459 |
| FC | DAN | VAN | between | PPF2.4 | 0.318 | 0.278 | -0.425 | 0.019 | -2.169 | 0.040 | 0.119 |
| FC | DAN | VAN | between | PPF2.7 | 0.459 | 0.194 | -1.471 | 0.035 | -7.643 | 0.000 | 0.000 |
| FC | DAN | LIM | between | LSD | 0.003 | 0.009 | 0.132 | 0.013 | 0.511 | 0.617 | 0.794 |
| FC | DAN | LIM | between | PSIL | 0.473 | 0.544 | 0.273 | 0.099 | 0.723 | 0.497 | 0.746 |
| FC | DAN | LIM | between | KTM | 0.172 | 0.185 | 0.209 | 0.018 | 0.725 | 0.483 | 0.746 |
| FC | DAN | LIM | between | N2O | 0.230 | 0.224 | -0.031 | 0.044 | -0.121 | 0.905 | 0.905 |
| FC | DAN | LIM | between | Sleep N1 | 0.342 | 0.339 | -0.033 | 0.017 | -0.191 | 0.849 | 0.905 |
| FC | DAN | LIM | between | Sleep N2 | 0.341 | 0.297 | -0.493 | 0.017 | -2.654 | 0.013 | 0.032 |
| FC | DAN | LIM | between | PPF1.9 | 0.322 | 0.165 | -0.755 | 0.056 | -2.823 | 0.014 | 0.032 |
| FC | DAN | LIM | between | PPF2.4 | 0.282 | 0.195 | -0.813 | 0.021 | -4.147 | 0.000 | 0.002 |
| FC | DAN | LIM | between | PPF2.7 | 0.293 | 0.100 | -1.587 | 0.023 | -8.247 | 0.000 | 0.000 |
| FC | DAN | FPN | between | LSD | 0.092 | 0.120 | 0.399 | 0.018 | 1.547 | 0.144 | 0.260 |
| FC | DAN | FPN | between | PSIL | 0.583 | 0.679 | 0.507 | 0.072 | 1.342 | 0.228 | 0.342 |
| FC | DAN | FPN | between | KTM | 0.253 | 0.281 | 0.493 | 0.016 | 1.706 | 0.116 | 0.260 |
| FC | DAN | FPN | between | N2O | 0.376 | 0.437 | 0.419 | 0.037 | 1.623 | 0.127 | 0.260 |
| FC | DAN | FPN | between | Sleep N1 | 0.431 | 0.419 | -0.141 | 0.015 | -0.811 | 0.424 | 0.545 |
| FC | DAN | FPN | between | Sleep N2 | 0.434 | 0.372 | -0.665 | 0.017 | -3.582 | 0.001 | 0.006 |
| FC | DAN | FPN | between | PPF1.9 | 0.372 | 0.378 | 0.035 | 0.048 | 0.132 | 0.897 | 0.954 |
| FC | DAN | FPN | between | PPF2.4 | 0.292 | 0.293 | 0.011 | 0.014 | 0.058 | 0.954 | 0.954 |
| FC | DAN | FPN | between | PPF2.7 | 0.413 | 0.241 | -1.022 | 0.032 | -5.308 | 0.000 | 0.000 |
| FC | DAN | DMN | between | LSD | -0.064 | 0.002 | 1.016 | 0.017 | 3.936 | 0.001 | 0.007 |
| FC | DAN | DMN | between | PSIL | 0.485 | 0.643 | 0.648 | 0.092 | 1.715 | 0.137 | 0.176 |
| FC | DAN | DMN | between | KTM | 0.170 | 0.227 | 1.147 | 0.015 | 3.972 | 0.002 | 0.007 |
| FC | DAN | DMN | between | N2O | 0.304 | 0.394 | 0.526 | 0.044 | 2.037 | 0.061 | 0.110 |
| FC | DAN | DMN | between | Sleep N1 | 0.389 | 0.380 | -0.084 | 0.020 | -0.484 | 0.631 | 0.631 |
| FC | DAN | DMN | between | Sleep N2 | 0.393 | 0.322 | -0.543 | 0.024 | -2.922 | 0.007 | 0.015 |
| FC | DAN | DMN | between | PPF1.9 | 0.319 | 0.276 | -0.175 | 0.067 | -0.654 | 0.524 | 0.590 |
| FC | DAN | DMN | between | PPF2.4 | 0.230 | 0.192 | -0.333 | 0.022 | -1.697 | 0.102 | 0.153 |
| FC | DAN | DMN | between | PPF2.7 | 0.353 | 0.121 | -1.193 | 0.038 | -6.200 | 0.000 | 0.000 |
| FC | VAN | VAN | within | LSD | 0.385 | 0.376 | -0.097 | 0.023 | -0.377 | 0.712 | 0.801 |
| FC | VAN | VAN | within | PSIL | 0.646 | 0.739 | 0.498 | 0.070 | 1.317 | 0.236 | 0.424 |
| FC | VAN | VAN | within | KTM | 0.390 | 0.379 | -0.247 | 0.013 | -0.854 | 0.411 | 0.529 |
| FC | VAN | VAN | within | N2O | 0.512 | 0.502 | -0.055 | 0.047 | -0.212 | 0.836 | 0.836 |
| FC | VAN | VAN | within | Sleep N1 | 0.504 | 0.483 | -0.248 | 0.014 | -1.425 | 0.164 | 0.369 |
| FC | VAN | VAN | within | Sleep N2 | 0.501 | 0.463 | -0.422 | 0.017 | -2.271 | 0.031 | 0.093 |
| FC | VAN | VAN | within | PPF1.9 | 0.480 | 0.432 | -0.270 | 0.048 | -1.009 | 0.331 | 0.497 |
| FC | VAN | VAN | within | PPF2.4 | 0.430 | 0.372 | -0.551 | 0.021 | -2.812 | 0.009 | 0.042 |
| FC | VAN | VAN | within | PPF2.7 | 0.557 | 0.296 | -1.368 | 0.037 | -7.111 | 0.000 | 0.000 |
| FC | VAN | LIM | between | LSD | 0.021 | 0.007 | -0.276 | 0.013 | -1.069 | 0.303 | 0.546 |
| FC | VAN | LIM | between | PSIL | 0.466 | 0.523 | 0.212 | 0.102 | 0.560 | 0.596 | 0.894 |
| FC | VAN | LIM | between | KTM | 0.170 | 0.177 | 0.083 | 0.023 | 0.287 | 0.780 | 0.961 |
| FC | VAN | LIM | between | N2O | 0.170 | 0.171 | 0.002 | 0.045 | 0.006 | 0.995 | 0.995 |
| FC | VAN | LIM | between | Sleep N1 | 0.275 | 0.272 | -0.032 | 0.016 | -0.185 | 0.854 | 0.961 |
| FC | VAN | LIM | between | Sleep N2 | 0.274 | 0.253 | -0.256 | 0.016 | -1.380 | 0.179 | 0.402 |
| FC | VAN | LIM | between | PPF1.9 | 0.277 | 0.172 | -0.558 | 0.050 | -2.089 | 0.057 | 0.171 |
| FC | VAN | LIM | between | PPF2.4 | 0.243 | 0.196 | -0.412 | 0.022 | -2.102 | 0.046 | 0.171 |
| FC | VAN | LIM | between | PPF2.7 | 0.270 | 0.106 | -1.416 | 0.022 | -7.357 | 0.000 | 0.000 |
| FC | VAN | FPN | between | LSD | 0.052 | 0.098 | 0.526 | 0.023 | 2.038 | 0.061 | 0.137 |
| FC | VAN | FPN | between | PSIL | 0.556 | 0.651 | 0.419 | 0.085 | 1.109 | 0.310 | 0.419 |
| FC | VAN | FPN | between | KTM | 0.231 | 0.262 | 0.639 | 0.014 | 2.212 | 0.049 | 0.137 |
| FC | VAN | FPN | between | N2O | 0.359 | 0.403 | 0.274 | 0.041 | 1.060 | 0.307 | 0.419 |
| FC | VAN | FPN | between | Sleep N1 | 0.380 | 0.365 | -0.174 | 0.016 | -0.998 | 0.326 | 0.419 |
| FC | VAN | FPN | between | Sleep N2 | 0.382 | 0.319 | -0.613 | 0.019 | -3.302 | 0.003 | 0.012 |
| FC | VAN | FPN | between | PPF1.9 | 0.330 | 0.334 | 0.021 | 0.053 | 0.080 | 0.937 | 0.937 |
| FC | VAN | FPN | between | PPF2.4 | 0.262 | 0.253 | -0.107 | 0.017 | -0.545 | 0.591 | 0.664 |
| FC | VAN | FPN | between | PPF2.7 | 0.378 | 0.170 | -1.079 | 0.037 | -5.605 | 0.000 | 0.000 |
| FC | VAN | DMN | between | LSD | -0.028 | 0.014 | 0.810 | 0.013 | 3.135 | 0.007 | 0.022 |
| FC | VAN | DMN | between | PSIL | 0.485 | 0.618 | 0.501 | 0.100 | 1.327 | 0.233 | 0.349 |
| FC | VAN | DMN | between | KTM | 0.176 | 0.230 | 1.211 | 0.013 | 4.194 | 0.002 | 0.007 |
| FC | VAN | DMN | between | N2O | 0.307 | 0.372 | 0.391 | 0.043 | 1.514 | 0.152 | 0.274 |
| FC | VAN | DMN | between | Sleep N1 | 0.356 | 0.353 | -0.022 | 0.018 | -0.126 | 0.900 | 0.913 |
| FC | VAN | DMN | between | Sleep N2 | 0.357 | 0.324 | -0.296 | 0.021 | -1.595 | 0.122 | 0.274 |
| FC | VAN | DMN | between | PPF1.9 | 0.310 | 0.320 | 0.045 | 0.058 | 0.169 | 0.869 | 0.913 |
| FC | VAN | DMN | between | PPF2.4 | 0.233 | 0.236 | 0.022 | 0.024 | 0.111 | 0.913 | 0.913 |
| FC | VAN | DMN | between | PPF2.7 | 0.338 | 0.171 | -0.896 | 0.036 | -4.658 | 0.000 | 0.001 |
| FC | LIM | LIM | within | LSD | 0.186 | 0.189 | 0.048 | 0.016 | 0.186 | 0.855 | 0.855 |
| FC | LIM | LIM | within | PSIL | 0.509 | 0.527 | 0.084 | 0.083 | 0.222 | 0.832 | 0.855 |
| FC | LIM | LIM | within | KTM | 0.313 | 0.282 | -0.374 | 0.024 | -1.294 | 0.222 | 0.496 |
| FC | LIM | LIM | within | N2O | 0.443 | 0.450 | 0.091 | 0.020 | 0.351 | 0.731 | 0.855 |
| FC | LIM | LIM | within | Sleep N1 | 0.426 | 0.432 | 0.069 | 0.016 | 0.396 | 0.695 | 0.855 |
| FC | LIM | LIM | within | Sleep N2 | 0.421 | 0.410 | -0.206 | 0.010 | -1.112 | 0.276 | 0.496 |
| FC | LIM | LIM | within | PPF1.9 | 0.367 | 0.296 | -0.441 | 0.043 | -1.649 | 0.123 | 0.369 |
| FC | LIM | LIM | within | PPF2.4 | 0.366 | 0.317 | -0.494 | 0.019 | -2.520 | 0.018 | 0.083 |
| FC | LIM | LIM | within | PPF2.7 | 0.283 | 0.159 | -1.430 | 0.017 | -7.431 | 0.000 | 0.000 |
| FC | LIM | FPN | between | LSD | 0.024 | 0.042 | 0.365 | 0.013 | 1.413 | 0.180 | 0.323 |
| FC | LIM | FPN | between | PSIL | 0.483 | 0.538 | 0.236 | 0.088 | 0.624 | 0.555 | 0.714 |
| FC | LIM | FPN | between | KTM | 0.166 | 0.164 | -0.034 | 0.015 | -0.117 | 0.909 | 0.909 |
| FC | LIM | FPN | between | N2O | 0.195 | 0.221 | 0.213 | 0.031 | 0.826 | 0.423 | 0.634 |
| FC | LIM | FPN | between | Sleep N1 | 0.308 | 0.305 | -0.036 | 0.014 | -0.208 | 0.837 | 0.909 |
| FC | LIM | FPN | between | Sleep N2 | 0.308 | 0.275 | -0.478 | 0.013 | -2.572 | 0.016 | 0.050 |
| FC | LIM | FPN | between | PPF1.9 | 0.299 | 0.198 | -0.549 | 0.049 | -2.053 | 0.061 | 0.137 |
| FC | LIM | FPN | between | PPF2.4 | 0.234 | 0.194 | -0.503 | 0.016 | -2.565 | 0.017 | 0.050 |
| FC | LIM | FPN | between | PPF2.7 | 0.256 | 0.122 | -1.150 | 0.023 | -5.974 | 0.000 | 0.000 |
| FC | LIM | DMN | between | LSD | 0.084 | 0.098 | 0.281 | 0.013 | 1.088 | 0.295 | 0.531 |
| FC | LIM | DMN | between | PSIL | 0.512 | 0.547 | 0.178 | 0.074 | 0.471 | 0.654 | 0.841 |
| FC | LIM | DMN | between | KTM | 0.167 | 0.179 | 0.221 | 0.016 | 0.764 | 0.461 | 0.691 |
| FC | LIM | DMN | between | N2O | 0.280 | 0.277 | -0.027 | 0.028 | -0.105 | 0.918 | 0.918 |
| FC | LIM | DMN | between | Sleep N1 | 0.345 | 0.347 | 0.026 | 0.015 | 0.150 | 0.881 | 0.918 |
| FC | LIM | DMN | between | Sleep N2 | 0.344 | 0.315 | -0.441 | 0.012 | -2.374 | 0.025 | 0.056 |
| FC | LIM | DMN | between | PPF1.9 | 0.338 | 0.213 | -0.701 | 0.048 | -2.625 | 0.021 | 0.056 |
| FC | LIM | DMN | between | PPF2.4 | 0.290 | 0.210 | -0.907 | 0.017 | -4.623 | 0.000 | 0.000 |
| FC | LIM | DMN | between | PPF2.7 | 0.293 | 0.141 | -1.395 | 0.021 | -7.250 | 0.000 | 0.000 |
| FC | FPN | FPN | within | LSD | 0.269 | 0.262 | -0.151 | 0.013 | -0.585 | 0.568 | 0.798 |
| FC | FPN | FPN | within | PSIL | 0.610 | 0.692 | 0.479 | 0.065 | 1.267 | 0.252 | 0.567 |
| FC | FPN | FPN | within | KTM | 0.323 | 0.319 | -0.091 | 0.012 | -0.314 | 0.759 | 0.854 |
| FC | FPN | FPN | within | N2O | 0.406 | 0.438 | 0.267 | 0.031 | 1.034 | 0.319 | 0.573 |
| FC | FPN | FPN | within | Sleep N1 | 0.473 | 0.459 | -0.205 | 0.012 | -1.177 | 0.248 | 0.567 |
| FC | FPN | FPN | within | Sleep N2 | 0.473 | 0.422 | -0.659 | 0.015 | -3.547 | 0.001 | 0.006 |
| FC | FPN | FPN | within | PPF1.9 | 0.431 | 0.448 | 0.136 | 0.033 | 0.507 | 0.621 | 0.798 |
| FC | FPN | FPN | within | PPF2.4 | 0.361 | 0.361 | 0.003 | 0.013 | 0.017 | 0.987 | 0.987 |
| FC | FPN | FPN | within | PPF2.7 | 0.454 | 0.324 | -0.786 | 0.032 | -4.085 | 0.000 | 0.003 |
| FC | FPN | DMN | between | LSD | 0.062 | 0.123 | 1.521 | 0.010 | 5.890 | 0.000 | 0.000 |
| FC | FPN | DMN | between | PSIL | 0.526 | 0.647 | 0.612 | 0.075 | 1.619 | 0.157 | 0.225 |
| FC | FPN | DMN | between | KTM | 0.229 | 0.266 | 0.813 | 0.013 | 2.815 | 0.017 | 0.038 |
| FC | FPN | DMN | between | N2O | 0.326 | 0.403 | 0.587 | 0.034 | 2.272 | 0.039 | 0.071 |
| FC | FPN | DMN | between | Sleep N1 | 0.413 | 0.393 | -0.241 | 0.014 | -1.386 | 0.175 | 0.225 |
| FC | FPN | DMN | between | Sleep N2 | 0.415 | 0.347 | -0.757 | 0.017 | -4.077 | 0.000 | 0.001 |
| FC | FPN | DMN | between | PPF1.9 | 0.354 | 0.359 | 0.028 | 0.044 | 0.103 | 0.919 | 0.919 |
| FC | FPN | DMN | between | PPF2.4 | 0.276 | 0.273 | -0.028 | 0.016 | -0.143 | 0.888 | 0.919 |
| FC | FPN | DMN | between | PPF2.7 | 0.383 | 0.232 | -0.914 | 0.032 | -4.747 | 0.000 | 0.000 |
| FC | DMN | DMN | within | LSD | 0.208 | 0.237 | 0.552 | 0.014 | 2.138 | 0.051 | 0.152 |
| FC | DMN | DMN | within | PSIL | 0.612 | 0.683 | 0.531 | 0.051 | 1.404 | 0.210 | 0.270 |
| FC | DMN | DMN | within | KTM | 0.295 | 0.312 | 0.399 | 0.012 | 1.384 | 0.194 | 0.270 |
| FC | DMN | DMN | within | N2O | 0.426 | 0.463 | 0.309 | 0.031 | 1.195 | 0.252 | 0.283 |
| FC | DMN | DMN | within | Sleep N1 | 0.501 | 0.484 | -0.255 | 0.011 | -1.467 | 0.152 | 0.270 |
| FC | DMN | DMN | within | Sleep N2 | 0.502 | 0.451 | -0.718 | 0.013 | -3.865 | 0.001 | 0.003 |
| FC | DMN | DMN | within | PPF1.9 | 0.459 | 0.426 | -0.254 | 0.034 | -0.950 | 0.359 | 0.359 |
| FC | DMN | DMN | within | PPF2.4 | 0.361 | 0.336 | -0.290 | 0.017 | -1.478 | 0.152 | 0.270 |
| FC | DMN | DMN | within | PPF2.7 | 0.481 | 0.325 | -1.050 | 0.029 | -5.453 | 0.000 | 0.000 |
| Efficiency | VIS | VIS | within | LSD | 0.780 | 0.828 | 1.053 | 0.012 | 4.080 | 0.001 | 0.005 |
| Efficiency | VIS | VIS | within | PSIL | 0.813 | 0.860 | 0.498 | 0.036 | 1.317 | 0.236 | 0.303 |
| Efficiency | VIS | VIS | within | KTM | 0.795 | 0.784 | -0.273 | 0.012 | -0.946 | 0.364 | 0.410 |
| Efficiency | VIS | VIS | within | N2O | 0.780 | 0.774 | -0.115 | 0.014 | -0.445 | 0.663 | 0.663 |
| Efficiency | VIS | VIS | within | Sleep N1 | 0.801 | 0.783 | -0.384 | 0.008 | -2.203 | 0.035 | 0.052 |
| Efficiency | VIS | VIS | within | Sleep N2 | 0.799 | 0.734 | -0.913 | 0.013 | -4.916 | 0.000 | 0.000 |
| Efficiency | VIS | VIS | within | PPF1.9 | 0.757 | 0.711 | -0.676 | 0.018 | -2.529 | 0.025 | 0.049 |
| Efficiency | VIS | VIS | within | PPF2.4 | 0.779 | 0.749 | -0.460 | 0.013 | -2.344 | 0.027 | 0.049 |
| Efficiency | VIS | VIS | within | PPF2.7 | 0.786 | 0.742 | -0.469 | 0.018 | -2.436 | 0.022 | 0.049 |
| Efficiency | VIS | SMN | between | LSD | 0.657 | 0.701 | 0.745 | 0.015 | 2.883 | 0.012 | 0.018 |
| Efficiency | VIS | SMN | between | PSIL | 0.727 | 0.790 | 0.700 | 0.034 | 1.853 | 0.113 | 0.146 |
| Efficiency | VIS | SMN | between | KTM | 0.720 | 0.722 | 0.065 | 0.008 | 0.225 | 0.826 | 0.910 |
| Efficiency | VIS | SMN | between | N2O | 0.684 | 0.684 | 0.030 | 0.008 | 0.115 | 0.910 | 0.910 |
| Efficiency | VIS | SMN | between | Sleep N1 | 0.667 | 0.641 | -0.605 | 0.007 | -3.477 | 0.001 | 0.003 |
| Efficiency | VIS | SMN | between | Sleep N2 | 0.668 | 0.597 | -1.262 | 0.010 | -6.798 | 0.000 | 0.000 |
| Efficiency | VIS | SMN | between | PPF1.9 | 0.637 | 0.549 | -1.625 | 0.014 | -6.079 | 0.000 | 0.000 |
| Efficiency | VIS | SMN | between | PPF2.4 | 0.680 | 0.604 | -1.075 | 0.014 | -5.483 | 0.000 | 0.000 |
| Efficiency | VIS | SMN | between | PPF2.7 | 0.675 | 0.596 | -0.668 | 0.023 | -3.469 | 0.002 | 0.003 |
| Efficiency | VIS | DAN | between | LSD | 0.696 | 0.757 | 1.772 | 0.009 | 6.861 | 0.000 | 0.000 |
| Efficiency | VIS | DAN | between | PSIL | 0.766 | 0.832 | 0.681 | 0.036 | 1.802 | 0.122 | 0.156 |
| Efficiency | VIS | DAN | between | KTM | 0.715 | 0.716 | 0.025 | 0.012 | 0.087 | 0.932 | 0.932 |
| Efficiency | VIS | DAN | between | N2O | 0.728 | 0.727 | -0.038 | 0.008 | -0.147 | 0.886 | 0.932 |
| Efficiency | VIS | DAN | between | Sleep N1 | 0.729 | 0.718 | -0.313 | 0.006 | -1.797 | 0.082 | 0.123 |
| Efficiency | VIS | DAN | between | Sleep N2 | 0.727 | 0.677 | -0.882 | 0.010 | -4.752 | 0.000 | 0.000 |
| Efficiency | VIS | DAN | between | PPF1.9 | 0.677 | 0.642 | -0.584 | 0.016 | -2.185 | 0.048 | 0.097 |
| Efficiency | VIS | DAN | between | PPF2.4 | 0.722 | 0.695 | -0.397 | 0.013 | -2.024 | 0.054 | 0.097 |
| Efficiency | VIS | DAN | between | PPF2.7 | 0.717 | 0.684 | -0.393 | 0.016 | -2.042 | 0.051 | 0.097 |
| Efficiency | VIS | VAN | between | LSD | 0.694 | 0.760 | 1.457 | 0.012 | 5.642 | 0.000 | 0.001 |
| Efficiency | VIS | VAN | between | PSIL | 0.756 | 0.822 | 0.630 | 0.040 | 1.667 | 0.146 | 0.330 |
| Efficiency | VIS | VAN | between | KTM | 0.709 | 0.708 | -0.014 | 0.017 | -0.049 | 0.962 | 0.962 |
| Efficiency | VIS | VAN | between | N2O | 0.722 | 0.716 | -0.110 | 0.016 | -0.424 | 0.678 | 0.762 |
| Efficiency | VIS | VAN | between | Sleep N1 | 0.675 | 0.663 | -0.280 | 0.007 | -1.607 | 0.118 | 0.330 |
| Efficiency | VIS | VAN | between | Sleep N2 | 0.670 | 0.638 | -0.550 | 0.011 | -2.960 | 0.006 | 0.028 |
| Efficiency | VIS | VAN | between | PPF1.9 | 0.635 | 0.614 | -0.334 | 0.017 | -1.249 | 0.234 | 0.372 |
| Efficiency | VIS | VAN | between | PPF2.4 | 0.681 | 0.664 | -0.212 | 0.016 | -1.082 | 0.289 | 0.372 |
| Efficiency | VIS | VAN | between | PPF2.7 | 0.667 | 0.647 | -0.214 | 0.018 | -1.111 | 0.277 | 0.372 |
| Efficiency | VIS | LIM | between | LSD | 0.647 | 0.725 | 1.265 | 0.016 | 4.900 | 0.000 | 0.002 |
| Efficiency | VIS | LIM | between | PSIL | 0.693 | 0.741 | 0.372 | 0.049 | 0.984 | 0.363 | 0.653 |
| Efficiency | VIS | LIM | between | KTM | 0.672 | 0.663 | -0.205 | 0.013 | -0.711 | 0.492 | 0.738 |
| Efficiency | VIS | LIM | between | N2O | 0.645 | 0.641 | -0.055 | 0.018 | -0.212 | 0.835 | 0.939 |
| Efficiency | VIS | LIM | between | Sleep N1 | 0.628 | 0.625 | -0.064 | 0.008 | -0.370 | 0.714 | 0.918 |
| Efficiency | VIS | LIM | between | Sleep N2 | 0.624 | 0.602 | -0.425 | 0.010 | -2.288 | 0.030 | 0.135 |
| Efficiency | VIS | LIM | between | PPF1.9 | 0.604 | 0.581 | -0.452 | 0.014 | -1.689 | 0.115 | 0.345 |
| Efficiency | VIS | LIM | between | PPF2.4 | 0.659 | 0.643 | -0.186 | 0.017 | -0.950 | 0.351 | 0.653 |
| Efficiency | VIS | LIM | between | PPF2.7 | 0.577 | 0.577 | 0.008 | 0.017 | 0.039 | 0.969 | 0.969 |
| Efficiency | VIS | FPN | between | LSD | 0.643 | 0.714 | 2.361 | 0.008 | 9.145 | 0.000 | 0.000 |
| Efficiency | VIS | FPN | between | PSIL | 0.748 | 0.809 | 0.537 | 0.043 | 1.422 | 0.205 | 0.369 |
| Efficiency | VIS | FPN | between | KTM | 0.669 | 0.684 | 0.263 | 0.016 | 0.909 | 0.383 | 0.527 |
| Efficiency | VIS | FPN | between | N2O | 0.679 | 0.699 | 0.428 | 0.012 | 1.656 | 0.120 | 0.360 |
| Efficiency | VIS | FPN | between | Sleep N1 | 0.649 | 0.645 | -0.100 | 0.007 | -0.572 | 0.571 | 0.571 |
| Efficiency | VIS | FPN | between | Sleep N2 | 0.645 | 0.626 | -0.320 | 0.011 | -1.725 | 0.095 | 0.360 |
| Efficiency | VIS | FPN | between | PPF1.9 | 0.611 | 0.622 | 0.177 | 0.017 | 0.663 | 0.519 | 0.571 |
| Efficiency | VIS | FPN | between | PPF2.4 | 0.647 | 0.661 | 0.164 | 0.017 | 0.838 | 0.410 | 0.527 |
| Efficiency | VIS | FPN | between | PPF2.7 | 0.617 | 0.645 | 0.276 | 0.020 | 1.434 | 0.164 | 0.368 |
| Efficiency | VIS | DMN | between | LSD | 0.659 | 0.719 | 2.035 | 0.008 | 7.882 | 0.000 | 0.000 |
| Efficiency | VIS | DMN | between | PSIL | 0.746 | 0.793 | 0.394 | 0.044 | 1.043 | 0.337 | 0.506 |
| Efficiency | VIS | DMN | between | KTM | 0.677 | 0.689 | 0.290 | 0.013 | 1.004 | 0.337 | 0.506 |
| Efficiency | VIS | DMN | between | N2O | 0.701 | 0.697 | -0.102 | 0.010 | -0.395 | 0.699 | 0.699 |
| Efficiency | VIS | DMN | between | Sleep N1 | 0.681 | 0.676 | -0.119 | 0.007 | -0.682 | 0.500 | 0.613 |
| Efficiency | VIS | DMN | between | Sleep N2 | 0.676 | 0.642 | -0.587 | 0.011 | -3.159 | 0.004 | 0.017 |
| Efficiency | VIS | DMN | between | PPF1.9 | 0.628 | 0.592 | -0.809 | 0.012 | -3.026 | 0.010 | 0.029 |
| Efficiency | VIS | DMN | between | PPF2.4 | 0.668 | 0.658 | -0.120 | 0.017 | -0.613 | 0.545 | 0.613 |
| Efficiency | VIS | DMN | between | PPF2.7 | 0.643 | 0.613 | -0.335 | 0.017 | -1.742 | 0.093 | 0.210 |
| Efficiency | SMN | SMN | within | LSD | 0.781 | 0.787 | 0.123 | 0.014 | 0.475 | 0.642 | 0.723 |
| Efficiency | SMN | SMN | within | PSIL | 0.810 | 0.863 | 1.175 | 0.017 | 3.109 | 0.021 | 0.031 |
| Efficiency | SMN | SMN | within | KTM | 0.832 | 0.838 | 0.219 | 0.009 | 0.760 | 0.463 | 0.596 |
| Efficiency | SMN | SMN | within | N2O | 0.783 | 0.783 | 0.006 | 0.009 | 0.022 | 0.983 | 0.983 |
| Efficiency | SMN | SMN | within | Sleep N1 | 0.770 | 0.750 | -0.700 | 0.005 | -4.020 | 0.000 | 0.001 |
| Efficiency | SMN | SMN | within | Sleep N2 | 0.770 | 0.716 | -0.899 | 0.011 | -4.840 | 0.000 | 0.000 |
| Efficiency | SMN | SMN | within | PPF1.9 | 0.760 | 0.682 | -1.525 | 0.014 | -5.704 | 0.000 | 0.000 |
| Efficiency | SMN | SMN | within | PPF2.4 | 0.755 | 0.717 | -0.687 | 0.011 | -3.505 | 0.002 | 0.004 |
| Efficiency | SMN | SMN | within | PPF2.7 | 0.767 | 0.704 | -0.633 | 0.019 | -3.289 | 0.003 | 0.005 |
| Efficiency | SMN | DAN | between | LSD | 0.644 | 0.694 | 0.809 | 0.016 | 3.133 | 0.007 | 0.017 |
| Efficiency | SMN | DAN | between | PSIL | 0.746 | 0.815 | 1.186 | 0.022 | 3.137 | 0.020 | 0.036 |
| Efficiency | SMN | DAN | between | KTM | 0.724 | 0.744 | 0.624 | 0.009 | 2.161 | 0.054 | 0.077 |
| Efficiency | SMN | DAN | between | N2O | 0.717 | 0.724 | 0.159 | 0.012 | 0.615 | 0.548 | 0.548 |
| Efficiency | SMN | DAN | between | Sleep N1 | 0.688 | 0.666 | -0.642 | 0.006 | -3.686 | 0.001 | 0.003 |
| Efficiency | SMN | DAN | between | Sleep N2 | 0.692 | 0.624 | -1.016 | 0.012 | -5.470 | 0.000 | 0.000 |
| Efficiency | SMN | DAN | between | PPF1.9 | 0.661 | 0.599 | -1.283 | 0.013 | -4.802 | 0.000 | 0.002 |
| Efficiency | SMN | DAN | between | PPF2.4 | 0.659 | 0.641 | -0.286 | 0.013 | -1.459 | 0.157 | 0.177 |
| Efficiency | SMN | DAN | between | PPF2.7 | 0.675 | 0.634 | -0.379 | 0.021 | -1.968 | 0.060 | 0.077 |
| Efficiency | SMN | VAN | between | LSD | 0.685 | 0.725 | 0.667 | 0.015 | 2.582 | 0.022 | 0.049 |
| Efficiency | SMN | VAN | between | PSIL | 0.757 | 0.828 | 2.039 | 0.013 | 5.394 | 0.002 | 0.005 |
| Efficiency | SMN | VAN | between | KTM | 0.739 | 0.757 | 0.480 | 0.010 | 1.664 | 0.124 | 0.160 |
| Efficiency | SMN | VAN | between | N2O | 0.732 | 0.739 | 0.205 | 0.010 | 0.792 | 0.441 | 0.497 |
| Efficiency | SMN | VAN | between | Sleep N1 | 0.686 | 0.671 | -0.399 | 0.007 | -2.294 | 0.028 | 0.051 |
| Efficiency | SMN | VAN | between | Sleep N2 | 0.687 | 0.642 | -0.904 | 0.009 | -4.868 | 0.000 | 0.000 |
| Efficiency | SMN | VAN | between | PPF1.9 | 0.665 | 0.607 | -1.208 | 0.013 | -4.519 | 0.001 | 0.003 |
| Efficiency | SMN | VAN | between | PPF2.4 | 0.673 | 0.667 | -0.113 | 0.012 | -0.574 | 0.571 | 0.571 |
| Efficiency | SMN | VAN | between | PPF2.7 | 0.685 | 0.650 | -0.338 | 0.020 | -1.759 | 0.090 | 0.136 |
| Efficiency | SMN | LIM | between | LSD | 0.521 | 0.556 | 0.529 | 0.017 | 2.049 | 0.060 | 0.134 |
| Efficiency | SMN | LIM | between | PSIL | 0.624 | 0.659 | 0.265 | 0.050 | 0.702 | 0.509 | 0.764 |
| Efficiency | SMN | LIM | between | KTM | 0.639 | 0.669 | 0.674 | 0.013 | 2.336 | 0.039 | 0.118 |
| Efficiency | SMN | LIM | between | N2O | 0.595 | 0.614 | 0.301 | 0.016 | 1.166 | 0.263 | 0.474 |
| Efficiency | SMN | LIM | between | Sleep N1 | 0.577 | 0.577 | 0.020 | 0.006 | 0.113 | 0.911 | 0.911 |
| Efficiency | SMN | LIM | between | Sleep N2 | 0.579 | 0.558 | -0.457 | 0.008 | -2.460 | 0.020 | 0.091 |
| Efficiency | SMN | LIM | between | PPF1.9 | 0.586 | 0.522 | -2.035 | 0.008 | -7.613 | 0.000 | 0.000 |
| Efficiency | SMN | LIM | between | PPF2.4 | 0.597 | 0.592 | -0.070 | 0.014 | -0.357 | 0.724 | 0.814 |
| Efficiency | SMN | LIM | between | PPF2.7 | 0.534 | 0.541 | 0.069 | 0.021 | 0.357 | 0.724 | 0.814 |
| Efficiency | SMN | FPN | between | LSD | 0.538 | 0.598 | 1.020 | 0.015 | 3.951 | 0.001 | 0.006 |
| Efficiency | SMN | FPN | between | PSIL | 0.721 | 0.789 | 0.872 | 0.030 | 2.308 | 0.060 | 0.109 |
| Efficiency | SMN | FPN | between | KTM | 0.677 | 0.709 | 0.962 | 0.010 | 3.332 | 0.007 | 0.015 |
| Efficiency | SMN | FPN | between | N2O | 0.654 | 0.699 | 0.989 | 0.012 | 3.831 | 0.002 | 0.006 |
| Efficiency | SMN | FPN | between | Sleep N1 | 0.603 | 0.590 | -0.290 | 0.008 | -1.667 | 0.105 | 0.158 |
| Efficiency | SMN | FPN | between | Sleep N2 | 0.608 | 0.562 | -0.773 | 0.011 | -4.164 | 0.000 | 0.002 |
| Efficiency | SMN | FPN | between | PPF1.9 | 0.578 | 0.562 | -0.270 | 0.015 | -1.012 | 0.330 | 0.371 |
| Efficiency | SMN | FPN | between | PPF2.4 | 0.573 | 0.597 | 0.302 | 0.016 | 1.542 | 0.136 | 0.174 |
| Efficiency | SMN | FPN | between | PPF2.7 | 0.569 | 0.584 | 0.132 | 0.023 | 0.684 | 0.500 | 0.500 |
| Efficiency | SMN | DMN | between | LSD | 0.564 | 0.625 | 1.046 | 0.015 | 4.049 | 0.001 | 0.005 |
| Efficiency | SMN | DMN | between | PSIL | 0.742 | 0.789 | 0.536 | 0.033 | 1.419 | 0.206 | 0.309 |
| Efficiency | SMN | DMN | between | KTM | 0.675 | 0.703 | 0.749 | 0.011 | 2.596 | 0.025 | 0.056 |
| Efficiency | SMN | DMN | between | N2O | 0.683 | 0.706 | 0.506 | 0.012 | 1.960 | 0.070 | 0.126 |
| Efficiency | SMN | DMN | between | Sleep N1 | 0.645 | 0.640 | -0.186 | 0.005 | -1.070 | 0.293 | 0.376 |
| Efficiency | SMN | DMN | between | Sleep N2 | 0.647 | 0.614 | -0.740 | 0.008 | -3.987 | 0.000 | 0.004 |
| Efficiency | SMN | DMN | between | PPF1.9 | 0.598 | 0.567 | -0.746 | 0.011 | -2.792 | 0.015 | 0.046 |
| Efficiency | SMN | DMN | between | PPF2.4 | 0.612 | 0.616 | 0.051 | 0.015 | 0.260 | 0.797 | 0.797 |
| Efficiency | SMN | DMN | between | PPF2.7 | 0.606 | 0.589 | -0.178 | 0.019 | -0.927 | 0.362 | 0.408 |
| Efficiency | DAN | DAN | within | LSD | 0.792 | 0.835 | 1.070 | 0.010 | 4.144 | 0.001 | 0.002 |
| Efficiency | DAN | DAN | within | PSIL | 0.865 | 0.910 | 1.193 | 0.014 | 3.156 | 0.020 | 0.025 |
| Efficiency | DAN | DAN | within | KTM | 0.837 | 0.842 | 0.117 | 0.013 | 0.407 | 0.692 | 0.692 |
| Efficiency | DAN | DAN | within | N2O | 0.844 | 0.839 | -0.170 | 0.007 | -0.657 | 0.522 | 0.587 |
| Efficiency | DAN | DAN | within | Sleep N1 | 0.851 | 0.833 | -0.608 | 0.005 | -3.493 | 0.001 | 0.003 |
| Efficiency | DAN | DAN | within | Sleep N2 | 0.850 | 0.820 | -0.698 | 0.008 | -3.760 | 0.001 | 0.002 |
| Efficiency | DAN | DAN | within | PPF1.9 | 0.827 | 0.757 | -1.405 | 0.013 | -5.258 | 0.000 | 0.001 |
| Efficiency | DAN | DAN | within | PPF2.4 | 0.831 | 0.798 | -0.606 | 0.011 | -3.092 | 0.005 | 0.007 |
| Efficiency | DAN | DAN | within | PPF2.7 | 0.860 | 0.802 | -0.910 | 0.012 | -4.729 | 0.000 | 0.001 |
| Efficiency | DAN | VAN | between | LSD | 0.709 | 0.748 | 0.919 | 0.011 | 3.559 | 0.003 | 0.006 |
| Efficiency | DAN | VAN | between | PSIL | 0.815 | 0.852 | 0.744 | 0.019 | 1.967 | 0.097 | 0.124 |
| Efficiency | DAN | VAN | between | KTM | 0.747 | 0.768 | 0.474 | 0.013 | 1.640 | 0.129 | 0.145 |
| Efficiency | DAN | VAN | between | N2O | 0.787 | 0.775 | -0.347 | 0.009 | -1.345 | 0.200 | 0.200 |
| Efficiency | DAN | VAN | between | Sleep N1 | 0.775 | 0.764 | -0.395 | 0.005 | -2.267 | 0.030 | 0.045 |
| Efficiency | DAN | VAN | between | Sleep N2 | 0.773 | 0.745 | -0.709 | 0.007 | -3.818 | 0.001 | 0.002 |
| Efficiency | DAN | VAN | between | PPF1.9 | 0.704 | 0.645 | -1.159 | 0.014 | -4.335 | 0.001 | 0.002 |
| Efficiency | DAN | VAN | between | PPF2.4 | 0.762 | 0.726 | -0.649 | 0.011 | -3.309 | 0.003 | 0.006 |
| Efficiency | DAN | VAN | between | PPF2.7 | 0.780 | 0.728 | -0.846 | 0.012 | -4.396 | 0.000 | 0.001 |
| Efficiency | DAN | LIM | between | LSD | 0.647 | 0.689 | 0.545 | 0.020 | 2.110 | 0.053 | 0.160 |
| Efficiency | DAN | LIM | between | PSIL | 0.744 | 0.742 | -0.014 | 0.042 | -0.037 | 0.972 | 0.972 |
| Efficiency | DAN | LIM | between | KTM | 0.742 | 0.749 | 0.172 | 0.012 | 0.596 | 0.563 | 0.724 |
| Efficiency | DAN | LIM | between | N2O | 0.722 | 0.702 | -0.291 | 0.017 | -1.127 | 0.279 | 0.493 |
| Efficiency | DAN | LIM | between | Sleep N1 | 0.742 | 0.742 | -0.006 | 0.006 | -0.037 | 0.971 | 0.972 |
| Efficiency | DAN | LIM | between | Sleep N2 | 0.738 | 0.748 | 0.251 | 0.008 | 1.350 | 0.188 | 0.423 |
| Efficiency | DAN | LIM | between | PPF1.9 | 0.714 | 0.613 | -1.000 | 0.027 | -3.743 | 0.002 | 0.022 |
| Efficiency | DAN | LIM | between | PPF2.4 | 0.757 | 0.710 | -0.529 | 0.017 | -2.696 | 0.012 | 0.056 |
| Efficiency | DAN | LIM | between | PPF2.7 | 0.692 | 0.674 | -0.192 | 0.017 | -0.996 | 0.329 | 0.493 |
| Efficiency | DAN | FPN | between | LSD | 0.673 | 0.692 | 0.371 | 0.013 | 1.437 | 0.173 | 0.518 |
| Efficiency | DAN | FPN | between | PSIL | 0.791 | 0.816 | 0.421 | 0.022 | 1.114 | 0.308 | 0.554 |
| Efficiency | DAN | FPN | between | KTM | 0.725 | 0.728 | 0.073 | 0.013 | 0.252 | 0.806 | 0.907 |
| Efficiency | DAN | FPN | between | N2O | 0.739 | 0.736 | -0.077 | 0.011 | -0.297 | 0.771 | 0.907 |
| Efficiency | DAN | FPN | between | Sleep N1 | 0.752 | 0.749 | -0.123 | 0.005 | -0.708 | 0.484 | 0.726 |
| Efficiency | DAN | FPN | between | Sleep N2 | 0.752 | 0.752 | -0.006 | 0.006 | -0.034 | 0.973 | 0.973 |
| Efficiency | DAN | FPN | between | PPF1.9 | 0.685 | 0.647 | -0.625 | 0.016 | -2.340 | 0.036 | 0.323 |
| Efficiency | DAN | FPN | between | PPF2.4 | 0.728 | 0.713 | -0.218 | 0.013 | -1.112 | 0.277 | 0.554 |
| Efficiency | DAN | FPN | between | PPF2.7 | 0.740 | 0.723 | -0.279 | 0.012 | -1.448 | 0.160 | 0.518 |
| Efficiency | DAN | DMN | between | LSD | 0.627 | 0.678 | 1.223 | 0.011 | 4.735 | 0.000 | 0.002 |
| Efficiency | DAN | DMN | between | PSIL | 0.766 | 0.803 | 0.336 | 0.042 | 0.888 | 0.409 | 0.526 |
| Efficiency | DAN | DMN | between | KTM | 0.694 | 0.721 | 0.488 | 0.016 | 1.689 | 0.119 | 0.215 |
| Efficiency | DAN | DMN | between | N2O | 0.706 | 0.714 | 0.161 | 0.012 | 0.622 | 0.544 | 0.612 |
| Efficiency | DAN | DMN | between | Sleep N1 | 0.721 | 0.719 | -0.058 | 0.006 | -0.332 | 0.742 | 0.742 |
| Efficiency | DAN | DMN | between | Sleep N2 | 0.719 | 0.699 | -0.583 | 0.007 | -3.140 | 0.004 | 0.009 |
| Efficiency | DAN | DMN | between | PPF1.9 | 0.679 | 0.613 | -1.238 | 0.014 | -4.631 | 0.000 | 0.002 |
| Efficiency | DAN | DMN | between | PPF2.4 | 0.713 | 0.697 | -0.266 | 0.011 | -1.357 | 0.187 | 0.280 |
| Efficiency | DAN | DMN | between | PPF2.7 | 0.707 | 0.644 | -0.669 | 0.018 | -3.477 | 0.002 | 0.005 |
| Efficiency | VAN | VAN | within | LSD | 0.807 | 0.837 | 0.753 | 0.010 | 2.916 | 0.011 | 0.020 |
| Efficiency | VAN | VAN | within | PSIL | 0.878 | 0.908 | 1.315 | 0.009 | 3.480 | 0.013 | 0.020 |
| Efficiency | VAN | VAN | within | KTM | 0.830 | 0.863 | 0.947 | 0.010 | 3.279 | 0.007 | 0.020 |
| Efficiency | VAN | VAN | within | N2O | 0.859 | 0.863 | 0.108 | 0.009 | 0.418 | 0.682 | 0.682 |
| Efficiency | VAN | VAN | within | Sleep N1 | 0.830 | 0.825 | -0.153 | 0.006 | -0.880 | 0.386 | 0.434 |
| Efficiency | VAN | VAN | within | Sleep N2 | 0.829 | 0.807 | -0.563 | 0.007 | -3.030 | 0.005 | 0.020 |
| Efficiency | VAN | VAN | within | PPF1.9 | 0.824 | 0.783 | -0.792 | 0.014 | -2.963 | 0.011 | 0.020 |
| Efficiency | VAN | VAN | within | PPF2.4 | 0.852 | 0.845 | -0.250 | 0.005 | -1.277 | 0.213 | 0.274 |
| Efficiency | VAN | VAN | within | PPF2.7 | 0.859 | 0.835 | -0.527 | 0.009 | -2.740 | 0.011 | 0.020 |
| Efficiency | VAN | LIM | between | LSD | 0.660 | 0.685 | 0.374 | 0.018 | 1.448 | 0.170 | 0.258 |
| Efficiency | VAN | LIM | between | PSIL | 0.767 | 0.728 | -0.743 | 0.020 | -1.965 | 0.097 | 0.218 |
| Efficiency | VAN | LIM | between | KTM | 0.726 | 0.766 | 0.904 | 0.013 | 3.131 | 0.010 | 0.043 |
| Efficiency | VAN | LIM | between | N2O | 0.717 | 0.719 | 0.040 | 0.014 | 0.156 | 0.878 | 0.878 |
| Efficiency | VAN | LIM | between | Sleep N1 | 0.731 | 0.739 | 0.243 | 0.005 | 1.397 | 0.172 | 0.258 |
| Efficiency | VAN | LIM | between | Sleep N2 | 0.733 | 0.732 | -0.034 | 0.007 | -0.186 | 0.854 | 0.878 |
| Efficiency | VAN | LIM | between | PPF1.9 | 0.734 | 0.680 | -0.632 | 0.023 | -2.365 | 0.034 | 0.103 |
| Efficiency | VAN | LIM | between | PPF2.4 | 0.766 | 0.773 | 0.155 | 0.010 | 0.790 | 0.437 | 0.562 |
| Efficiency | VAN | LIM | between | PPF2.7 | 0.686 | 0.738 | 0.610 | 0.017 | 3.168 | 0.004 | 0.035 |
| Efficiency | VAN | FPN | between | LSD | 0.692 | 0.724 | 0.656 | 0.012 | 2.539 | 0.024 | 0.054 |
| Efficiency | VAN | FPN | between | PSIL | 0.811 | 0.822 | 0.194 | 0.020 | 0.514 | 0.626 | 0.626 |
| Efficiency | VAN | FPN | between | KTM | 0.756 | 0.783 | 0.794 | 0.010 | 2.750 | 0.019 | 0.054 |
| Efficiency | VAN | FPN | between | N2O | 0.779 | 0.770 | -0.214 | 0.011 | -0.830 | 0.420 | 0.626 |
| Efficiency | VAN | FPN | between | Sleep N1 | 0.772 | 0.775 | 0.105 | 0.004 | 0.604 | 0.550 | 0.626 |
| Efficiency | VAN | FPN | between | Sleep N2 | 0.772 | 0.759 | -0.443 | 0.005 | -2.388 | 0.024 | 0.054 |
| Efficiency | VAN | FPN | between | PPF1.9 | 0.715 | 0.677 | -0.967 | 0.010 | -3.617 | 0.003 | 0.028 |
| Efficiency | VAN | FPN | between | PPF2.4 | 0.770 | 0.764 | -0.115 | 0.010 | -0.589 | 0.561 | 0.626 |
| Efficiency | VAN | FPN | between | PPF2.7 | 0.759 | 0.745 | -0.256 | 0.011 | -1.332 | 0.195 | 0.350 |
| Efficiency | VAN | DMN | between | LSD | 0.660 | 0.700 | 0.989 | 0.011 | 3.830 | 0.002 | 0.007 |
| Efficiency | VAN | DMN | between | PSIL | 0.783 | 0.801 | 0.199 | 0.035 | 0.527 | 0.617 | 0.617 |
| Efficiency | VAN | DMN | between | KTM | 0.706 | 0.735 | 0.758 | 0.011 | 2.626 | 0.024 | 0.053 |
| Efficiency | VAN | DMN | between | N2O | 0.734 | 0.723 | -0.264 | 0.011 | -1.022 | 0.324 | 0.486 |
| Efficiency | VAN | DMN | between | Sleep N1 | 0.723 | 0.730 | 0.239 | 0.005 | 1.373 | 0.179 | 0.323 |
| Efficiency | VAN | DMN | between | Sleep N2 | 0.720 | 0.714 | -0.162 | 0.007 | -0.870 | 0.392 | 0.504 |
| Efficiency | VAN | DMN | between | PPF1.9 | 0.693 | 0.635 | -1.004 | 0.015 | -3.756 | 0.002 | 0.007 |
| Efficiency | VAN | DMN | between | PPF2.4 | 0.739 | 0.734 | -0.133 | 0.008 | -0.679 | 0.503 | 0.566 |
| Efficiency | VAN | DMN | between | PPF2.7 | 0.718 | 0.662 | -0.803 | 0.013 | -4.172 | 0.000 | 0.003 |
| Efficiency | LIM | LIM | within | LSD | 0.893 | 0.894 | 0.030 | 0.011 | 0.117 | 0.908 | 0.908 |
| Efficiency | LIM | LIM | within | PSIL | 0.955 | 0.949 | -0.244 | 0.008 | -0.645 | 0.543 | 0.682 |
| Efficiency | LIM | LIM | within | KTM | 0.849 | 0.840 | -0.191 | 0.013 | -0.663 | 0.521 | 0.682 |
| Efficiency | LIM | LIM | within | N2O | 0.894 | 0.872 | -0.493 | 0.011 | -1.910 | 0.077 | 0.346 |
| Efficiency | LIM | LIM | within | Sleep N1 | 0.862 | 0.858 | -0.091 | 0.008 | -0.521 | 0.606 | 0.682 |
| Efficiency | LIM | LIM | within | Sleep N2 | 0.863 | 0.850 | -0.290 | 0.008 | -1.562 | 0.130 | 0.389 |
| Efficiency | LIM | LIM | within | PPF1.9 | 0.872 | 0.844 | -0.278 | 0.027 | -1.040 | 0.317 | 0.571 |
| Efficiency | LIM | LIM | within | PPF2.4 | 0.883 | 0.872 | -0.213 | 0.011 | -1.089 | 0.287 | 0.571 |
| Efficiency | LIM | LIM | within | PPF2.7 | 0.825 | 0.882 | 0.583 | 0.019 | 3.030 | 0.005 | 0.049 |
| Efficiency | LIM | FPN | between | LSD | 0.681 | 0.704 | 0.352 | 0.017 | 1.362 | 0.195 | 0.292 |
| Efficiency | LIM | FPN | between | PSIL | 0.777 | 0.722 | -0.644 | 0.032 | -1.704 | 0.139 | 0.251 |
| Efficiency | LIM | FPN | between | KTM | 0.740 | 0.748 | 0.195 | 0.011 | 0.675 | 0.514 | 0.578 |
| Efficiency | LIM | FPN | between | N2O | 0.728 | 0.715 | -0.244 | 0.014 | -0.946 | 0.360 | 0.463 |
| Efficiency | LIM | FPN | between | Sleep N1 | 0.757 | 0.772 | 0.498 | 0.005 | 2.859 | 0.007 | 0.022 |
| Efficiency | LIM | FPN | between | Sleep N2 | 0.757 | 0.773 | 0.507 | 0.006 | 2.730 | 0.011 | 0.024 |
| Efficiency | LIM | FPN | between | PPF1.9 | 0.734 | 0.641 | -0.933 | 0.027 | -3.491 | 0.004 | 0.018 |
| Efficiency | LIM | FPN | between | PPF2.4 | 0.788 | 0.751 | -0.663 | 0.011 | -3.382 | 0.002 | 0.018 |
| Efficiency | LIM | FPN | between | PPF2.7 | 0.706 | 0.702 | -0.038 | 0.016 | -0.197 | 0.845 | 0.845 |
| Efficiency | LIM | DMN | between | LSD | 0.641 | 0.677 | 0.616 | 0.015 | 2.387 | 0.032 | 0.063 |
| Efficiency | LIM | DMN | between | PSIL | 0.748 | 0.694 | -0.389 | 0.052 | -1.029 | 0.343 | 0.386 |
| Efficiency | LIM | DMN | between | KTM | 0.663 | 0.687 | 0.425 | 0.016 | 1.472 | 0.169 | 0.217 |
| Efficiency | LIM | DMN | between | N2O | 0.688 | 0.656 | -0.529 | 0.016 | -2.047 | 0.060 | 0.090 |
| Efficiency | LIM | DMN | between | Sleep N1 | 0.681 | 0.698 | 0.452 | 0.007 | 2.598 | 0.014 | 0.042 |
| Efficiency | LIM | DMN | between | Sleep N2 | 0.676 | 0.680 | 0.074 | 0.009 | 0.399 | 0.693 | 0.693 |
| Efficiency | LIM | DMN | between | PPF1.9 | 0.681 | 0.567 | -1.135 | 0.027 | -4.249 | 0.001 | 0.006 |
| Efficiency | LIM | DMN | between | PPF2.4 | 0.743 | 0.701 | -0.706 | 0.012 | -3.601 | 0.001 | 0.006 |
| Efficiency | LIM | DMN | between | PPF2.7 | 0.651 | 0.611 | -0.428 | 0.018 | -2.226 | 0.035 | 0.063 |
| Efficiency | FPN | FPN | within | LSD | 0.812 | 0.835 | 0.604 | 0.010 | 2.338 | 0.035 | 0.078 |
| Efficiency | FPN | FPN | within | PSIL | 0.872 | 0.883 | 0.226 | 0.019 | 0.598 | 0.572 | 0.643 |
| Efficiency | FPN | FPN | within | KTM | 0.837 | 0.852 | 0.458 | 0.009 | 1.588 | 0.141 | 0.253 |
| Efficiency | FPN | FPN | within | N2O | 0.828 | 0.815 | -0.347 | 0.010 | -1.345 | 0.200 | 0.300 |
| Efficiency | FPN | FPN | within | Sleep N1 | 0.851 | 0.849 | -0.033 | 0.006 | -0.191 | 0.850 | 0.850 |
| Efficiency | FPN | FPN | within | Sleep N2 | 0.850 | 0.845 | -0.164 | 0.006 | -0.883 | 0.385 | 0.495 |
| Efficiency | FPN | FPN | within | PPF1.9 | 0.832 | 0.776 | -1.052 | 0.014 | -3.936 | 0.002 | 0.015 |
| Efficiency | FPN | FPN | within | PPF2.4 | 0.848 | 0.824 | -0.540 | 0.009 | -2.755 | 0.011 | 0.032 |
| Efficiency | FPN | FPN | within | PPF2.7 | 0.837 | 0.810 | -0.596 | 0.009 | -3.098 | 0.005 | 0.021 |
| Efficiency | FPN | DMN | between | LSD | 0.688 | 0.722 | 0.747 | 0.012 | 2.892 | 0.012 | 0.035 |
| Efficiency | FPN | DMN | between | PSIL | 0.789 | 0.794 | 0.067 | 0.031 | 0.177 | 0.865 | 0.865 |
| Efficiency | FPN | DMN | between | KTM | 0.721 | 0.741 | 0.435 | 0.013 | 1.505 | 0.160 | 0.291 |
| Efficiency | FPN | DMN | between | N2O | 0.731 | 0.727 | -0.084 | 0.011 | -0.324 | 0.751 | 0.844 |
| Efficiency | FPN | DMN | between | Sleep N1 | 0.739 | 0.743 | 0.097 | 0.006 | 0.557 | 0.581 | 0.747 |
| Efficiency | FPN | DMN | between | Sleep N2 | 0.737 | 0.725 | -0.267 | 0.009 | -1.438 | 0.162 | 0.291 |
| Efficiency | FPN | DMN | between | PPF1.9 | 0.700 | 0.646 | -0.865 | 0.017 | -3.235 | 0.007 | 0.029 |
| Efficiency | FPN | DMN | between | PPF2.4 | 0.749 | 0.742 | -0.157 | 0.009 | -0.798 | 0.432 | 0.648 |
| Efficiency | FPN | DMN | between | PPF2.7 | 0.726 | 0.679 | -0.669 | 0.013 | -3.476 | 0.002 | 0.016 |
| Efficiency | DMN | DMN | within | LSD | 0.767 | 0.805 | 1.322 | 0.007 | 5.119 | 0.000 | 0.001 |
| Efficiency | DMN | DMN | within | PSIL | 0.847 | 0.859 | 0.167 | 0.026 | 0.443 | 0.673 | 0.758 |
| Efficiency | DMN | DMN | within | KTM | 0.789 | 0.804 | 0.383 | 0.011 | 1.328 | 0.211 | 0.363 |
| Efficiency | DMN | DMN | within | N2O | 0.795 | 0.781 | -0.316 | 0.011 | -1.222 | 0.242 | 0.363 |
| Efficiency | DMN | DMN | within | Sleep N1 | 0.804 | 0.801 | -0.080 | 0.006 | -0.458 | 0.650 | 0.758 |
| Efficiency | DMN | DMN | within | Sleep N2 | 0.800 | 0.782 | -0.436 | 0.008 | -2.348 | 0.026 | 0.059 |
| Efficiency | DMN | DMN | within | PPF1.9 | 0.784 | 0.718 | -1.016 | 0.017 | -3.801 | 0.002 | 0.007 |
| Efficiency | DMN | DMN | within | PPF2.4 | 0.786 | 0.785 | -0.011 | 0.006 | -0.058 | 0.954 | 0.954 |
| Efficiency | DMN | DMN | within | PPF2.7 | 0.789 | 0.736 | -0.880 | 0.012 | -4.575 | 0.000 | 0.001 |
| Complexity | VIS | VIS | within | LSD | 0.240 | 0.242 | 0.304 | 0.002 | 1.176 | 0.259 | 0.291 |
| Complexity | VIS | VIS | within | PSIL | 0.253 | 0.256 | 0.319 | 0.004 | 0.844 | 0.431 | 0.431 |
| Complexity | VIS | VIS | within | KTM | 0.291 | 0.281 | -0.973 | 0.003 | -3.371 | 0.006 | 0.011 |
| Complexity | VIS | VIS | within | N2O | 0.270 | 0.265 | -0.361 | 0.004 | -1.397 | 0.184 | 0.237 |
| Complexity | VIS | VIS | within | Sleep N1 | 0.268 | 0.266 | -0.240 | 0.001 | -1.379 | 0.177 | 0.237 |
| Complexity | VIS | VIS | within | Sleep N2 | 0.268 | 0.260 | -0.841 | 0.002 | -4.529 | 0.000 | 0.000 |
| Complexity | VIS | VIS | within | PPF1.9 | 0.274 | 0.266 | -0.994 | 0.002 | -3.718 | 0.003 | 0.008 |
| Complexity | VIS | VIS | within | PPF2.4 | 0.179 | 0.165 | -1.461 | 0.002 | -7.451 | 0.000 | 0.000 |
| Complexity | VIS | VIS | within | PPF2.7 | 0.228 | 0.219 | -0.622 | 0.003 | -3.229 | 0.003 | 0.008 |
| Complexity | VIS | SMN | between | LSD | 0.227 | 0.232 | 0.427 | 0.003 | 1.653 | 0.121 | 0.181 |
| Complexity | VIS | SMN | between | PSIL | 0.244 | 0.250 | 0.535 | 0.004 | 1.414 | 0.207 | 0.266 |
| Complexity | VIS | SMN | between | KTM | 0.280 | 0.278 | -0.111 | 0.003 | -0.384 | 0.708 | 0.708 |
| Complexity | VIS | SMN | between | N2O | 0.258 | 0.254 | -0.238 | 0.004 | -0.921 | 0.373 | 0.419 |
| Complexity | VIS | SMN | between | Sleep N1 | 0.257 | 0.254 | -0.388 | 0.001 | -2.231 | 0.033 | 0.059 |
| Complexity | VIS | SMN | between | Sleep N2 | 0.257 | 0.249 | -0.790 | 0.002 | -4.255 | 0.000 | 0.001 |
| Complexity | VIS | SMN | between | PPF1.9 | 0.265 | 0.251 | -1.293 | 0.003 | -4.839 | 0.000 | 0.001 |
| Complexity | VIS | SMN | between | PPF2.4 | 0.167 | 0.149 | -1.733 | 0.002 | -8.839 | 0.000 | 0.000 |
| Complexity | VIS | SMN | between | PPF2.7 | 0.216 | 0.201 | -0.971 | 0.003 | -5.044 | 0.000 | 0.000 |
| Complexity | VIS | DAN | between | LSD | 0.236 | 0.246 | 0.981 | 0.003 | 3.798 | 0.002 | 0.004 |
| Complexity | VIS | DAN | between | PSIL | 0.260 | 0.267 | 0.466 | 0.006 | 1.232 | 0.264 | 0.339 |
| Complexity | VIS | DAN | between | KTM | 0.294 | 0.294 | 0.018 | 0.004 | 0.061 | 0.952 | 0.952 |
| Complexity | VIS | DAN | between | N2O | 0.278 | 0.274 | -0.232 | 0.004 | -0.897 | 0.385 | 0.433 |
| Complexity | VIS | DAN | between | Sleep N1 | 0.276 | 0.274 | -0.210 | 0.002 | -1.208 | 0.236 | 0.339 |
| Complexity | VIS | DAN | between | Sleep N2 | 0.276 | 0.266 | -0.689 | 0.003 | -3.713 | 0.001 | 0.003 |
| Complexity | VIS | DAN | between | PPF1.9 | 0.280 | 0.267 | -1.102 | 0.003 | -4.122 | 0.001 | 0.003 |
| Complexity | VIS | DAN | between | PPF2.4 | 0.174 | 0.159 | -1.331 | 0.002 | -6.785 | 0.000 | 0.000 |
| Complexity | VIS | DAN | between | PPF2.7 | 0.228 | 0.217 | -0.817 | 0.003 | -4.247 | 0.000 | 0.001 |
| Complexity | VIS | VAN | between | LSD | 0.239 | 0.245 | 0.658 | 0.002 | 2.548 | 0.023 | 0.042 |
| Complexity | VIS | VAN | between | PSIL | 0.258 | 0.265 | 0.499 | 0.006 | 1.320 | 0.235 | 0.302 |
| Complexity | VIS | VAN | between | KTM | 0.293 | 0.291 | -0.256 | 0.003 | -0.887 | 0.394 | 0.444 |
| Complexity | VIS | VAN | between | N2O | 0.277 | 0.274 | -0.189 | 0.005 | -0.733 | 0.476 | 0.476 |
| Complexity | VIS | VAN | between | Sleep N1 | 0.272 | 0.270 | -0.261 | 0.002 | -1.501 | 0.143 | 0.215 |
| Complexity | VIS | VAN | between | Sleep N2 | 0.272 | 0.265 | -0.601 | 0.002 | -3.238 | 0.003 | 0.007 |
| Complexity | VIS | VAN | between | PPF1.9 | 0.280 | 0.269 | -1.011 | 0.003 | -3.783 | 0.002 | 0.007 |
| Complexity | VIS | VAN | between | PPF2.4 | 0.169 | 0.156 | -1.272 | 0.002 | -6.485 | 0.000 | 0.000 |
| Complexity | VIS | VAN | between | PPF2.7 | 0.223 | 0.211 | -1.131 | 0.002 | -5.875 | 0.000 | 0.000 |
| Complexity | VIS | LIM | between | LSD | 0.280 | 0.284 | 0.291 | 0.003 | 1.126 | 0.279 | 0.365 |
| Complexity | VIS | LIM | between | PSIL | 0.306 | 0.313 | 0.391 | 0.007 | 1.034 | 0.341 | 0.384 |
| Complexity | VIS | LIM | between | KTM | 0.343 | 0.339 | -0.390 | 0.003 | -1.352 | 0.204 | 0.365 |
| Complexity | VIS | LIM | between | N2O | 0.322 | 0.314 | -0.401 | 0.005 | -1.553 | 0.143 | 0.321 |
| Complexity | VIS | LIM | between | Sleep N1 | 0.319 | 0.321 | 0.138 | 0.002 | 0.792 | 0.434 | 0.434 |
| Complexity | VIS | LIM | between | Sleep N2 | 0.319 | 0.316 | -0.203 | 0.002 | -1.093 | 0.284 | 0.365 |
| Complexity | VIS | LIM | between | PPF1.9 | 0.330 | 0.317 | -0.566 | 0.006 | -2.119 | 0.054 | 0.162 |
| Complexity | VIS | LIM | between | PPF2.4 | 0.199 | 0.183 | -1.568 | 0.002 | -7.995 | 0.000 | 0.000 |
| Complexity | VIS | LIM | between | PPF2.7 | 0.257 | 0.250 | -0.430 | 0.003 | -2.232 | 0.034 | 0.155 |
| Complexity | VIS | FPN | between | LSD | 0.225 | 0.232 | 0.680 | 0.003 | 2.632 | 0.020 | 0.059 |
| Complexity | VIS | FPN | between | PSIL | 0.250 | 0.259 | 0.549 | 0.006 | 1.454 | 0.196 | 0.294 |
| Complexity | VIS | FPN | between | KTM | 0.280 | 0.280 | -0.050 | 0.003 | -0.172 | 0.866 | 0.975 |
| Complexity | VIS | FPN | between | N2O | 0.267 | 0.267 | 0.002 | 0.005 | 0.009 | 0.993 | 0.993 |
| Complexity | VIS | FPN | between | Sleep N1 | 0.261 | 0.261 | -0.032 | 0.001 | -0.184 | 0.855 | 0.975 |
| Complexity | VIS | FPN | between | Sleep N2 | 0.261 | 0.256 | -0.525 | 0.002 | -2.829 | 0.009 | 0.038 |
| Complexity | VIS | FPN | between | PPF1.9 | 0.268 | 0.260 | -0.654 | 0.003 | -2.446 | 0.029 | 0.066 |
| Complexity | VIS | FPN | between | PPF2.4 | 0.160 | 0.151 | -0.887 | 0.002 | -4.524 | 0.000 | 0.001 |
| Complexity | VIS | FPN | between | PPF2.7 | 0.211 | 0.205 | -0.420 | 0.002 | -2.181 | 0.038 | 0.069 |
| Complexity | VIS | DMN | between | LSD | 0.218 | 0.224 | 0.523 | 0.003 | 2.025 | 0.062 | 0.112 |
| Complexity | VIS | DMN | between | PSIL | 0.241 | 0.250 | 0.552 | 0.006 | 1.460 | 0.194 | 0.290 |
| Complexity | VIS | DMN | between | KTM | 0.271 | 0.271 | -0.067 | 0.004 | -0.234 | 0.819 | 0.819 |
| Complexity | VIS | DMN | between | N2O | 0.262 | 0.257 | -0.327 | 0.004 | -1.268 | 0.226 | 0.290 |
| Complexity | VIS | DMN | between | Sleep N1 | 0.255 | 0.254 | -0.124 | 0.001 | -0.711 | 0.482 | 0.543 |
| Complexity | VIS | DMN | between | Sleep N2 | 0.255 | 0.246 | -0.897 | 0.002 | -4.829 | 0.000 | 0.000 |
| Complexity | VIS | DMN | between | PPF1.9 | 0.259 | 0.249 | -1.136 | 0.002 | -4.251 | 0.001 | 0.002 |
| Complexity | VIS | DMN | between | PPF2.4 | 0.155 | 0.144 | -1.011 | 0.002 | -5.155 | 0.000 | 0.000 |
| Complexity | VIS | DMN | between | PPF2.7 | 0.205 | 0.197 | -0.763 | 0.002 | -3.966 | 0.001 | 0.002 |
| Complexity | SMN | SMN | within | LSD | 0.236 | 0.236 | 0.037 | 0.003 | 0.145 | 0.887 | 0.887 |
| Complexity | SMN | SMN | within | PSIL | 0.242 | 0.244 | 0.276 | 0.003 | 0.731 | 0.492 | 0.686 |
| Complexity | SMN | SMN | within | KTM | 0.281 | 0.282 | 0.114 | 0.002 | 0.393 | 0.702 | 0.789 |
| Complexity | SMN | SMN | within | N2O | 0.261 | 0.259 | -0.243 | 0.003 | -0.942 | 0.362 | 0.652 |
| Complexity | SMN | SMN | within | Sleep N1 | 0.258 | 0.257 | -0.110 | 0.001 | -0.630 | 0.533 | 0.686 |
| Complexity | SMN | SMN | within | Sleep N2 | 0.257 | 0.252 | -0.433 | 0.002 | -2.334 | 0.027 | 0.061 |
| Complexity | SMN | SMN | within | PPF1.9 | 0.264 | 0.253 | -0.883 | 0.003 | -3.303 | 0.006 | 0.017 |
| Complexity | SMN | SMN | within | PPF2.4 | 0.172 | 0.158 | -1.577 | 0.002 | -8.040 | 0.000 | 0.000 |
| Complexity | SMN | SMN | within | PPF2.7 | 0.217 | 0.204 | -1.161 | 0.002 | -6.033 | 0.000 | 0.000 |
| Complexity | SMN | DAN | between | LSD | 0.241 | 0.250 | 0.506 | 0.004 | 1.960 | 0.070 | 0.126 |
| Complexity | SMN | DAN | between | PSIL | 0.262 | 0.269 | 0.610 | 0.005 | 1.613 | 0.158 | 0.203 |
| Complexity | SMN | DAN | between | KTM | 0.302 | 0.307 | 0.454 | 0.003 | 1.571 | 0.144 | 0.203 |
| Complexity | SMN | DAN | between | N2O | 0.283 | 0.286 | 0.187 | 0.004 | 0.723 | 0.482 | 0.482 |
| Complexity | SMN | DAN | between | Sleep N1 | 0.281 | 0.278 | -0.229 | 0.002 | -1.313 | 0.199 | 0.223 |
| Complexity | SMN | DAN | between | Sleep N2 | 0.281 | 0.271 | -0.608 | 0.003 | -3.276 | 0.003 | 0.006 |
| Complexity | SMN | DAN | between | PPF1.9 | 0.283 | 0.267 | -1.085 | 0.004 | -4.059 | 0.001 | 0.004 |
| Complexity | SMN | DAN | between | PPF2.4 | 0.175 | 0.159 | -1.436 | 0.002 | -7.320 | 0.000 | 0.000 |
| Complexity | SMN | DAN | between | PPF2.7 | 0.229 | 0.212 | -1.092 | 0.003 | -5.674 | 0.000 | 0.000 |
| Complexity | SMN | VAN | between | LSD | 0.247 | 0.250 | 0.199 | 0.003 | 0.769 | 0.455 | 0.585 |
| Complexity | SMN | VAN | between | PSIL | 0.263 | 0.270 | 0.556 | 0.005 | 1.470 | 0.192 | 0.288 |
| Complexity | SMN | VAN | between | KTM | 0.304 | 0.309 | 0.473 | 0.003 | 1.640 | 0.129 | 0.233 |
| Complexity | SMN | VAN | between | N2O | 0.287 | 0.289 | 0.118 | 0.003 | 0.457 | 0.655 | 0.660 |
| Complexity | SMN | VAN | between | Sleep N1 | 0.282 | 0.281 | -0.077 | 0.002 | -0.444 | 0.660 | 0.660 |
| Complexity | SMN | VAN | between | Sleep N2 | 0.282 | 0.275 | -0.610 | 0.002 | -3.284 | 0.003 | 0.008 |
| Complexity | SMN | VAN | between | PPF1.9 | 0.287 | 0.276 | -0.662 | 0.004 | -2.476 | 0.028 | 0.063 |
| Complexity | SMN | VAN | between | PPF2.4 | 0.175 | 0.162 | -1.186 | 0.002 | -6.047 | 0.000 | 0.000 |
| Complexity | SMN | VAN | between | PPF2.7 | 0.229 | 0.215 | -1.254 | 0.002 | -6.516 | 0.000 | 0.000 |
| Complexity | SMN | LIM | between | LSD | 0.280 | 0.282 | 0.117 | 0.004 | 0.453 | 0.658 | 0.920 |
| Complexity | SMN | LIM | between | PSIL | 0.310 | 0.309 | -0.040 | 0.004 | -0.105 | 0.920 | 0.920 |
| Complexity | SMN | LIM | between | KTM | 0.348 | 0.348 | -0.032 | 0.002 | -0.112 | 0.913 | 0.920 |
| Complexity | SMN | LIM | between | N2O | 0.320 | 0.315 | -0.243 | 0.005 | -0.941 | 0.363 | 0.653 |
| Complexity | SMN | LIM | between | Sleep N1 | 0.322 | 0.322 | 0.032 | 0.002 | 0.182 | 0.857 | 0.920 |
| Complexity | SMN | LIM | between | Sleep N2 | 0.322 | 0.319 | -0.210 | 0.003 | -1.132 | 0.267 | 0.602 |
| Complexity | SMN | LIM | between | PPF1.9 | 0.334 | 0.314 | -0.764 | 0.007 | -2.859 | 0.013 | 0.060 |
| Complexity | SMN | LIM | between | PPF2.4 | 0.199 | 0.183 | -1.392 | 0.002 | -7.096 | 0.000 | 0.000 |
| Complexity | SMN | LIM | between | PPF2.7 | 0.255 | 0.247 | -0.426 | 0.003 | -2.214 | 0.036 | 0.108 |
| Complexity | SMN | FPN | between | LSD | 0.226 | 0.234 | 0.679 | 0.003 | 2.630 | 0.020 | 0.036 |
| Complexity | SMN | FPN | between | PSIL | 0.253 | 0.260 | 0.549 | 0.005 | 1.453 | 0.196 | 0.253 |
| Complexity | SMN | FPN | between | KTM | 0.285 | 0.290 | 0.445 | 0.003 | 1.541 | 0.152 | 0.227 |
| Complexity | SMN | FPN | between | N2O | 0.270 | 0.273 | 0.272 | 0.004 | 1.055 | 0.309 | 0.348 |
| Complexity | SMN | FPN | between | Sleep N1 | 0.266 | 0.266 | -0.022 | 0.002 | -0.127 | 0.900 | 0.900 |
| Complexity | SMN | FPN | between | Sleep N2 | 0.266 | 0.260 | -0.538 | 0.002 | -2.896 | 0.007 | 0.022 |
| Complexity | SMN | FPN | between | PPF1.9 | 0.271 | 0.260 | -0.726 | 0.004 | -2.716 | 0.018 | 0.036 |
| Complexity | SMN | FPN | between | PPF2.4 | 0.161 | 0.151 | -0.903 | 0.002 | -4.605 | 0.000 | 0.001 |
| Complexity | SMN | FPN | between | PPF2.7 | 0.211 | 0.204 | -0.601 | 0.002 | -3.122 | 0.004 | 0.020 |
| Complexity | SMN | DMN | between | LSD | 0.208 | 0.211 | 0.279 | 0.003 | 1.081 | 0.298 | 0.383 |
| Complexity | SMN | DMN | between | PSIL | 0.230 | 0.237 | 0.506 | 0.005 | 1.340 | 0.229 | 0.343 |
| Complexity | SMN | DMN | between | KTM | 0.260 | 0.264 | 0.375 | 0.003 | 1.297 | 0.221 | 0.343 |
| Complexity | SMN | DMN | between | N2O | 0.245 | 0.247 | 0.114 | 0.004 | 0.442 | 0.665 | 0.748 |
| Complexity | SMN | DMN | between | Sleep N1 | 0.243 | 0.243 | -0.026 | 0.002 | -0.148 | 0.883 | 0.883 |
| Complexity | SMN | DMN | between | Sleep N2 | 0.244 | 0.237 | -0.646 | 0.002 | -3.481 | 0.002 | 0.005 |
| Complexity | SMN | DMN | between | PPF1.9 | 0.246 | 0.236 | -0.771 | 0.003 | -2.883 | 0.013 | 0.029 |
| Complexity | SMN | DMN | between | PPF2.4 | 0.147 | 0.137 | -0.986 | 0.002 | -5.029 | 0.000 | 0.000 |
| Complexity | SMN | DMN | between | PPF2.7 | 0.193 | 0.185 | -0.806 | 0.002 | -4.188 | 0.000 | 0.001 |
| Complexity | DAN | DAN | within | LSD | 0.255 | 0.261 | 0.543 | 0.003 | 2.104 | 0.054 | 0.097 |
| Complexity | DAN | DAN | within | PSIL | 0.269 | 0.273 | 0.375 | 0.004 | 0.992 | 0.359 | 0.539 |
| Complexity | DAN | DAN | within | KTM | 0.309 | 0.310 | 0.151 | 0.003 | 0.524 | 0.610 | 0.687 |
| Complexity | DAN | DAN | within | N2O | 0.293 | 0.291 | -0.104 | 0.004 | -0.404 | 0.692 | 0.692 |
| Complexity | DAN | DAN | within | Sleep N1 | 0.290 | 0.289 | -0.109 | 0.001 | -0.627 | 0.535 | 0.687 |
| Complexity | DAN | DAN | within | Sleep N2 | 0.291 | 0.285 | -0.445 | 0.002 | -2.399 | 0.023 | 0.053 |
| Complexity | DAN | DAN | within | PPF1.9 | 0.292 | 0.277 | -1.399 | 0.003 | -5.236 | 0.000 | 0.000 |
| Complexity | DAN | DAN | within | PPF2.4 | 0.187 | 0.172 | -1.415 | 0.002 | -7.214 | 0.000 | 0.000 |
| Complexity | DAN | DAN | within | PPF2.7 | 0.241 | 0.230 | -0.881 | 0.002 | -4.577 | 0.000 | 0.000 |
| Complexity | DAN | VAN | between | LSD | 0.242 | 0.247 | 0.396 | 0.003 | 1.534 | 0.147 | 0.201 |
| Complexity | DAN | VAN | between | PSIL | 0.261 | 0.269 | 0.518 | 0.005 | 1.371 | 0.220 | 0.247 |
| Complexity | DAN | VAN | between | KTM | 0.300 | 0.304 | 0.440 | 0.003 | 1.523 | 0.156 | 0.201 |
| Complexity | DAN | VAN | between | N2O | 0.286 | 0.287 | 0.065 | 0.004 | 0.250 | 0.806 | 0.806 |
| Complexity | DAN | VAN | between | Sleep N1 | 0.282 | 0.279 | -0.314 | 0.001 | -1.805 | 0.081 | 0.145 |
| Complexity | DAN | VAN | between | Sleep N2 | 0.282 | 0.273 | -0.748 | 0.002 | -4.030 | 0.000 | 0.001 |
| Complexity | DAN | VAN | between | PPF1.9 | 0.285 | 0.271 | -0.988 | 0.004 | -3.696 | 0.003 | 0.006 |
| Complexity | DAN | VAN | between | PPF2.4 | 0.172 | 0.158 | -1.252 | 0.002 | -6.382 | 0.000 | 0.000 |
| Complexity | DAN | VAN | between | PPF2.7 | 0.228 | 0.213 | -1.364 | 0.002 | -7.090 | 0.000 | 0.000 |
| Complexity | DAN | LIM | between | LSD | 0.276 | 0.281 | 0.308 | 0.003 | 1.194 | 0.252 | 0.568 |
| Complexity | DAN | LIM | between | PSIL | 0.304 | 0.306 | 0.113 | 0.006 | 0.299 | 0.775 | 0.775 |
| Complexity | DAN | LIM | between | KTM | 0.345 | 0.344 | -0.103 | 0.002 | -0.358 | 0.727 | 0.775 |
| Complexity | DAN | LIM | between | N2O | 0.320 | 0.315 | -0.208 | 0.005 | -0.804 | 0.435 | 0.652 |
| Complexity | DAN | LIM | between | Sleep N1 | 0.325 | 0.326 | 0.097 | 0.002 | 0.556 | 0.582 | 0.749 |
| Complexity | DAN | LIM | between | Sleep N2 | 0.325 | 0.323 | -0.169 | 0.002 | -0.912 | 0.370 | 0.652 |
| Complexity | DAN | LIM | between | PPF1.9 | 0.331 | 0.311 | -0.806 | 0.007 | -3.016 | 0.010 | 0.030 |
| Complexity | DAN | LIM | between | PPF2.4 | 0.197 | 0.181 | -1.504 | 0.002 | -7.669 | 0.000 | 0.000 |
| Complexity | DAN | LIM | between | PPF2.7 | 0.259 | 0.248 | -0.771 | 0.003 | -4.007 | 0.000 | 0.002 |
| Complexity | DAN | FPN | between | LSD | 0.230 | 0.239 | 0.705 | 0.003 | 2.731 | 0.016 | 0.037 |
| Complexity | DAN | FPN | between | PSIL | 0.261 | 0.266 | 0.335 | 0.006 | 0.888 | 0.409 | 0.526 |
| Complexity | DAN | FPN | between | KTM | 0.294 | 0.298 | 0.389 | 0.003 | 1.347 | 0.205 | 0.308 |
| Complexity | DAN | FPN | between | N2O | 0.282 | 0.283 | 0.071 | 0.004 | 0.274 | 0.788 | 0.788 |
| Complexity | DAN | FPN | between | Sleep N1 | 0.279 | 0.279 | 0.108 | 0.001 | 0.623 | 0.538 | 0.605 |
| Complexity | DAN | FPN | between | Sleep N2 | 0.279 | 0.275 | -0.393 | 0.002 | -2.116 | 0.043 | 0.078 |
| Complexity | DAN | FPN | between | PPF1.9 | 0.280 | 0.269 | -0.810 | 0.004 | -3.031 | 0.010 | 0.029 |
| Complexity | DAN | FPN | between | PPF2.4 | 0.168 | 0.159 | -0.898 | 0.002 | -4.579 | 0.000 | 0.001 |
| Complexity | DAN | FPN | between | PPF2.7 | 0.224 | 0.215 | -0.728 | 0.002 | -3.784 | 0.001 | 0.004 |
| Complexity | DAN | DMN | between | LSD | 0.233 | 0.237 | 0.376 | 0.003 | 1.458 | 0.167 | 0.256 |
| Complexity | DAN | DMN | between | PSIL | 0.255 | 0.266 | 0.580 | 0.007 | 1.535 | 0.176 | 0.256 |
| Complexity | DAN | DMN | between | KTM | 0.291 | 0.295 | 0.394 | 0.004 | 1.367 | 0.199 | 0.256 |
| Complexity | DAN | DMN | between | N2O | 0.278 | 0.279 | 0.085 | 0.004 | 0.329 | 0.747 | 0.760 |
| Complexity | DAN | DMN | between | Sleep N1 | 0.276 | 0.277 | 0.054 | 0.002 | 0.308 | 0.760 | 0.760 |
| Complexity | DAN | DMN | between | Sleep N2 | 0.276 | 0.269 | -0.535 | 0.003 | -2.883 | 0.007 | 0.017 |
| Complexity | DAN | DMN | between | PPF1.9 | 0.276 | 0.263 | -0.927 | 0.004 | -3.470 | 0.004 | 0.012 |
| Complexity | DAN | DMN | between | PPF2.4 | 0.164 | 0.154 | -1.089 | 0.002 | -5.552 | 0.000 | 0.000 |
| Complexity | DAN | DMN | between | PPF2.7 | 0.221 | 0.210 | -0.937 | 0.002 | -4.868 | 0.000 | 0.000 |
| Complexity | VAN | VAN | within | LSD | 0.262 | 0.262 | 0.032 | 0.003 | 0.122 | 0.904 | 0.904 |
| Complexity | VAN | VAN | within | PSIL | 0.269 | 0.273 | 0.375 | 0.004 | 0.993 | 0.359 | 0.646 |
| Complexity | VAN | VAN | within | KTM | 0.312 | 0.312 | 0.074 | 0.002 | 0.256 | 0.803 | 0.903 |
| Complexity | VAN | VAN | within | N2O | 0.296 | 0.294 | -0.104 | 0.005 | -0.404 | 0.692 | 0.890 |
| Complexity | VAN | VAN | within | Sleep N1 | 0.293 | 0.292 | -0.136 | 0.001 | -0.781 | 0.441 | 0.661 |
| Complexity | VAN | VAN | within | Sleep N2 | 0.293 | 0.287 | -0.464 | 0.002 | -2.499 | 0.019 | 0.042 |
| Complexity | VAN | VAN | within | PPF1.9 | 0.301 | 0.288 | -0.898 | 0.004 | -3.358 | 0.005 | 0.015 |
| Complexity | VAN | VAN | within | PPF2.4 | 0.186 | 0.172 | -1.275 | 0.002 | -6.503 | 0.000 | 0.000 |
| Complexity | VAN | VAN | within | PPF2.7 | 0.243 | 0.226 | -1.806 | 0.002 | -9.383 | 0.000 | 0.000 |
| Complexity | VAN | LIM | between | LSD | 0.277 | 0.280 | 0.250 | 0.002 | 0.969 | 0.349 | 0.523 |
| Complexity | VAN | LIM | between | PSIL | 0.305 | 0.307 | 0.133 | 0.005 | 0.353 | 0.737 | 0.829 |
| Complexity | VAN | LIM | between | KTM | 0.349 | 0.345 | -0.434 | 0.002 | -1.502 | 0.161 | 0.363 |
| Complexity | VAN | LIM | between | N2O | 0.319 | 0.316 | -0.155 | 0.005 | -0.600 | 0.558 | 0.718 |
| Complexity | VAN | LIM | between | Sleep N1 | 0.326 | 0.326 | -0.001 | 0.002 | -0.008 | 0.994 | 0.994 |
| Complexity | VAN | LIM | between | Sleep N2 | 0.326 | 0.323 | -0.231 | 0.002 | -1.243 | 0.224 | 0.403 |
| Complexity | VAN | LIM | between | PPF1.9 | 0.339 | 0.321 | -0.699 | 0.007 | -2.617 | 0.021 | 0.064 |
| Complexity | VAN | LIM | between | PPF2.4 | 0.196 | 0.182 | -1.230 | 0.002 | -6.272 | 0.000 | 0.000 |
| Complexity | VAN | LIM | between | PPF2.7 | 0.261 | 0.250 | -0.813 | 0.003 | -4.223 | 0.000 | 0.001 |
| Complexity | VAN | FPN | between | LSD | 0.232 | 0.237 | 0.506 | 0.003 | 1.961 | 0.070 | 0.126 |
| Complexity | VAN | FPN | between | PSIL | 0.258 | 0.264 | 0.348 | 0.007 | 0.922 | 0.392 | 0.504 |
| Complexity | VAN | FPN | between | KTM | 0.294 | 0.300 | 0.521 | 0.003 | 1.805 | 0.099 | 0.148 |
| Complexity | VAN | FPN | between | N2O | 0.282 | 0.284 | 0.100 | 0.004 | 0.386 | 0.705 | 0.794 |
| Complexity | VAN | FPN | between | Sleep N1 | 0.279 | 0.279 | 0.033 | 0.001 | 0.190 | 0.850 | 0.850 |
| Complexity | VAN | FPN | between | Sleep N2 | 0.279 | 0.272 | -0.629 | 0.002 | -3.388 | 0.002 | 0.006 |
| Complexity | VAN | FPN | between | PPF1.9 | 0.285 | 0.275 | -0.726 | 0.004 | -2.718 | 0.018 | 0.040 |
| Complexity | VAN | FPN | between | PPF2.4 | 0.167 | 0.157 | -0.978 | 0.002 | -4.986 | 0.000 | 0.000 |
| Complexity | VAN | FPN | between | PPF2.7 | 0.222 | 0.211 | -1.103 | 0.002 | -5.729 | 0.000 | 0.000 |
| Complexity | VAN | DMN | between | LSD | 0.231 | 0.236 | 0.522 | 0.003 | 2.023 | 0.063 | 0.113 |
| Complexity | VAN | DMN | between | PSIL | 0.255 | 0.265 | 0.541 | 0.007 | 1.430 | 0.203 | 0.260 |
| Complexity | VAN | DMN | between | KTM | 0.292 | 0.297 | 0.539 | 0.003 | 1.866 | 0.089 | 0.133 |
| Complexity | VAN | DMN | between | N2O | 0.276 | 0.278 | 0.123 | 0.005 | 0.475 | 0.642 | 0.661 |
| Complexity | VAN | DMN | between | Sleep N1 | 0.276 | 0.277 | 0.077 | 0.001 | 0.443 | 0.661 | 0.661 |
| Complexity | VAN | DMN | between | Sleep N2 | 0.277 | 0.270 | -0.597 | 0.002 | -3.217 | 0.003 | 0.010 |
| Complexity | VAN | DMN | between | PPF1.9 | 0.283 | 0.275 | -0.556 | 0.004 | -2.080 | 0.058 | 0.113 |
| Complexity | VAN | DMN | between | PPF2.4 | 0.165 | 0.155 | -0.938 | 0.002 | -4.782 | 0.000 | 0.000 |
| Complexity | VAN | DMN | between | PPF2.7 | 0.221 | 0.210 | -1.134 | 0.002 | -5.895 | 0.000 | 0.000 |
| Complexity | LIM | LIM | within | LSD | 0.298 | 0.298 | -0.007 | 0.005 | -0.028 | 0.978 | 0.978 |
| Complexity | LIM | LIM | within | PSIL | 0.325 | 0.327 | 0.241 | 0.003 | 0.637 | 0.548 | 0.616 |
| Complexity | LIM | LIM | within | KTM | 0.358 | 0.353 | -0.307 | 0.004 | -1.062 | 0.311 | 0.591 |
| Complexity | LIM | LIM | within | N2O | 0.343 | 0.339 | -0.261 | 0.004 | -1.013 | 0.328 | 0.591 |
| Complexity | LIM | LIM | within | Sleep N1 | 0.340 | 0.342 | 0.133 | 0.002 | 0.763 | 0.451 | 0.616 |
| Complexity | LIM | LIM | within | Sleep N2 | 0.340 | 0.341 | 0.131 | 0.003 | 0.703 | 0.488 | 0.616 |
| Complexity | LIM | LIM | within | PPF1.9 | 0.350 | 0.338 | -0.407 | 0.008 | -1.525 | 0.151 | 0.454 |
| Complexity | LIM | LIM | within | PPF2.4 | 0.213 | 0.200 | -1.089 | 0.002 | -5.555 | 0.000 | 0.000 |
| Complexity | LIM | LIM | within | PPF2.7 | 0.277 | 0.267 | -0.627 | 0.003 | -3.259 | 0.003 | 0.014 |
| Complexity | LIM | FPN | between | LSD | 0.273 | 0.278 | 0.293 | 0.004 | 1.136 | 0.275 | 0.495 |
| Complexity | LIM | FPN | between | PSIL | 0.307 | 0.310 | 0.232 | 0.005 | 0.613 | 0.562 | 0.651 |
| Complexity | LIM | FPN | between | KTM | 0.342 | 0.343 | 0.126 | 0.004 | 0.437 | 0.671 | 0.671 |
| Complexity | LIM | FPN | between | N2O | 0.324 | 0.319 | -0.303 | 0.004 | -1.175 | 0.260 | 0.495 |
| Complexity | LIM | FPN | between | Sleep N1 | 0.327 | 0.328 | 0.098 | 0.002 | 0.561 | 0.578 | 0.651 |
| Complexity | LIM | FPN | between | Sleep N2 | 0.327 | 0.326 | -0.107 | 0.002 | -0.579 | 0.567 | 0.651 |
| Complexity | LIM | FPN | between | PPF1.9 | 0.339 | 0.320 | -0.669 | 0.007 | -2.502 | 0.026 | 0.079 |
| Complexity | LIM | FPN | between | PPF2.4 | 0.194 | 0.181 | -1.399 | 0.002 | -7.134 | 0.000 | 0.000 |
| Complexity | LIM | FPN | between | PPF2.7 | 0.262 | 0.250 | -0.876 | 0.002 | -4.553 | 0.000 | 0.000 |
| Complexity | LIM | DMN | between | LSD | 0.278 | 0.284 | 0.342 | 0.005 | 1.325 | 0.207 | 0.310 |
| Complexity | LIM | DMN | between | PSIL | 0.313 | 0.316 | 0.225 | 0.004 | 0.595 | 0.574 | 0.590 |
| Complexity | LIM | DMN | between | KTM | 0.343 | 0.345 | 0.160 | 0.004 | 0.555 | 0.590 | 0.590 |
| Complexity | LIM | DMN | between | N2O | 0.330 | 0.319 | -0.539 | 0.005 | -2.086 | 0.056 | 0.126 |
| Complexity | LIM | DMN | between | Sleep N1 | 0.330 | 0.331 | 0.100 | 0.002 | 0.572 | 0.571 | 0.590 |
| Complexity | LIM | DMN | between | Sleep N2 | 0.330 | 0.326 | -0.260 | 0.002 | -1.399 | 0.173 | 0.310 |
| Complexity | LIM | DMN | between | PPF1.9 | 0.341 | 0.320 | -0.775 | 0.007 | -2.901 | 0.012 | 0.037 |
| Complexity | LIM | DMN | between | PPF2.4 | 0.199 | 0.182 | -1.646 | 0.002 | -8.394 | 0.000 | 0.000 |
| Complexity | LIM | DMN | between | PPF2.7 | 0.265 | 0.252 | -0.887 | 0.003 | -4.607 | 0.000 | 0.000 |
| Complexity | FPN | FPN | within | LSD | 0.239 | 0.247 | 0.666 | 0.003 | 2.580 | 0.022 | 0.039 |
| Complexity | FPN | FPN | within | PSIL | 0.260 | 0.264 | 0.328 | 0.005 | 0.867 | 0.419 | 0.626 |
| Complexity | FPN | FPN | within | KTM | 0.296 | 0.298 | 0.208 | 0.002 | 0.720 | 0.487 | 0.626 |
| Complexity | FPN | FPN | within | N2O | 0.285 | 0.283 | -0.100 | 0.004 | -0.387 | 0.704 | 0.756 |
| Complexity | FPN | FPN | within | Sleep N1 | 0.282 | 0.283 | 0.055 | 0.002 | 0.313 | 0.756 | 0.756 |
| Complexity | FPN | FPN | within | Sleep N2 | 0.282 | 0.278 | -0.454 | 0.002 | -2.447 | 0.021 | 0.039 |
| Complexity | FPN | FPN | within | PPF1.9 | 0.287 | 0.276 | -0.777 | 0.004 | -2.907 | 0.012 | 0.037 |
| Complexity | FPN | FPN | within | PPF2.4 | 0.174 | 0.165 | -0.989 | 0.002 | -5.043 | 0.000 | 0.000 |
| Complexity | FPN | FPN | within | PPF2.7 | 0.229 | 0.221 | -0.763 | 0.002 | -3.965 | 0.001 | 0.002 |
| Complexity | FPN | DMN | between | LSD | 0.221 | 0.231 | 0.837 | 0.003 | 3.241 | 0.006 | 0.013 |
| Complexity | FPN | DMN | between | PSIL | 0.251 | 0.261 | 0.578 | 0.007 | 1.530 | 0.177 | 0.227 |
| Complexity | FPN | DMN | between | KTM | 0.287 | 0.293 | 0.505 | 0.003 | 1.748 | 0.108 | 0.162 |
| Complexity | FPN | DMN | between | N2O | 0.275 | 0.278 | 0.125 | 0.005 | 0.484 | 0.636 | 0.715 |
| Complexity | FPN | DMN | between | Sleep N1 | 0.274 | 0.274 | 0.023 | 0.002 | 0.132 | 0.896 | 0.896 |
| Complexity | FPN | DMN | between | Sleep N2 | 0.274 | 0.265 | -0.752 | 0.002 | -4.052 | 0.000 | 0.001 |
| Complexity | FPN | DMN | between | PPF1.9 | 0.276 | 0.266 | -0.734 | 0.004 | -2.747 | 0.017 | 0.030 |
| Complexity | FPN | DMN | between | PPF2.4 | 0.162 | 0.153 | -0.879 | 0.002 | -4.480 | 0.000 | 0.001 |
| Complexity | FPN | DMN | between | PPF2.7 | 0.218 | 0.208 | -0.913 | 0.002 | -4.744 | 0.000 | 0.001 |
| Complexity | DMN | DMN | within | LSD | 0.210 | 0.216 | 0.567 | 0.003 | 2.196 | 0.045 | 0.082 |
| Complexity | DMN | DMN | within | PSIL | 0.230 | 0.234 | 0.388 | 0.004 | 1.025 | 0.345 | 0.443 |
| Complexity | DMN | DMN | within | KTM | 0.260 | 0.263 | 0.253 | 0.003 | 0.875 | 0.400 | 0.450 |
| Complexity | DMN | DMN | within | N2O | 0.253 | 0.249 | -0.296 | 0.004 | -1.148 | 0.270 | 0.405 |
| Complexity | DMN | DMN | within | Sleep N1 | 0.249 | 0.248 | -0.077 | 0.001 | -0.444 | 0.660 | 0.660 |
| Complexity | DMN | DMN | within | Sleep N2 | 0.249 | 0.241 | -0.724 | 0.002 | -3.897 | 0.001 | 0.002 |
| Complexity | DMN | DMN | within | PPF1.9 | 0.252 | 0.241 | -0.880 | 0.003 | -3.293 | 0.006 | 0.013 |
| Complexity | DMN | DMN | within | PPF2.4 | 0.153 | 0.143 | -1.238 | 0.002 | -6.315 | 0.000 | 0.000 |
| Complexity | DMN | DMN | within | PPF2.7 | 0.203 | 0.194 | -1.106 | 0.002 | -5.746 | 0.000 | 0.000 |

**Abbreviations:** FC: functional connectivity; LSD: lysergic acid diethylamide; PSIL: psilocybin; KTM: ketamine; N<sub>2</sub>O: nitrous oxide; PPF1.9 / PPF2.4 / PPF2.7: propofol (effect-site concentrations: 1.9 / 2.4 / 2.7 µg·mL<sup>-1</sup>). U-U: unimodal-unimodal; A-A: attention-attention; T-T: transmodal-transmodal; U-A: unimodal-attention; U-T: unimodal-transmodal; A-T: attention-transmodal. VIS: visual network; SMN: somatomotor network; DAN: dorsal attention network; VAN: ventral attention network; LIM: limbic network; FPN: frontoparietal network; DMN: default-mode network.

## References

1. Chalmers, D. J. Facing Up to the Problem of Consciousness. J. Conscious. Stud. 2, 200–219 (1995).

2. Crick, F. & Koch, C. Towards a neurobiological theory of consciousness. Semin Neurosci 1990; 2: 263–75. *Acad. Press* **2**, 263–275 (1990).

3. Koch, C., Massimini, M., Boly, M. & Tononi, G. Neural correlates of consciousness: progress and problems. Nat. Rev. Neurosci. 17, 307–21 (2016).

4. Huang, Z., Liu, X., Mashour, G. A. & Hudetz, A. G. Timescales of Intrinsic BOLD Signal Dynamics and Functional Connectivity in Pharmacologic and Neuropathologic States of Unconsciousness. J. Neurosci. 38, 2304–2317 (2018).

5. Mashour, G. A. Anesthesia and the neurobiology of consciousness. Neuron 112, 1553–1567 (2024).

6. Carhart-Harris, R. L. et al. Neural correlates of the LSD experience revealed by multimodal neuroimaging. Proc. Natl. Acad. Sci. U. S. A. 113, 4853–8 (2016).

7. Erritzoe, D. et al. Exploring mechanisms of psychedelic action using neuroimaging. Nat. Ment. Heal. 2, 141–153 (2024).

8. Dai, R. et al. Classical and non-classical psychedelic drugs induce common network changes in human cortex. Neuroimage 273, 120097 (2023).

9. Boveroux, P. et al. Breakdown of within- and between-network resting state functional magnetic resonance imaging connectivity during propofol-induced loss of consciousness. Anesthesiology 113, 1038–1053 (2010).

10. Liu, X., Lauer, K. K., Ward, B. D., Li, S. J. & Hudetz, A. G. Differential effects of deep sedation with propofol on the specific and nonspecific thalamocortical systems: A functional magnetic resonance imaging study. Anesthesiology 118, 59–69 (2013).

11. Zelmann, R. et al. Differential cortical network engagement during states of un/consciousness in humans. Neuron 111, 3479–3495.e6 (2023).

12. Lee, U. & Mashour, G. A. Role of Network Science in the Study of Anesthetic State Transitions. Anesthesiology 129, 1029–1044 (2018).

13. Luppi, A. I. et al. Brain network integration dynamics are associated with loss and recovery of consciousness induced by sevoflurane. Hum. Brain Mapp. 42, 2802–2822 (2021).

14. Barttfeld, P. et al. Signature of consciousness in the dynamics of resting-state brain activity. Proc. Natl. Acad. Sci. U. S. A. 112, 887–92 (2015).

15. Luppi, A. I. et al. Distributed harmonic patterns of structure-function dependence orchestrate human consciousness. *Commun*. Biol. 6, 117 (2023).

16. Preller, K. H. et al. Changes in global and thalamic brain connectivity in LSD-induced altered states of consciousness are attributable to the 5-HT2A receptor. Elife 7, 1–31 (2018).

17. Timmermann, C. et al. Human brain effects of DMT assessed via EEG-fMRI. Proc. Natl. Acad. Sci. 120, 2017 (2023).

18. Girn, M. et al. Serotonergic psychedelic drugs LSD and psilocybin reduce the hierarchical differentiation of unimodal and transmodal cortex. Neuroimage 256, 119220 (2022).

19. Luppi, A. I. et al. LSD alters dynamic integration and segregation in the human brain. Neuroimage 227, 117653 (2021).

20. Dai, R. et al. Psychedelic concentrations of nitrous oxide reduce functional differentiation in frontoparietal and somatomotor cortical networks. *Commun*. Biol. 6, 1–10 (2023).

21. Atasoy, S., Vohryzek, J., Deco, G., Carhart-harris, R. L. & Kringelbach, M. L. Common neural signatures of psychedelics: Frequency-specific energy changes and repertoire expansion revealed using connectome-harmonic decomposition. Psychedelic Neuroscience vol. 242 (Elsevier B.V., 2018).

22. McCulloch, D. E.-W. et al. Psychedelic resting-state neuroimaging: A review and perspective on balancing replication and novel analyses. Neurosci. Biobehav. Rev. 138, 104689 (2022).

23. Singleton, S. P. et al. Receptor-informed network control theory links LSD and psilocybin to a flattening of the brain’s control energy landscape. Nat. Commun. 13, 1–13 (2022).

24. Jang, H., Mashour, G. A., Hudetz, A. G. & Huang, Z. Measuring the dynamic balance of integration and segregation underlying consciousness, anesthesia, and sleep in humans. Nat. Commun. 15, 9164 (2024).

25. Siegel, J. S. et al. Psilocybin desynchronizes the human brain. Nature 632, (2024).

26. Shinn, M. et al. Functional brain networks reflect spatial and temporal autocorrelation. Nat. Neurosci. 26, 867–878 (2023).

27. Horovitz, S. G. et al. Decoupling of the brain’s default mode network during deep sleep. Proc. Natl. Acad. Sci. U. S. A. 106, 11376–11381 (2009).

28. Tagliazucchi, E. & Laufs, H. Decoding Wakefulness Levels from Typical fMRI Resting-State Data Reveals Reliable Drifts between Wakefulness and Sleep. Neuron 82, 695–708 (2014).

29. Palanca, B. J. A. et al. Resting-state functional magnetic resonance imaging correlates of sevoflurane-induced unconsciousness. Anesthesiology 123, 346–356 (2015).

30. Bonhomme, V. et al. Resting-state Network-specific Breakdown of Functional Connectivity during Ketamine Alteration of Consciousness in Volunteers. Anesthesiology 125, 873–888 (2016).

31. Jordan, D. et al. Simultaneous electroencephalographic and functional magnetic resonance imaging indicate impaired cortical top-down processing in association with anesthetic-induced unconsciousness. Anesthesiology 119, 1031–42 (2013).

32. Ranft, A. et al. Neural Correlates of Sevoflurane-induced Unconsciousness Identified by Simultaneous Functional Magnetic Resonance Imaging and Electroencephalography. Anesthesiology 125, 861–872 (2016).

33. Margulies, D. S. et al. Situating the default-mode network along a principal gradient of macroscale cortical organization. Proc. Natl. Acad. Sci. U. S. A. 113, 12574–12579 (2016).

34. Hong, S. J. et al. Atypical functional connectome hierarchy in autism (Supplementary Information). Nat. Commun. 10, 1–13 (2019).

35. Bethlehem, R. A. I. et al. Dispersion of functional gradients across the adult lifespan. Neuroimage 222, 117299 (2020).

36. Cross, N. et al. Cortical gradients of functional connectivity are robust to state-dependent changes following sleep deprivation. Neuroimage 226, (2021).

37. Huang, Z., Mashour, G. A. & Hudetz, A. G. Functional geometry of the cortex encodes dimensions of consciousness. Nat. Commun. 14, 72 (2023).

38. Huang, Z., Mashour, G. A. & Hudetz, A. G. Propofol disrupts the functional core-matrix architecture of the thalamus in humans. Nat. Commun. 15, 1–13 (2024).

39. Tagliazucchi, E. et al. Increased Global Functional Connectivity Correlates with LSD-Induced Ego Dissolution. Curr. Biol. 26, 1043–1050 (2016).

40. Studerus, E., Gamma, A. & Vollenweider, F. X. Psychometric evaluation of the altered states of consciousness rating scale (OAV). PLoS One 5, e12412 (2010).

41. Huang, Z., Zhang, J., Wu, J., Mashour, G. A. & Hudetz, A. G. Temporal circuit of macroscale dynamic brain activity supports human consciousness. Sci. Adv. 6, 1–15 (2020).

42. Gu, Y., Sainburg, L. E., Han, F. & Liu, X. Simultaneous EEG and functional MRI data during rest and sleep from humans. Data Br. 48, 109059 (2023).

43. Gu, Y. et al. An orderly sequence of autonomic and neural events at transient arousal changes. Neuroimage 264, 119720 (2022).

44. Liu, X. et al. Propofol attenuates low-frequency fluctuations of resting-state fMRI BOLD signal in the anterior frontal cortex upon loss of consciousness. Neuroimage 147, 295–301 (2017).

45. Huang, Z. et al. Anterior insula regulates brain network transitions that gate conscious access. Cell Rep. 35, 109081 (2021).

46. Huang, Z. et al. Asymmetric neural dynamics characterize loss and recovery of consciousness. Neuroimage 236, 118042 (2021).

47. Talairach, J. & Tournoux, P. Co-Planar Stereotaxis Atlas of the Human Brain: : an approach to cerebral imaging. Stuttgart, New York, New York G. (1988).

48. Schaefer, A. et al. Local-Global Parcellation of the Human Cerebral Cortex from Intrinsic Functional Connectivity MRI. Cereb. Cortex 28, 3095–3114 (2018).

49. Yeo, B. T. T. et al. The organization of the human cerebral cortex estimated by intrinsic functional connectivity. J. Neurophysiol. 106, 1125–65 (2011).

50. Rubinov, M. & Sporns, O. Complex network measures of brain connectivity: Uses and interpretations. Neuroimage 52, 1059–1069 (2010).

51. Muldoon, S. F., Bridgeford, E. W. & Bassett, D. S. Small-World Propensity and Weighted Brain Networks. Sci. Rep. 6, 22057 (2016).

